# Sequence context and partner availability specify divergent cell fates from a single transcription factor

**DOI:** 10.64898/2026.11.09.758913

**Authors:** Hammad Farooq, Kenneth Joves, Aydolun Petenkaya, Itay Solomon, Eva H.W Leung, Sen Zhang, Mary Menhart, Ali Farhat, Lin Du, Nissim Hay, Sandra Pinho, Constantinos Chronis

## Abstract

Transcription factors drive the gene regulatory networks that control developmental transitions, yet which of their many targets a cell uses remains unresolved. Expressing Cebpα, Cebpβ or Cebpδ identically, alone or with Oct4, Sox2, Klf4 and cMyc, gave four outcomes: adipocyte from fibroblasts and neural stem cells, trophectoderm from embryonic stem cells, myeloid from B cells. Raising Cebp levels changed how many sites it reached, not which sequence it read: clustered motifs maintained somatic enhancers, while spillover onto low-copy sites created de novo enhancers carrying composites of Cebp and cell-type-restricted partners. In fibroblasts, Cebp also redistributed Oct4, Sox2 and Klf4 from early targets, blocking iPSC formation. Supplying the predicted partners rewrote the output: Spi1 redirected Cebpα into parallel adipogenic and myeloid programs, via a transient *Pparg*⁺CD45⁺ state using separate enhancers for each; TFAP2C abolished adipogenic conversion. Motif density and partner composites therefore assemble a factor’s instructive output in the receiving cell.

## Introduction

The same transcription factor (TF) can drive strikingly different outcomes across different cell types, and explaining this context dependence is central to understanding development and controlling cell fate. However, how a TF selects its genomic targets remains unresolved even in principle. Pioneer access to closed chromatin^1, 2^, TF dose^3, 4^, and cooperative binding^5–7^ have each been proposed as determinants, yet none explains why in a given cell only a small fraction of available sites are bound, or how the same factor activates different gene regulatory networks (GRNs) in different contexts. Instructive capacity therefore appears not to be exclusively intrinsic to the protein but assembled from the regulatory context in which it acts.

The Cebp family illustrates this directly. Cebpα instructs adipocyte^8–10^, myeloid^11, 12^, and hepatocyte^13, 14^ differentiation, drives granulocyte/macrophage progenitor formation in hematopoiesis^15^, and, alongside Cebpβ, is required for extraembryonic, placental development^16^. It converts B cells into macrophages by partnering with endogenous PU.1^17–19^, and when co-expressed with Oct4, Sox2, Klf4, cMyc (OSKM), or delivered as a brief preceding pulse, primes B cells for rapid and efficient reprogramming to induced pluripotent stem cells (iPSCs)^20, 21^. A single protein therefore activates distinct GRNs depending on where it is expressed, yet what licenses one program rather than another remains poorly understood.

Context dependence has been inferred almost entirely from single factors in single systems, where redundant family members, resident partners, and lineage-optimized media conditions ^22^ confound interpretations. The converse experiment, delivering one TF identically into multiple unrelated starting cells, has not been performed. Additionally, resistance to reprogramming is attributed to reinforcement of the resident somatic program^5, 23^, and OSKM, which reprograms a broad range of somatic cell types^24–26^, is assumed to override competing lineage factors. This has been tested only by co-expressing individual Yamanaka factors with somatic factors, where none dominates^22^. Since the four reprogram only as a cooperative unit^5, 27^, whether the intact quartet of TFs can be overridden by a lineage-instructive TF remains unknown. Is a TF’s instructive output intrinsic to the protein, or set by the partners present in each cell, and therefore predictable from them and reprogrammable by adding or removing them?

We addressed both by expressing Cebpα, Cebpβ or Cebpδ identically, alone or with OSKM, in mouse embryonic fibroblasts (MEFs), neural stem cells (NSCs), embryonic stem cells (ESCs) and B cells, profiling transcriptome, chromatin accessibility, histone modifications and Cebp occupancy in parallel, and training base-resolution models (ChromBPNet)^28^ to recover the motifs driving accessibility and the partners contributing at Cebp-bound sites. The same protein gave opposite fates: adipocyte in fibroblasts and neural stem cells, trophectoderm in embryonic stem cells, myeloid in B cells. In fibroblasts it did not reinforce the resident program but overwrote it, transdifferentiating OSKM-expressing cells into adipocytes and blocking iPSC formation, to our knowledge the first instance of one somatic-lineage TF defeating the intact OSKM cocktail. The fates tracked the partner grammar at de novo Cebp enhancers: NFI, NF-Y and AP-1/ATF in fibroblasts and neural stem cells, SPI1 in B cells, TFAP2C in embryonic stem cells. Supplying those partners was sufficient to install or abolish a fate, with SPI1 adding a myeloid output to fibroblasts and TFAP2C removing the adipogenic one. Single-cell transcriptome, surface-protein and chromatin profiling showed that these two outputs need not be exclusive: a transient Pparg⁺CD45⁺ hybrid ran the adipogenic and myeloid networks together, on separate elements within the same nucleus, preceding the terminal adipocyte and macrophage states.

## Results

### Cebpα, Cebpβ and Cebpδ instruct fibroblasts for rapid transdifferentiation into adipocyte-like cells

To determine whether Cebp factors exert a comparable pro-pluripotency role outside the B-cell lineage, we turned to MEFs, the cell type most widely used for OSKM reprogramming. Retroviral expression of Flag-tagged mCebpα or mCebpβ in C57BL/6 MEFs (E13.5), in the absence of any reprogramming factor, generated lipid-laden, Oil Red O (ORO)-positive adipocyte-like cells by day 10, whereas empty-vector cultures remained fibroblastic (Fig. 1a–c and Supplementary Fig. 1a), consistent with the capacity of ectopic Cebpα to impose adipogenesis on diverse fibroblast lines^8^.

**Fig. 1.**
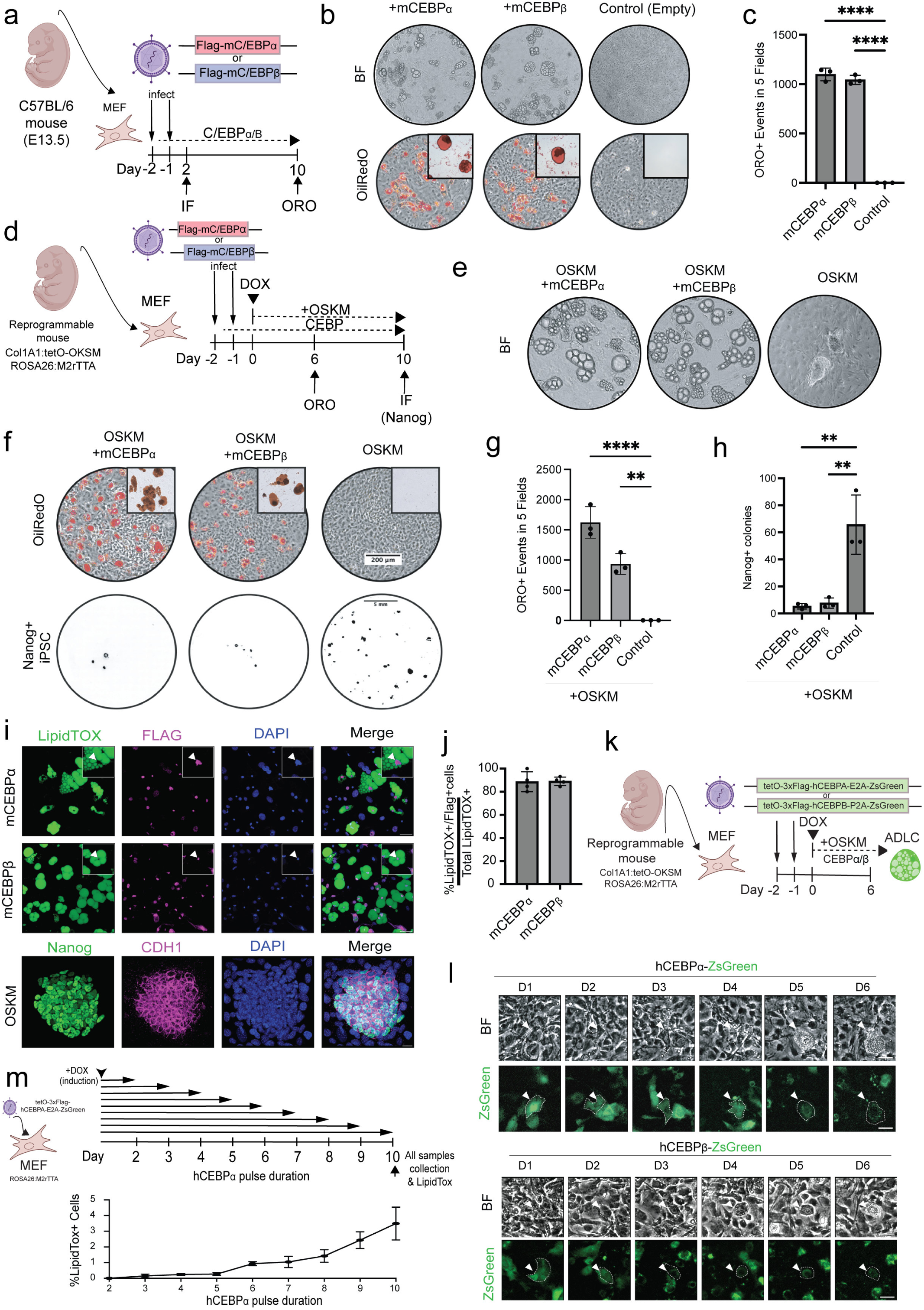
Cebp transcription factors promote the adipogenic fate and block MEF-to-iPSC conversion. **a**, Schematic of the Cebpα/β adipogenic-potential assay. MEFs (C57BL/6, E13.5) transduced with retroviral Flag-mCebpα, Flag-mCebpβ or empty vector and stained with Oil Red O (ORO) at day 10. **b**, Representative brightfield (BF; top) and ORO-stained (bottom; red) six-well plates at day 10. **c**, Quantification of ORO-positive events across five microscopic fields per experiment (mean ± s.d.; n = 3). \*\*\*\**P* < 0.0001, unpaired two-tailed Student’s t-test. **d**, Schematic of concurrent mCebp overexpression and OSKM (Oct4, Sox2, Klf4, cMyc)-mediated reprogramming. Reprogrammable MEFs (Col1a1:tetO-OSKM; ROSA26:M2rtTA) transduced with mCebp for 48 h, then induced with doxycycline (dox); ORO at day 6 and Nanog immunofluorescence (IF) at day 10. **e**, Representative brightfield images at day 10 of MEFs co-expressing OSKM with mCebpα or mCebpβ, or OSKM alone. **f**, Representative wells showing ORO staining at day 6 (top) and Nanog immunostaining at day 10 (bottom) in OSKM-only and OSKM + mCebpα/β MEFs; insets show high-magnification ORO-positive lipid droplets. Scale bar, 200 μm (ORO inset) and 5 mm (Nanog immunofluorescence, whole-well scan). **g**, Quantification of ORO-positive structures across five microscopic fields at day 6 per experiment (mean ± s.d.; n = 3). \*\*\*\**P* < 0.0001, \*\**P* < 0.01, unpaired two-tailed Student’s t-test. **h**, Quantification of Nanog-positive iPSC colonies at day 10 (mean ± s.d.; n = 3). \*\**P* < 0.01, unpaired two-tailed Student’s t-test. **i**, Immunofluorescence at day 6 of OSKM + Flag-mCebpα (top) or Flag-mCebpβ (middle) for LipidTOX (green), Flag (magenta) and DAPI (blue); OSKM-only stained for Nanog (green) and CDH1 (magenta) as a pluripotent reference (bottom). Scale bar, 50 μm. **j**, Proportion of LipidTOX-positive cells among total Flag-positive cells in OSKM + Flag-mCebpα and OSKM + Flag-mCebpβ cultures at day 6 per experiment (mean ± s.d.; n = 4). **k**, Live-cell tracking of adipocyte-like cell (ADLC) emergence: reprogrammable MEFs co-transduced with tetO-3×Flag-hCebpα-E2A-ZsGreen or tetO-3×Flag-hCebpβ-P2A-ZsGreen, OSKM induced at day 0. **l**, Sequential brightfield (top) and ZsGreen-fluorescence (bottom) images from day 1 to day 6 of MEFs co-expressing OSKM with hCebpα-ZsGreen (upper) or hCebpβ-ZsGreen (lower). Arrowheads, cells acquiring ADLC morphology coincident with ZsGreen. Scale bar, 50 μm. **m**, Top, dox added at day 0 to tetO-3×Flag-hCebpα MEFs (ROSA26:M2rtTA) and withdrawn at days 2-9; continuous induction as control; all conditions collected at day 10. Bottom, percentage of LipidTOX-positive cells at day 10 across the dox-induction intervals.

To determine how this activity intersects with reprogramming, we co-expressed mCebpα or mCebpβ with the polycistronic OSKM cassette in reprogrammable MEFs (Col1a1:tetO-OSKM; ROSA26:M2rtTA) (Fig. 1d). The two programs were mutually exclusive. Cebp co-expressing cultures adopted adipocyte-like morphology (Fig. 1e) and accumulated ORO-positive lipid droplets by day 6 (Fig. 1f,g), whereas OSKM-alone cultures formed compact iPSC-like colonies (Fig. 1e-g). The two outcomes were mutually exclusive: the same cultures that gained ORO-positive structures yielded almost no Nanog-positive colonies at day 10 (Fig. 1f.h). Rather than accelerating pluripotency, as in pre-B and B cells^20, 21, 24^, Cebp factors diverted fibroblasts onto an adipogenic trajectory.

The switch was cell autonomous. At day 6, LipidTOX-positive lipid accumulation co-registered almost completely with Flag-positive cells (Fig. 1i,j), and live-cell imaging of MEFs co-transduced with ZsGreen-linked inducible hCebpα or hCebpβ captured individual cells acquiring adipocyte-like morphology between days 3 and 4, coincident with a pause in cell division (Fig. 1k,l). Varying the duration of induction before dox withdrawal (Fig. 1m) showed that terminal lipid accumulation increased with pulse length, consistent with asynchronous commitment. Nevertheless, a 3-5-day pulse was sufficient to commit cells, which retained their adipocyte fate at day 10 without further induction.

This adipogenic-iPS reprogramming antagonism was restricted to a subset of the family. Of five inducible human paralogues tested, only hCebpα, hCebpβ, and hCebpδ produced ORO-positive cells and suppressed Nanog-positive colony formation; hCebpε and hCebpζ were inert despite comparable nuclear expression (Supplementary Fig. 1b–g). The switch operated across mouse and human homologs, independently of reprogramming factor composition and delivery order (Supplementary Fig. 1h–k), and even under the N2/B27/2i conditions used for B-cell-to-iPSC reprogramming ^21^ (Supplementary Fig. 1l,m).

### Cebpα drives embryonic stem cells toward a trophectoderm-like fate

To determine whether the adipogenic outcome is intrinsic to fibroblasts or gated by cell identity, we expressed Cebpα in cells spanning a range of identities, beginning with pluripotent cells. Cebpα is expressed in the trophectoderm^29^ and potentiates trophectoderm differentiation in mouse ESCs^30^. We generated a dox-inducible knock-in ESC line (Col1a1:tetO-Cebpα-ZsGreen; ROSA26:M2rtTA) and followed it by immunofluorescence (24–72 hpi) and RNA-seq (24 and 48 hpi) (Fig. 2a). Within 72 h, Cebpα-expressing colonies lost their compact, dome-shaped morphology and adopted a flattened, scattered appearance but accumulated no LipidTOX-positive lipid (Fig. 2b).

**Fig. 2.**
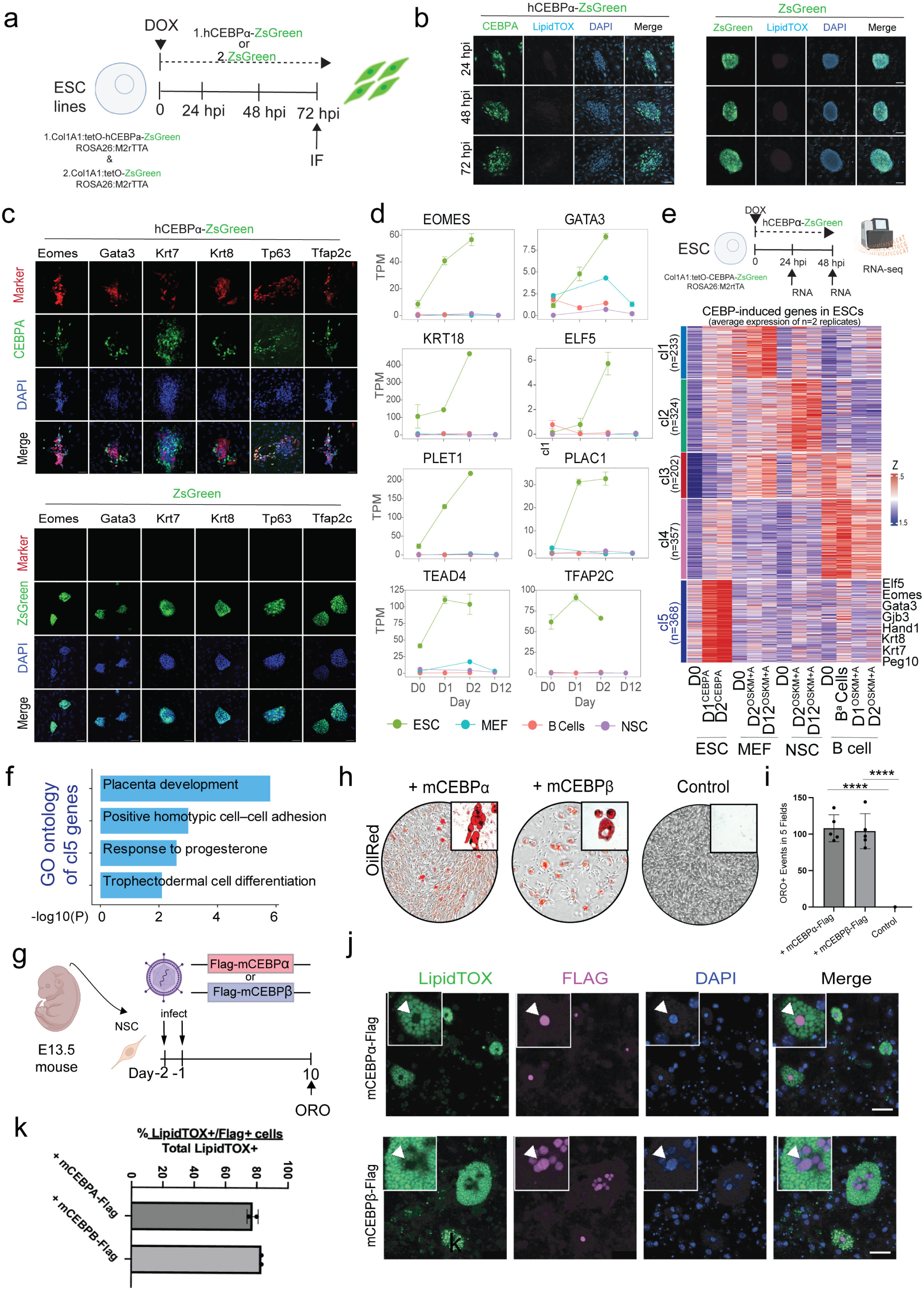
Cebpα induces a trophectoderm-associated identity in ESCs and adipocyte-like conversion in NSCs. **a**, Experimental schematic for Cebpα induction in bona fide embryonic stem cells (ESCs). Dox-inducible Cebpα-ZsGreen ESCs (Col1a1:tetO-Cebpα-ZsGreen; ROSA26:M2rtTA); immunofluorescence at 24, 48 and 72 h post-induction (hpi) and RNA-seq at 24 and 48 hpi. **b**, Immunofluorescence of Cebpα-ZsGreen ESCs (left) and control ESCs (right) at 24, 48 and 72 hpi, stained for Cebpα/ZsGreen (green), LipidTOX (cyan) and DAPI (blue). Cebpα-expressing colonies flatten and scatter without LipidTOX-positive lipid accumulation. Scale bar, 20 μm. **c**, Immunofluorescence of Cebpα-ZsGreen ESCs (left) and controls (right) at 72 hpi, stained for the trophectoderm markers Eomes, Gata3, Krt7, Krt8, Tp63, and Tfap2c (red), ZsGreen (green), and DAPI (blue). Scale bar, 20 μm. **d**, TPM expression of trophectoderm genes (*EOMES*, *TFAP2C*, *TEAD4*, *GATA3*, *PLET1*, *PLAC1*, *ELF5*, *KRT18*) at days 0, 1, 2 and 12 post-induction in ESCs (green), MEFs (cyan), B cells (red) and NSCs (purple). Data are mean ± s.e.m. **e**, Top: RNA-seq schematic; dox-induced Cebpα-ZsGreen ESCs were collected at 24 and 48 hpi. Bottom: heatmap of differentially expressed genes across ESC, Cebpα-induced ESCs (Day1, Day2), MEF, MEF-derived Day2^OSKM+A^ and MEF-derived D12^OSKM+A^, NSC, NSC-derived Day2^OSKM+A^ and NSC-derived D12^OSKM+A^, primary B cells, and B cells expressing CEBPα alone (B^α^) or with OSKM for 1 (D1^OSKM+A^) or 2 days (D2^OSKM+A^); Define five clusters (n = 233, 324, 202, 357, 368 genes); Z-score-normalized; trophectoderm genes annotated. **f**, Gene Ontology (GO) enrichment of the cluster-5 genes in e; −log_10_(*P*). **g**, Schematic of the Cebpα/β adipogenic-potential assay in neural stem cells (NSCs). NSCs from E13.5 embryos were transduced with retroviral Flag-mCebpα, Flag-mCebpβ, or empty vector and assayed by ORO staining at day 10. **h**, Representative brightfield/ORO-stained (red) images at day 10 of NSCs expressing mCebpα, mCebpβ or empty vector (control); insets show high-magnification ORO-positive lipid droplets. **i**, Quantification of ORO-positive events across five microscopic fields per experiment (mean ± s.d.; n = 5). \*\*\*\**P* < 0.0001, unpaired two-tailed Student’s t-test. **j**, Immunofluorescence of NSCs expressing Flag-mCebpα (top) or Flag-mCebpβ (bottom) at day 6, stained for LipidTOX (green), Flag (magenta) and DAPI (blue); merge at right. Arrowheads and insets show high-magnification cells co-positive for LipidTOX and Flag. Scale bar, 50 μm. **k**, LipidTOX-positive/Flag-positive cells as a percentage of total LipidTOX-positive cells in Flag-mCebpα and Flag-mCebpβ NSC cultures at day 6 (mean ± s.d.; n = 2).

Instead, Cebpα activated a trophectoderm-associated identity. By 72 hpi, Cebpα-expressing colonies induced EOMES, GATA3, KRT7, KRT8, TP63 and TFAP2C, none detected in controls (Fig. 2c). The response was directed rather than a generic loss of pluripotency: a trophectoderm panel (*Eomes*, *Tfap2c*, *Tead4*, *Gata3*, *Plet1*, *Plac1*, *Elf5*, *Krt18*) was progressively upregulated in Cebpα-induced ESCs, but not in the parallel MEF, NSC or B-cell time courses (Fig. 2d). K-means clustering across all four systems resolved five modules (Fig. 2e), of which the ESC-restricted cluster 5 was enriched with genes associated with ontologies related to placenta development and trophectoderm differentiation (Fig. 2f) and induced exclusively in Cebpα expressing ESCs (Supplementary Fig. 2a).

### Cebpα redirects neural stem cells toward an adipocyte-like fate

We next asked how a multipotent somatic stem cell would respond. Adherent Neural stem cells (NSCs) were derived from E13.5/E14.5 neurospheres and confirmed by Sox1, Sox2, Vimentin, and Neurod1 expression (Supplementary Fig. 2b,c), generated lipid-droplet-laden, ORO-positive adipocyte-like cells by day 10 upon retroviral expression of Flag-mCebpα or -mCebpβ, whereas empty-vector NSCs did not (Fig. 2g–i). As in fibroblasts, conversion was cell-autonomous; LipidTOX-positive staining co-registered almost completely with Flag-positive cells at day 6 (Fig. 2j,k), and the adipogenic response again dominated when Cebpα or Cebpβ was co-expressed with OSKM (Supplementary Fig. 2d-g). NSCs, therefore, phenocopy fibroblasts.

### Cebp reroutes OSKM reprogramming towards a bona fide adipogenic program

To define the transcriptional basis of this fate switch, we carried out bulk RNA-seq on MEFs and their OSKM, OSKM + Cebpα and OSKM + Cebpβ derivatives at day 2 and day 12 post induction (Fig. 3a). Cultures were lysed in situ, since adipocytes are buoyant and fragile and are readily lost on dissociation, so that converting, converted, and non-converted cells were captured alike. Principal-component analysis resolved two trajectories from the common MEF origin: OSKM alone progressed toward an iPSC state, whereas OSKM with Cebpα or Cebpβ moved along an orthogonal axis toward an adipocyte (iAdipocyte) state (Fig. 3b), a rerouting rather than a block.

**Fig. 3.**
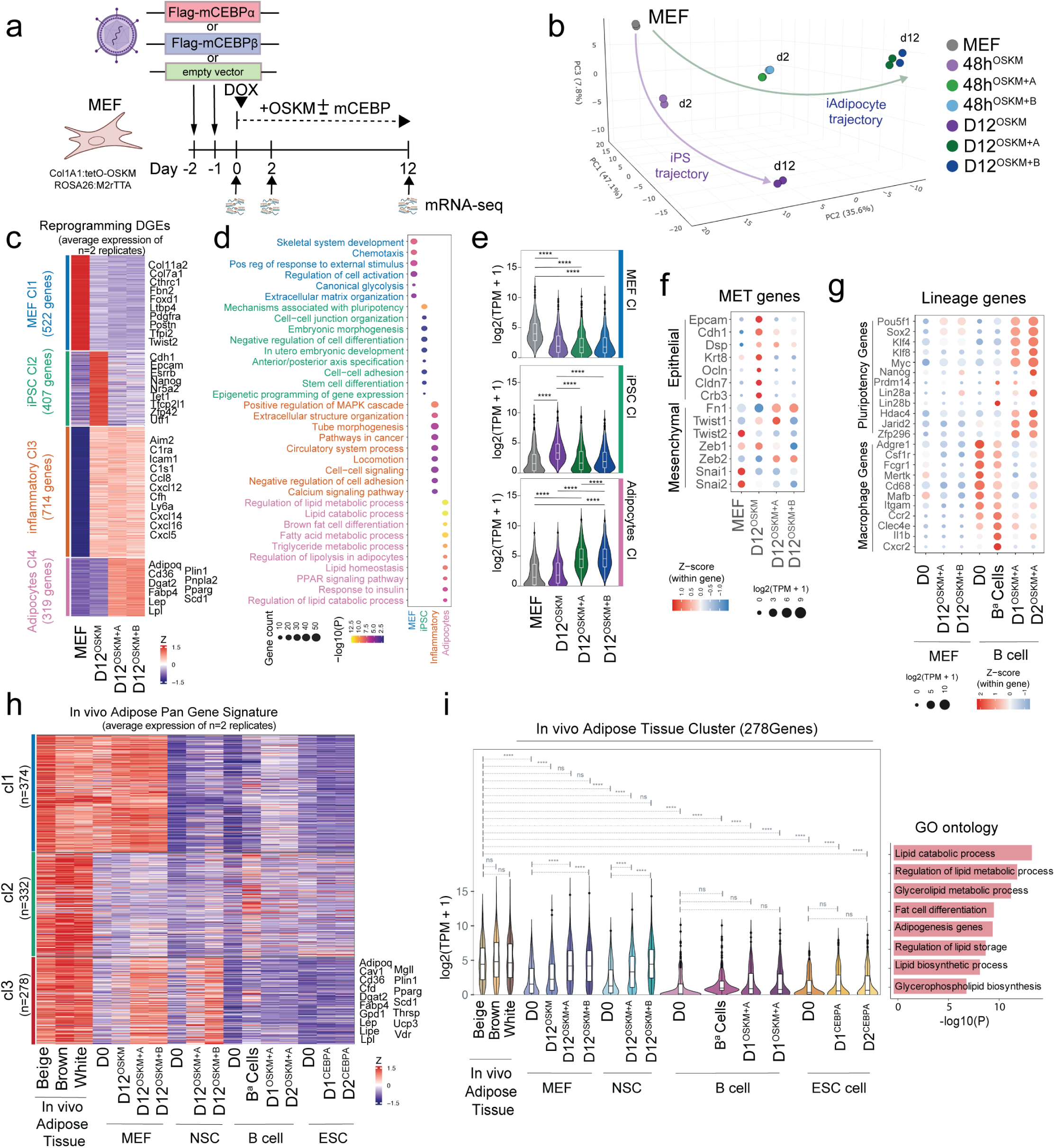
Cebpα/β rewires the MEF transcriptome to drive adipocyte transdifferentiation. **a**, Schematic of the bulk RNA-seq design. OSKM induced 48 h after mCebpα, mCebpβ or empty-vector infection of reprogrammable MEFs; samples at day 0 and days 2 and 12. **b**, Principal-component analysis (PCA) of bulk RNA-seq across MEF, OSKM, OSKM + mCebpα and OSKM + mCebpβ at 48 h and day 12 post-induction; two divergent trajectories, iPSC (purple) and iAdipocyte (green). **c**, k-means-clustered heatmap of differentially expressed genes across MEF, MEF-derived D12^OSKM^, MEF-derived D12^OSKM+A,^ and MEF-derived D12^OSKM+B^ samples (Z-score-normalized), resolving four clusters: MEF (522 genes), iPSC (407), Inflammatory (714), and Adipocyte (319) associated clusters (annotation by ontology); representative genes annotated at right. **d**, GO biological-process enrichment of the clusters in c; dot size indicates gene count and color indicates −log_10_(*P*). **e**, Violin plots of aggregate log_2_(TPM + 1) expression of the MEF, iPSC and Adipocyte cluster genes across MEF, MEF-derived D12^OSKM^, D12^OSKM+A^ and D12^OSKM+B^ samples. Asterisks mark significant differences between conditions (two-sided Wilcoxon rank-sum test with Benjamini–Hochberg correction for multiple comparisons; \*\*\*\**P* < 0.0001). **f**, Dot plot of epithelial (*Epcam*, *Cdh1*, *Dsp*, *Krt8*, *Ocln*, *Cldn7*, *Crb3*) and mesenchymal (*Fn1*, *Twist1*, *Twist2*, *Zeb1*, *Zeb2*, *Snai1*, *Snai2*) marker expression across MEF, MEF-derived D12^OSKM^, D12^OSKM+A^ and D12^OSKM+B^ samples; dot size indicates mean log_2_(TPM + 1) and color indicates within-gene Z-score. The mesenchymal-to-epithelial transition (MET) evident in D12^OSKM^ samples is absent in the Cebp-co-expressing conditions. **g**, Dot plots of pluripotency-and macrophage-associated gene expression across MEF, MEF-derived D12^OSKM+A^ and D12^OSKM+B^, primary B cells, and B cells expressing CEBPα alone (B^α^) or with OSKM for 1 (D1^OSKM+A^) or 2 days (D2^OSKM+A^). **h**, k-means-clustered heatmap across in vivo adipose subtypes (GSE198107; beige, brown, white); MEF, MEF-derived D12^OSKM^, D12^OSKM+A^ and D12^OSKM+B^, primary B cells, and B cells expressing CEBPα alone (B^α^) or with OSKM for 1 (D1^OSKM+A^) or 2 days (D2^OSKM+A^); NSC, NSC-derived D12^OSKM+A^ and D12^OSKM+B^, and ESCs expressing Cebpα for 1 or 2 days (Z-score-normalized). A 278-gene cluster (bottom, red) is selectively co-enriched in OSKM + Cebp-induced MEFs and NSCs and in all three in vivo adipose references; representative genes annotated at right. **i**, Left: violin plots of the 278-gene cluster (red; h) for the samples in h. Asterisks mark significant differences between conditions (two-sided Wilcoxon rank-sum test with Benjamini–Hochberg correction for multiple comparisons; \*\*\*\**P* < 0.0001; ns, not significant). Right: GO biological-process enrichment of the 278-gene cluster; −log_10_(*P*).

Unsupervised clustering resolved four gene programs: MEF (522 genes), iPSC (407), inflammatory (714) and adipocyte (319) confirmed by Gene Ontology (Fig. 3c,d). OSKM alone repressed the MEF program and activated the iPSC program^5, 31^, whereas adding Cebpα or Cebpβ left the iPSC program silent and induced an adipocyte gene network (Fig. 3c,e). The mesenchymal-to-epithelial transition (MET), the first committed event in MEF-to-iPSC reprogramming^32, 33^ failed: OSKM induced Epcam, Cdh1, and Krt8, whereas no epithelial induction occurred with the addition of Cebp (Fig. 3f), placing the block at the onset. Fibroblast identity was nonetheless dismantled in all conditions, with Twist1/2 downregulated in the OSKM-containing conditions^32–34^ and core fibroblast genes including Fbn2 and collagens^35, 36^ also downregulated (Fig. 3c,f). Cells therefore exit the fibroblast state without epithelializing. The adipogenic state was also discrete: at day 12, neither the OSKM pluripotency signature^5^ nor the myeloid program Cebp installs in B^17, 18^ or T cells^37^ was engaged (Fig. 3g and Supplementary Fig. 3f).

The induced program included the hallmark machinery of mature fat cells. *Pparg*, perilipin-1 (*Plin1*), the lipases *Lpl* and *Lipe*, caveolins (*Cav1/2*) and the adipokines adiponectin (*Adipoq*) and leptin (*Lep*), of which *Adipoq* and *Plin1* are essentially restricted to terminal adipocytes^13, 38, 39^ (Fig. 3c). A 278-gene module was co-enriched in OSKM + Cebp MEFs and in all three in vivo adipose references (beige, brown, and white fat)^40^, with lipid-and adipocyte-differentiation ontologies (Fig. 3h,i). Transcription-factor clustering exposed the switch beneath this: OSKM alone induced the core pluripotency network (Esrrb, Nanog, Sall4, Lin28a), which remained off whenever Cebp was present, whereas OSKM + Cebp induced *Pparg*, *Mlxipl* and *Nr1h3* (Supplementary Fig. 3d). The surface-marker repertoire shifted in parallel toward an adipocyte profile ^41–45^ (Supplementary Fig. 3e).

### Neural stem cells converge on the same adipocyte gene regulatory network as fibroblasts

We next profiled the NSC conversion by RNA-seq. Cebpα and Cebpβ, alone or with OSKM, induced an adipocyte-remodeling program and repressed the resident NSC and stem-cell-proliferation programs, while the ESC/pluripotency program remained off (Supplementary Fig. 3a,b,f). Each factor gave the signature alone as with OSKM, indicating that the Cebp factor specifies the output. The two differed in a second module: Cebpβ induced an adipocyte-metabolism program (*Pparg*, *Plin1*, *Lep*, *Lpl*, *Lipe*) and Cebpα a myelination program (*Sox10*, *Mbp*, *Plp1*, *Mog*, *Mag*) (Supplementary Fig. 3a–c). Fibroblasts and NSCs nonetheless converged on the same adipocyte program. The 278-gene adipocyte module was co-enriched in both systems and in all three in vivo adipose references (Fig. 3h,i), as were the core adipogenic regulators *Pparg*, *Cebp*α, *Cebp*β, *Nr1h3* and *Mlxipl* (Supplementary Fig. 3a,d). Lineage choice thus varies with the starting cell, but once the adipocyte fate is selected, it is established through transcriptional programming of a common program.

These comparisons resolve two questions. First, the OSKM-coupled conversion of Cebp-induced fibroblasts is direct transdifferentiation rather than reprogramming followed by re-differentiation: pluripotent cells respond to Cebpα by converting toward trophectoderm rather than fat (Fig. 2c-f), so a pluripotent intermediate would be expected to yield trophectoderm. Adipogenic output within 3-5 days (before iPSC colonies arise Fig. 1i–l), therefore reflects Cebp acting on cells still in a somatic state. Second, lineage outcome does not follow developmental potency: a multipotent NSC and a lineage-restricted fibroblast converge on adipocyte, whereas the pluripotent ESC diverges into trophectoderm. Lineage choice is set by the identity of the starting cell rather than by its position on the potency hierarchy.

### Cebp levels reshape genome-wide binding to induce adipogenic genes and silence the fibroblast and NSC programs

To ask how Cebp binding produces this transcriptional switch, we mapped Cebpβ occupancy by ChIP-seq in MEFs and in MEFs induced for 48 h with OSKM or OSKM + Cebpβ, partitioned bound sites into seven binary occupancy clusters (digits denoting binding in MEF, 48 h OSKM and 48 h OSKM + Cebpβ; Fig. 4a and Supplementary Fig. 4a). Two distinct redistributions were apparent. OSKM alone moved endogenous Cebpβ onto one set of new sites (cluster 010), as previously reported^5^, whereas raising Cebpβ levels engaged a separate, largely non-overlapping set of de novo sites (cluster 001). Cebpβ overexpressed alone in MEFs occupied the same de novo sites (Supplementary Fig. 4b), placing the redistribution under Cebpβ dosage rather than OSKM context. These 001 sites became accessible and H3K27ac-marked only when Cebpβ was elevated (Fig. 4b, c) and were neither accessible nor active in pre-iPSCs or ESCs. Overexpressed Cebpα and Cebpδ engaged the same locations (Supplementary Fig. 4c,d), making the level-dependent redistribution a shared family property consistent with the common adipocyte phenotype produced by all three factors.

**Fig. 4.**
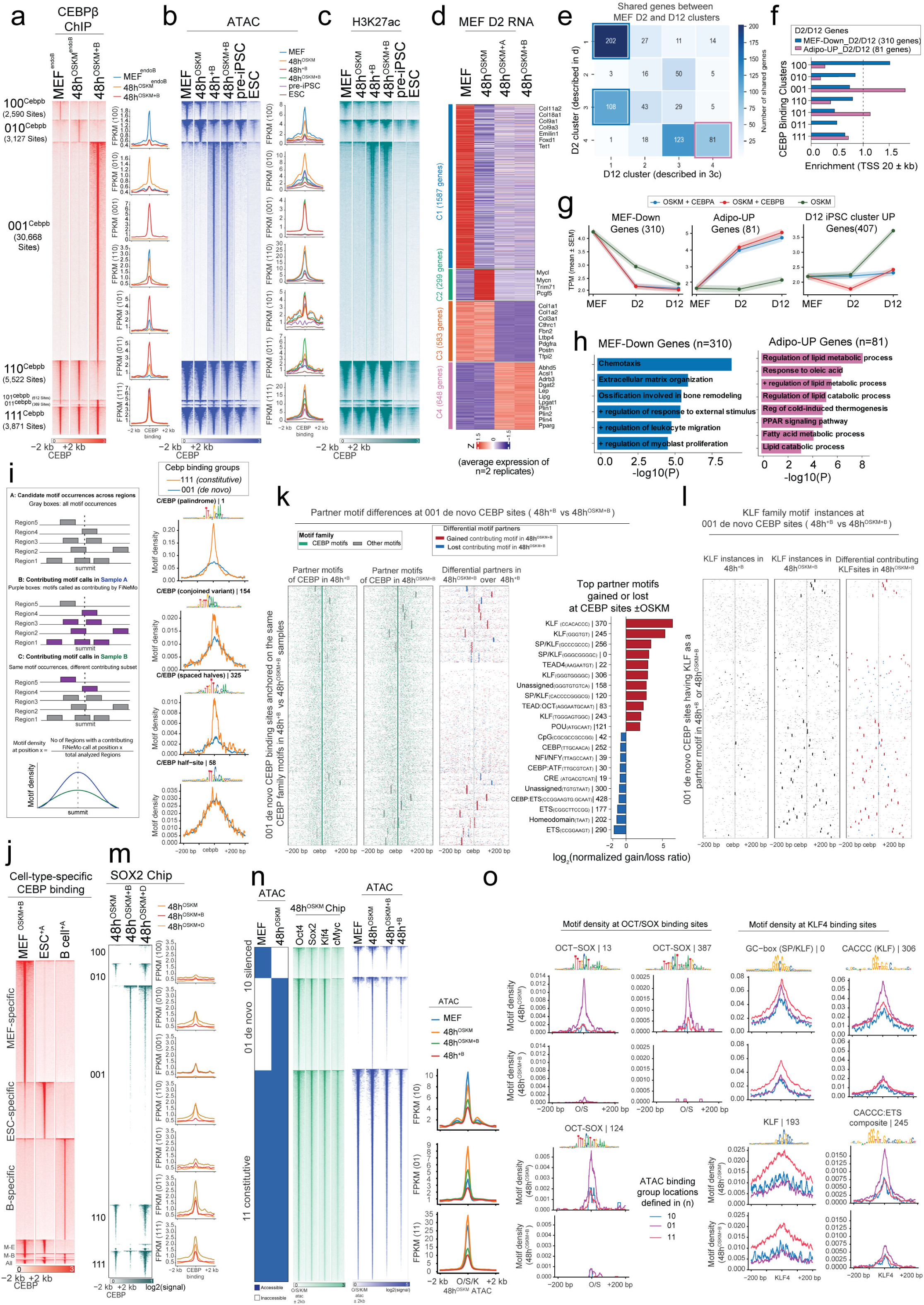
Cebpβ overexpression redistributes its occupancy to de novo enhancers associated with adipogenic gene induction and KLF recruitment. **a**, Cebpβ ChIP-seq at Cebpβ-bound sites, as described in S4a, in MEF and in MEFs induced 48 h with OSKM or OSKM + Cebpβ (±2 kb); mean-density metaplots (FPKM) at right. Seven binary occupancy clusters; digits denote binding in MEF, 48 h OSKM and 48 h OSKM + Cebpβ. **b**, ATAC-seq heatmaps and mean-density metaplots at the sites in a (±2 kb) for MEF, 48 h Cebpβ ± OSKM, pre-iPSC and ESC. **c**, H3K27ac ChIP-seq heatmaps at the sites in a, ordered as in b, for MEF, 48 h Cebpβ ± OSKM, pre-iPSC and ESC. **d**, k-means clustering of differentially expressed genes (row Z-score) across MEF and MEFs induced for 48 h with OSKM, OSKM + Cebpα or OSKM + Cebpβ, resolving four clusters: C1-4; representative genes at right. **e**, Overlap between the MEF day-2 gene clusters (C1–C4 from d; rows) and the MEF day-12 clusters (cl1–cl4, corresponding to the MEF, iPSC, inflammatory and adipocyte programs; Fig. 3c; columns). Cell color and number indicate the number of shared genes. **f**, Enrichment of the Cebpβ occupancy clusters from a within ±20 kb of TSSs, for genes commonly downregulated at days 2 and 12 (MEF-Down_D2/D12; n = 310) or commonly upregulated by day 2 within the day-12 adipocyte cluster (Adipo-UP_D2/D12; n = 81). Dashed line, enrichment = 1. **g**, Expression trajectories (TPM; mean ± s.e.m.) across MEF, day 2 and day 12 for OSKM + Cebpα (blue), OSKM + Cebpβ (red) and OSKM alone (green): commonly downregulated genes (MEF-Down; n = 310; left), commonly upregulated genes (Adipo-Up; n = 81; middle) and day-12 iPSC-cluster upregulated genes (n = 407; right). **h**, GO biological-process enrichment for the downregulated (n = 310; left) and upregulated (n = 81; right) gene sets in g; −log_10_(*P*). **i**, Left, schematic of the motif-density metric (see Methods): FiNeMo calls the contributing motif instances (purple) among all candidate occurrences (grey); density at position *x*, fraction of regions with a contributing FiNeMo call at *x*. Right, motif density of Cebp-family motifs (CEBP|1, CEBP|154, CEBP|325, CEBP|58) at 111 constitutive and 001 *de novo* Cebpβ binding groups, centered on the Cebpβ summit (±200 bp). **j**, Cell-type-specific Cebp occupancy. Cebpα/β ChIP-seq at Cebp-bound sites in OSKM + Cebpβ MEFs, Cebpα-ESCs and Cebpα-B cells (±2 kb); digits denote binding in MEF, ESC and B cell: 100 MEF-specific, 010 ESC-specific, 001 B-cell-specific. **k**, CEBP-centric view of contributing partner motifs at 001 *de novo* CEBP sites: CEBP overexpression (48h^+B^) versus CEBP + OSKM (48h^OSKM+B^). Heatmaps, contributing motif instances anchored on the CEBP motif (±200 bp); CEBP motifs green, others grey. Right, per-site difference, gained (red) or lost (blue) in 48h^OSKM+B^, revealing a net gain with OSKM. Far right, differential motifs ranked by log₂(normalized gain/loss ratio). **l**, KLF-family motif instances at the subset of 001 *de novo* CEBP sites carrying a KLF partner motif, anchored on the CEBP motif and centered on the summit (±200 bp). Left, KLF instances in 48h^+B^; middle, 48 h^OSKM+B^; right, differential contributing KLF instances, gained (red) or lost (blue) in 48h^OSKM+B^. KLF is preferentially gained in 48h^OSKM+B^ samples. **m**, SOX2 ChIP-seq at the seven Cebpβ binding clusters defined in a, centered on peak summits (±2 kb); per-cluster mean-density metaplots (FPKM) at right for 48h^OSKM^ (orange), 48h^OSKM+B^ (red), and 48h^OSKM+D^ (grey). **n**, ATAC accessibility clusters between MEF and 48h OSKM-induced MEF (binary block, left): 10 (MEF-only), 01 (*de novo* in 48h^OSKM^) and 11 (shared accessibility); digits, accessibility in MEF and 48h^OSKM^. Middle, ChIP-seq (Oct4, Sox2, Klf4, cMyc; GSE90895) and ATAC (MEF, 48h^OSKM^, 48h^OSKM+B^, 48h^+B^), centred on 48h^OSKM^ sample ATAC summits (±2 kb). Right, ATAC metaplots (FPKM): MEF, blue; 48h^OSKM^, orange; 48h^OSKM+B^, green; 48h^+B^, red. **o**, Motif density at ATAC sites, grouped by accessibility cluster between MEF and 48h^OSKM^ -induced MEF (10, blue; 01, purple; 11, red; from n). Left, OCT/SOX identified motifs (OCT/SOX|13, |387, |124) at ATAC sites under OCT/SOX ChIP-seq peaks; right, KLF identified motifs SP/KLF|0, CACCC/KLF|306, GCbox/KLF|193, CACCC:ETS|245) at ATAC sites under KLF4 ChIP-seq peaks. Motif density shown for 48h^OSKM^ (top of each pair) and 48h^OSKM+B^ (bottom), centered on the ChIP-seq summit (±200 bp).

This *de novo* (001) binding was transcriptionally productive. Already at day 2, OSKM + Cebpα and OSKM + Cebpβ shared an early induction of lipid and adipocyte genes including *Pparg*, *Plin1* and *Lep* that was absent under OSKM alone (Fig. 4d and Supplementary Fig. 4e). To follow sustained rather than transient changes, we intersected the MEF day-2 and day-12 expression clusters and retained genes moving consistently in one direction at both timepoints (Fig. 4e): a commonly downregulated set (MEF-Down; n = 310), and a commonly upregulated set drawn from the shared adipocyte cluster (Adipo-Up; n = 81). Relating Cebp binding to each set (Fig. 4f), MEF-Down genes were enriched for endogenous Cebpβ occupancy (the 100 sites bound in MEFs and lost on induction), while Adipo-Up genes enrich for the de novo (001) sites.

Expression trajectories separated the Cebp-specific events from the OSKM-driven ones (Fig. 4g), with chemotaxis and extracellular-matrix ontologies confirming the MEF identity of MEF-Down and lipid-metabolic ontologies the adipocyte identity of Adipo-Up (Fig. 4h). Adipo-Up genes rose only under OSKM + Cebpα or OSKM + Cebpβ and stayed low under OSKM alone, so adipocyte induction is contributed specifically by Cebp. MEF-Down genes fell comparably across all three conditions, consistent with silencing driven by loss of endogenous Cebpβ from the 100 maintenance sites (active MEF enhancers; Supplementary Fig. 4f), which OSKM triggers with or without added Cebp. Day-12 iPSC-cluster genes behaved oppositely, rising only under OSKM and held down by Cebpα or Cebpβ.

The same binding-to-expression logic held in neural stem cells: Cebpα and Cebpβ opened the de novo sites defined in fibroblasts (Supplementary Fig. 4g), an equivalent NSC response resolved analogous gene sets (NSC-Down, n = 85; Adipo-Up, n = 278; Supplementary Fig. 4h,i,k,l), and the de novo sites were again enriched near the induced adipocyte genes (Supplementary Fig. 4j). Elevated Cebp did not engage this repertoire in ESCs (Supplementary Fig. 4m) or B cells (Supplementary Fig. 4n), even though Cebp alone suffices to open it in MEFs (Supplementary Fig. 4b). Elevated Cebp is therefore required but not sufficient for engagement; the cell of origin sets which of the sampled sites can be opened.

### A concentration-driven spillover model explains de novo Cebp binding

To ask how transcription factor level sets the binding repertoire rather than merely its occupancy, we used ChromBPNet^46^ to infer the sequences responsible for accessibility at the de novo (001) and constitutive Cebp binding (111) sites. Motif discovery with TF-MoDISco^47^, which builds motifs from per-base contribution rather than sequence frequency, resolved these sequences into contribution weight matrices (CWMs), and FiNeMo called the instances driving accessibility, giving a contributing-motif density (Fig. 4i). Because a position-weight-matrix (PWM) scan cannot separate a sequence that acts from one that merely recurs^48–50^, CWM counts are preferred as they measure potential engagement rather than sequence content.

Four Cebp CWMs were recovered, named for their similarity to the canonical CEBP motifs in JASPAR^51, 52^. CEBP proteins bind as bZIP dimers to the dyad ATTGCGCAAT, a fusion of two ATTGC half-sites, whose spacing is set by the basic-region residues that read each half^53, 54^, though most occupied sites in cells are fusions of a canonical with a degenerate half-site rather than the full palindrome^55^. The four CWMs spanned this range: CEBP_1 (TTGCGCAAT), the conjoined palindrome; CEBP_154 (TGTTGCAACA), a conjoined variant with a shortened core; CEBP_325 (TTGTTGC---CAA), spaced half-sites; and CEBP_58 (TTGCACA), a single canonical half site with a degenerate partner. All four underpinned both site classes, what differed was their relative contribution. The three dimer-type variants contributed at higher density across the constitutive sites that maintain fibroblast identity than across the de novo sites, whereas the half-site CEBPA_58 contributed equally at both, making the constitutive sites motif-rich and the de novo sites motif-poor for Cebp (Fig. 4i).

The classes, therefore, differ in the relative abundance of Cebp sequences across regions. We term this effect ‘concentration-driven spillover’: motif-rich regions are occupied at basal factor levels and sustain fibroblast identity, whereas motif-poor regions are engaged only as Cebpβ accumulates. Because the same four CWMs are available to both classes (Fig. 4i), this gradient sets how many sites are probed, but cannot specify which are chosen; that selection must come from sequences outside the Cebp element. These 001 sites lie outside the normal reprogramming route, being the only binding groups not enriched in the chromatin trajectories that remodel during MEF-to-iPSC conversion (Supplementary Fig. 4f).

Cebp motifs vastly outnumber the bound sites, and overexpressed Cebp occupies a largely non-overlapping repertoire in MEFs, ESCs, and B cells (Fig. 4j and Supplementary Fig. 4m,n and Supplementary Fig. 5a). In MEFs, the de novo (001) sites are engaged only when Cebpβ is elevated (Fig. 4a) and are occupied whether or not OSKM is present (Fig. 4a and Supplementary Fig. 4b), so the partner elements contributing to their accessibility can be compared at identical locations.

With Cebpβ alone, the contributing partners fell into three groups. The first was a CCAAT box (TTGGCCAAT) read by NFI or NF-Y. The second comprised two bZIP elements, the cyclic-AMP response element (CRE, TGACGTCA) and the CEBP–ATF composite (TTGCGTCAT), in which a CEBP half-site (TTGC) is fused to a CRE half-site (GTCAT) and which is one of the heterodimer elements that account for around 40% of CEBP-occupied sites in cells^55^. The third was the class I ETS elements (CCGGAAGT) recovered as primary or fused to a CEBP half-site. Class I is shared across the ETS, TCF, ERG and PEA3 subfamilies, so the sequence specifies the class but not the factor^56^. These elements are nominated as co-factors that, alongside Cebp, may specify the adipogenic program (Fig. 4k,l, and Fig. 5c,d).

**Fig. 5.**
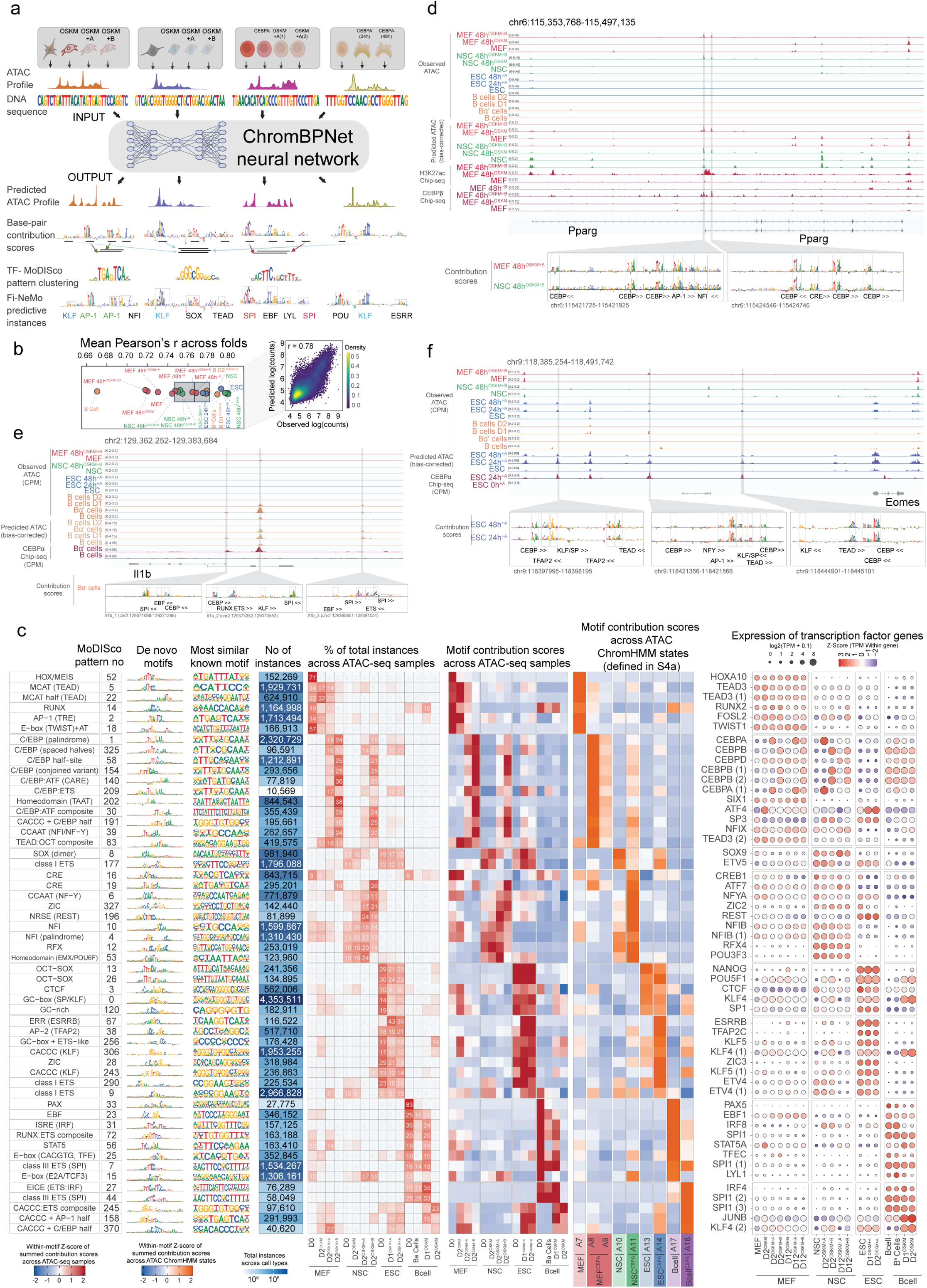
Cebp-driven chromatin-accessibility remodeling encodes cell-type-specific regulatory grammar across somatic and pluripotent contexts. **a**, ChromBPNet analysis pipeline. Cell-type-specific models were trained on ATAC-seq profiles from MEF, NSC, B-cell and ESC samples across basal and Cebp-and/or OSKM-induced conditions. DeepLIFT-derived per-base contribution scores were clustered with TF-MoDISco to build a de novo contribution-weight-matrix (CWM) motif repertoire, and motif instances were annotated genome-wide per condition with Fi-NeMo. **b**, ChromBPNet Model performance. Mean Pearson correlation between observed and predicted accessibility across cross-validation folds for each cell-type/condition model; inset, predicted versus observed log read counts (r=0.78). **c**, Summary of the most abundant de novo motifs identified by TF-MoDISco (one row per motif group). Left to right: putative associated TF; MoDISco pattern number; de novo motif as a CWM; most similar known TF binding motif (PWM); total genomic instances across all conditions and instances per condition; within-motif Z-score of the percentage of total instances across the ATAC-seq samples (MEF, NSC, ESC and B cell at basal state and under OSKM and/or Cebp induction); within-motif Z-score of summed motif contribution scores across the same samples; within-motif Z-score of summed contribution scores stratified by ChromHMM chromatin state; and expression of the corresponding TF genes (dot size, log_2_(TPM + 1); colour, within-gene Z-score) across MEF, NSC, ESC and B-cell conditions. Arrows indicate motifs whose contribution is concentrated in the adipogenic (NFI and NF-Y), trophectoderm (TFAP2C) and myeloid (SPI1) compartments. **d**, The *Pparg* locus, a MEF/NSC-specific Cebpα-responsive gene, showing observed and predicted chromatin accessibility, H3K27ac and Cebpβ ChIP-seq, per-base contribution scores, and annotated motif instances (Cebp, NFI, CRE, ATF, and FOSL1+JUN) across all conditions. **e**, As in d for *Il1b*, a Cebpα-induced accessibility target in B cells, with annotated SPI, EBF, KLF, RUNX:ETS and Cebp motif instances. **f**, As in d for *Eomes*.

Adding OSKM alongside Cebpβ did not exchange the discovered motif set but added to it. The gained elements were dominated by KLF and SP elements (CACCC and GC boxes), together with the POU octamer (ATGCAAT) and TEAD, the latter recovered both free-standing (AAGAATGT) and fused to an octamer half (AGGAATGCAAT). KLF and octamer are the recognition sequences of two cocktail components, Klf4 and Oct4, and were nearly absent from these sites when Cebpβ was expressed alone, whereas the Cebp-alone partners persisted when OSKM was supplied alongside it, at only modestly reduced density (Fig. 4k,l). Because Cebpβ is present in both conditions, the same Cebp-bound DNA is read by an expanded set of partners when OSKM is added, not by a substitute set.

### Elevated Cebp levels reciprocally redistributes the Yamanaka factors

Our earlier work described OSKM relocating the fibroblast’s endogenous transcription factors as it engages the genome^5^,which reanalysis of endogenous Cebpβ confirmed here (the 010 cluster; Fig. 4a). However, at elevated levels, Cebpβ acts back on the reprogramming factors. Sox2 was recruited onto the Cebp binding groups, including the de novo 001 sites, specifically when Cebpβ or Cebpδ was co-expressed with OSKM (Fig.4m). Redistribution therefore runs in both directions. At the regions bound by Oct4, Sox2 and Klf4 immediately after OSKM induction (cluster 01; Fig. 4n), co-expressing Cebpβ reduced the contribution of Oct/Sox and Klf motifs to predicted accessibility (Fig. 4o), and the OSKM/iPSC target genes failed to be induced (Fig. 4g).

The composite read by Oct4-Sox2 heterodimers was not among the gained elements (in OSKM+Cebp conditions), so Sox2 arrives at the *de novo* sites without a site of its own. Retargeted Klf4, with its motif presence, could recruit Oct4 and Sox2 through protein-protein interactions^57^ and relocate the Yamanaka factors as a group (Fig. 4m). Elevated Cebpβ therefore draws Oct, Sox and Klf from the pluripotency-associated sites they engage early in reprogramming into the adipogenic repertoire, so the pluripotency program is never adequately established.

### Distinct resident transcription-factor grammar specifies cell identity across somatic and pluripotent contexts

To ask whether the partner logic generalizes, we trained cell-type-specific ChromBPNET models across the induction timecourse of each system (MEFs, NSCs, B cells and ESCs in the basal state and after Cebp and/or OSKM induction), so that the same models report both the motifs holding a cell in its starting identity and those engaged as Cebp drives it toward a new fate (Fig. 5a). The models reproduced experimental accessibility profiles with a mean Pearson’s correlation of 0.66-0.80 across cross-validation folds and tracked observed read counts genome-wide (r = 0.78; Fig.5b). Stratifying the contributing motifs by ChromHMM-defined differential accessibility (Supplementary Fig. 5a), and overlaying transcription factor expression separated the factors defining each starting identity from the partners Cebp engages to redirect it (Fig. 5c).

Each starting cell carried a largely distinct resident grammar. Motif contribution scored within enhancers and promoters resolved by a stacked ChromHMM^58^ model over nine histone modifications (Supplementary Fig. 5d) cataloged CEBP, AP-1, TEAD, TWIST and HOXA in MEFs; SOX, and RFX in NSCs; OCT-SOX, ESRRB, TFAP2 and ZIC in ESCs; and PAX, EBF, E2A, SPI, and IRF in B cells (Supplementary Fig. 5e), each set matching the factors that maintain the corresponding cell state ^5, 59–62^. NFI and NF-Y were shared between MEFs and NSCs, the two systems that converge on the adipocyte fate, while RUNX was common between MEFs and B cells. The same factors emerged independently from the accessibility models (Fig. 5c) and at the cell-type specific *Col6a, Olig1/2, Zfp42 and Cd19* loci (Supplementary Fig. 5g–j). This is the grammar Cebp must act upon.

### Context-specific partners direct Cebp toward an adipocyte, trophectoderm or myeloid fate

If the resident grammar is what Cebp must overcome, the *de novo* sites it opens at elevated levels define a second one: the partner elements it engages to install the new fate. Across the four systems, de novo Cebp binding (Fig. 4j; Supplementary Fig. 5b) opened a distinct set of regions in each route (Supplementary Fig. 5c). Factor level, therefore, sets how much de novo binding occurs (Fig. 4), but not where: a context-specific partner must direct it.

Three loci make this concrete: each was inaccessible in all four starting cell types and opened only where elevated Cebp met the partners supplied by its cell of origin. The adipogenic master gene *Pparg* ^63–65^ opened on Cebp induction in MEFs and NSCs, with NFI, an AP-1 site, and a CRE contributing alongside Cebp (Fig. 5d). The macrophage effector *Il1b*^17, 19^, opened only in Cebpa-pulsed B cells with SPI, EBF and RUNX-ETS sites contributing alongside Cebp (Fig. 5e); MEFs, NSCs and ESCs, which express little Spi1 or Ebf1 left it closed. The trophectoderm regulator Eomes^30, 66^, opened only in Cebpa-induced ESCs, with TFAP2, TEAD and KLF/SP sites contributing alongside Cebp (Fig. 5f). In every case the other systems remained closed despite comparable Cebp induction.

Genome-wide, the partner sets grouped the four systems by outcome (Fig. 5c). One group contributed in every system at every time point: the GC-box, CACCC, CTCF and CpG elements, CG-rich sequences that mark promoters rather than fate. The remainder were cell-type restricted and contained the putative Cebp partners. NFI, NF-Y, CRE and the CRE::CEBP composite grouped MEFs with NSCs, matching their common adipocyte route. ERR, AP-2 and ZIC marked ESCs. SPI, EBF, RUNX and the IRF::ETS composite marked B cells.

The ETS classes are partitioned in the same way, consistent with the expression of the corresponding factors. Class III (AGAGGAAGT) in B cells, class I (ACCGGAAGT) in NSC, MEF, and ESCs, the two differing only in the bases immediately 5’ of a shared GGAA^56^. The composite C/EBP::ETS (209) was the exception, enriched in every conversion. It fuses a complete C/EBP site to a bare ETS core (GGAA) in which those discriminating 5’ bases are absent, so the element requires an ETS partner without specifying which and is read by whichever class the cell expresses. Cebp output is therefore assembled from each cell’s own grammar: a resident identity it must overcome and a partner set, largely drawn from that same grammar, that installs the new fate.

### SPI1 redirects Cebpa output toward a dual adipogenic-and-myeloid fate

The macrophage route in B cells nominated SPI1 as the partner accompanying Cebpα (Fig. 5). To test whether supplying it redirects Cebpα output, we expressed the two in MEFs FACS-purified of CD45^+^, CD11b^+^, CD48^+^ and Ly6G^+^ cells, excluding pre-existing hematopoietic cells as a source of myeloid output (Supplementary Fig. 6a). The factors were delivered as one bicistronic cassette (3×Flag-hCebpα-E2A-hSpi1), so that every cell received both, with each factor alone as a reference (Fig. 6a and Supplementary Fig. 6a,b).

**Fig. 6.**
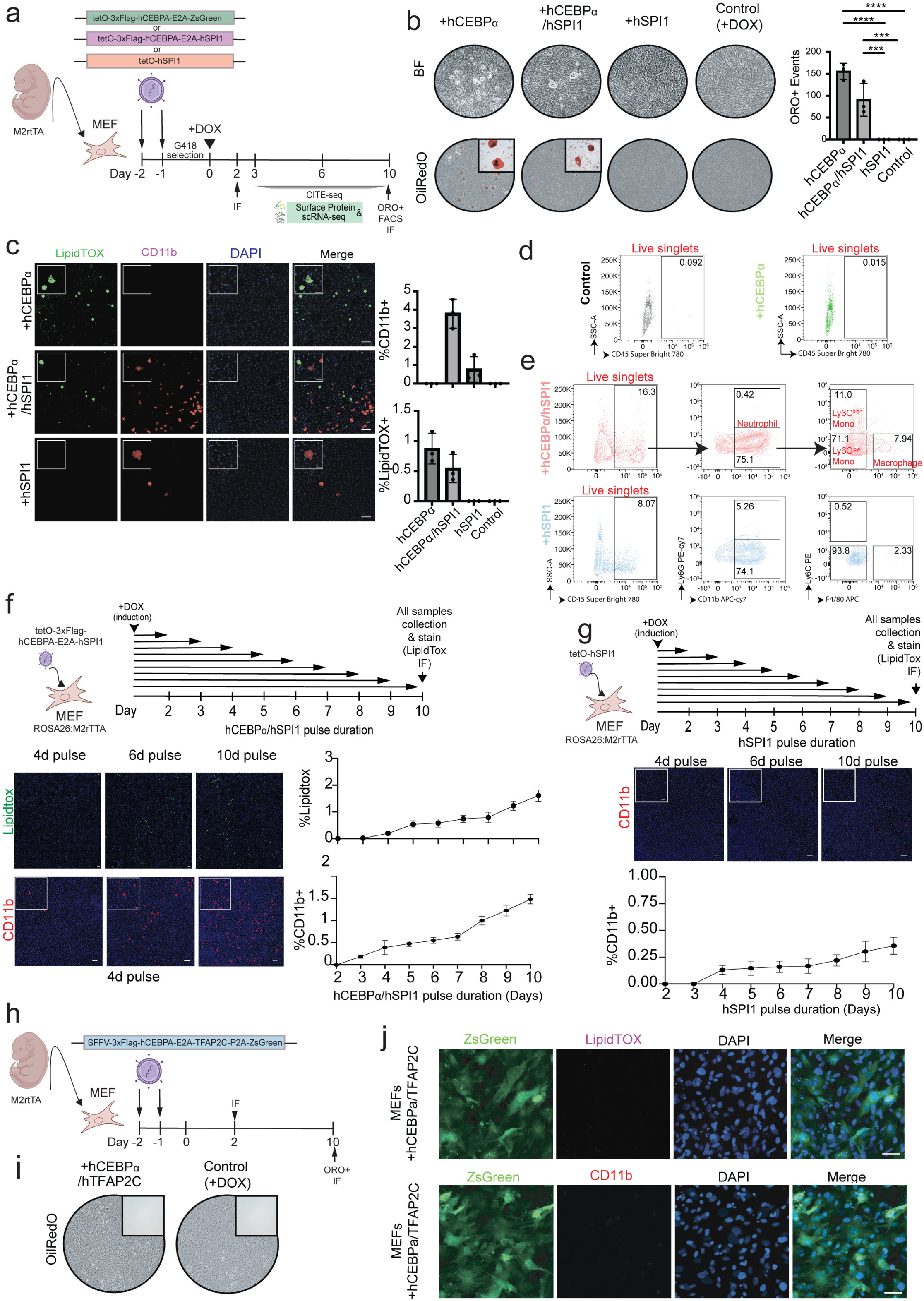
Partner factors redirect Cebpa output: SPI1 imposes a macrophage-like fate, TFAP2C abolishes adipogenic conversion. **a**, Combinatorial reprogramming assay. MEFs (ROSA26:M2rtTA) transduced with tetO-3×Flag-hCebpα-E2A-ZsGreen (Cebpα), tetO-3×Flag-hCebpα-E2A-hSpi1 (Cebpα/SPI1; single bicistronic cassette) or tetO-hSPI1 (SPI1),; G418-selected, dox from day 0 for 10 days; immunofluorescence at day 2, CITE-seq at days 3, 6 and 10, and ORO, flow cytometry and immunofluorescence at day 10. **b**, Brightfield (top) and ORO-stained (bottom; red) wells at day 10 for hCebpα, hCebpα/hSpi1, hSpi1 and dox-only control; insets, lipid droplets. Right, ORO-positive events across five fields (mean ± s.d.; n = 3). \*\*\*\**P* < 0.0001, \*\*\**P* < 0.001, unpaired two-tailed Student’s *t*-test. **c,** Immunofluorescence of MEFs expressing hCebpα (top), hCebpα/hSPI1 (middle) or hSPI1 (bottom) at day 10, stained for LipidTOX (green), CD11b (red) and DAPI (blue); merge at right. Insets show high-magnification views of the boxed regions. Right: percentage of CD11b-positive (top) and LipidTOX-positive (bottom) cells at day 10 across the hCebpα, hCebpα/hSPI1, hSPI1 and control (+dox) conditions (mean ± s.d.; n = 3). **d**, Flow cytometry at day 10, dox-only control (left) and hCebpα (right); live singlets gated on CD45 (Super Bright 780) versus SSC-A. Neither yields a CD45^+^ population (numbers, % CD45^+^ of live singlets). **e**, Flow cytometry at day 10, hCebpα/hSpi1 (top) and hSpi1 (bottom). Live singlets gated on CD45 versus SSC-A, then CD45^+^ cells on Ly6G versus CD11b (neutrophils, Ly6G^+^CD11b^+^) and on Ly6C versus F4/80 (Ly6C^high^ monocytes, Ly6C^low^ monocytes, macrophages [F4/80^+^]); numbers, % of parent gate. **f**, hCebpα/hSpi1 induction-withdrawal time course (tetO-3×Flag-hCebpα-E2A-hSpi1). Top, dox added at day 0 and withdrawn at defined intervals (days 2–9), continuously induced condition as control; all conditions collected at day 10. Bottom, LipidTOX (green) and CD11b (red) immunofluorescence at 4-, 6-and 10-day pulses, and LipidTOX-positive and CD11b-positive cells versus pulse duration (mean ± s.d.). A pulse of ≥3 days commits cells to both fates. **g**, As in f for SPI1 alone (tetO-hSpi1), stained for CD11b; a pulse of ≥3 days commits cells to a CD11b-positive fate. **h**, Cebpα/TFAP2C reprogramming assay. MEFs transduced with SFFV-3×Flag-hCebpα-E2A-hTfap2c-P2A-ZsGreen, selected with G418; analyzed by immunofluorescence at day 2 and by ORO and immunofluorescence at day 10. **i**, ORO-stained wells at day 10 for hCebpα/hTfap2c and dox-only control; no ORO-positive cells. **j**, Immunofluorescence of hCebpα/hTfap2c for ZsGreen (green) with LipidTOX (magenta; top) or CD11b (red; bottom) and DAPI (blue); merge, right. Neither adipocyte-like nor macrophage-like fate was produced.

Cebpα with SPI1 produced two phenotypes at once: ORO-and LipidTOX-positive adipocyte-like cells, at markedly lower frequency (∼50% reduction) than with Cebpα alone, and CD11b-positive cells never seen with Cebpα alone (Fig. 6b,c). A CD45-positive population arose under Cebpα/SPI1 and SPI1 but not Cebpα alone, representing predominantly monocytic and macrophage with a minor neutrophil fraction (Fig. 6d,e). Its composition differed by condition: SPI1 alone gave almost exclusively Ly6C-low monocytes, whereas adding Cebpα increased the CD45-positive fraction and shifted it toward Ly6C-high monocytes and F4/80-positive macrophages (Fig. 6e). Each factor therefore contributed its own fate. SPI1 alone produced no adipocytes alone, and adding it to Cebpα did not abolish the adipogenic output but roughly halved it, while adding a myeloid output Cebpα alone never gave. The two programs ran in parallel, with the share of cells entering each, and the maturity of the myeloid arm, set by the presence of the other rather than by one factor overriding the second. We call the CD11b-positive cells macrophage-like, following Feng et al^67^., and keep the term provisional, since surface flow gives only a suggestive read of monocyte-macrophage output.

Both fates arose while both factors remained expressed. At day 10, Flag-Cebpα and SPI1 colocalized in LipidTOX-positive adipocytes and in CD11b-positive cells, most scored cells being double-positive, while the adipogenic and myeloid markers themselves labelled non-overlapping cells (Supplementary Fig. 6c–e). The bifurcation was therefore not resolved by silencing one factor, leaving open what selects one fate over the other, which we address by single-cell profiling. Both new fates passed through the same commitment window as the adipocyte fate: a three-day pulse between days 3 and 5 enabled the respective outputs, which persisted to day 10 after dox withdrawal (Fig. 6f,g and Fig. 1m).

### TFAP2C abolishes Cebpa-driven adipogenic conversion

The trophectoderm route in ESCs had nominated a different partner, TFAP2C, posing the converse question: could a partner suppress rather than extend Cebpα’s output? Co-expressing the two in MEFs (SFFV-3×Flag-hCebpα-E2A-hTfap2c-ZsGreen; Supplementary Fig. 6f) yielded neither adipocyte-like cells, by ORO or LipidTOX, nor CD11b-positive cells (Fig. 6h–j), the cells retaining fibroblast morphology throughout despite co-expression of Flag and TFAP2C in the same ZsGreen-positive cells (Supplementary Fig. 6g,h). By these markers, TFAP2C blocked the adipogenic conversion Cebpα drives by default and permitted no myeloid one: a supplied partner that suppresses Cebpα’s output as readily as SPI1 extends it. This does not preclude the pair from installing a program of its own. Nominated via the trophectoderm route, the combination could engage a network partially related to trophectoderm that surface markers and morphology alone would not reveal.

### Single-cell multiomics resolves the states generated by Cebpα, SPI1, and their combination

MEFs transduced with hCebpα, hSPI1 or hCebpα/hSPI1 without OSKM were profiled by CITE-seq at days 3, 6 and 10 (Fig. 6a) and merged into an atlas of 110,751 cells (Fig. 7a; 38,214 / 37,101 / 35,436 per arm), with transgene and reporter detection confirming delivery in each arm (Supplementary Fig. 7a).

**Fig. 7.**
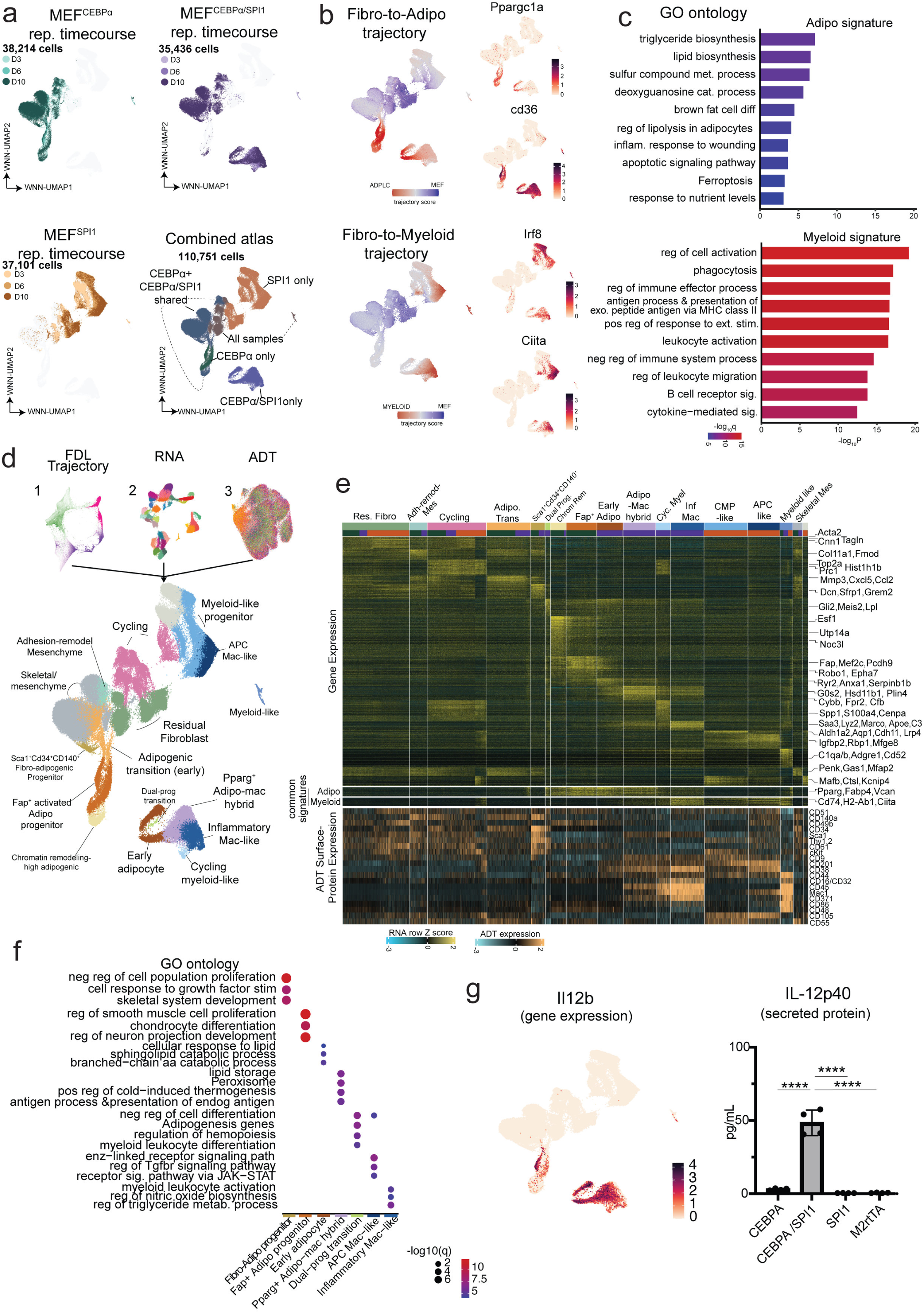
CITE-seq resolves parallel adipogenic and myeloid conversion and a dual-identity adipocyte– macrophage state. **a**, WNN-UMAP of CITE-seq time courses at days 3, 6 and 10: MEF^CEBPα^ (38,214 cells), MEF^CEBPα/SPI1^ (35,436) and MEF^SPI1^ (37,101), colored by day; combined atlas (110,751 cells) at bottom right. Dashed outlines, territory unique to or shared between conditions. **b**, Directional trajectory scores across the combined atlas: fibroblast-to-adipocyte (top) and fibroblast-to-myeloid (bottom). Right, *Ppargc1a* and *Cd36* (adipogenic) and *Irf8* and *Ciita* (myeloid); color, log-normalized gene expression. **c**, GO and pathway enrichment of the common adipocyte (*n* = 95 genes) and common myeloid (*n* = 83 genes) signatures defined in e; bar length, −log_10_(*P*); color, −log_10_(*q*). **d**, Force-directed layout trajectory (1), RNA (2) and ADT (3) combined by weighted-nearest-neighbour analysis; annotated embedding below, coloured by cell state. residual fibroblast; adhesion-remodeling mesenchyme; skeletal/mesenchymal; cycling; adipogenic transition (early); Sca1+Cd34+CD140a+ fibro-adipogenic progenitor; Fap+ activated adipogenic progenitor; chromatin-remodeling-high adipogenic; dual-program transition; early adipocyte; Pparg+ adipocyte-macrophage hybrid; cycling myeloid-like; inflammatory macrophage-like; common myeloid progenitor-like; antigen-presenting macrophage-like; myeloid-like. **e**, Stacked heat maps across the 16 states: population-signature genes (top; row Z-score), common adipocyte and common myeloid signatures (middle), and ADT surface proteins (bottom); representative genes and antigens at right. **f**, GO and pathway enrichment for seven of the cell states in d, computed on their signature genes from e; dot size, −log∼10∼(*q*); color, −log∼10∼(*P*). **g**, *Il12b* gene expression across the combined atlas; color, log-normalized expression. Right, IL-12p40 protein detection in day-10 supernatants of hCebpa, hCebpa/hSpi1, hSpi1 and dox-only (M2rtTA) cultures (mean ± s.d.; n = 4). ****P < 0.0001, unpaired two-tailed Student’s t-test; p40 secretion is restricted to hCebpa/hSpi1.

To place every cell relative to the three possible identities (Fig. 6), we derived mutually exclusive fibroblast, adipocyte and myeloid gene sets and scored each cell against each signature (Fig. 7b). Two trajectories emerged, gaining adipogenic or myeloid score as fibroblast score fell, and only the fibroblast, mesenchyme and cycling clusters drew from all three arms (Supplementary Fig. 7b). Conversion required loss of the starting identity: *Snai1*, *Meox1* and *Scx* were silenced and *Twist1*, *Hmga2* and *Fosb* downregulated in every reprogrammed state (Supplementary Fig. 7c).

The adipogenic signature (95 genes; Fig. 7b,c) comprised *Pparg*, *Ppargc1a*, *Cd36*, *Fabp4*, *Dgat1*/*Dgat2*, *Agpat2* and *Acsl1*, with the recently nominated adipogenic factor *Sox30*^68^ and the thermogenic genes *Ucp1* and *Dio2* (Supplementary Fig. 7e). The myeloid signature (83 genes; Fig. 7b,c) comprised *Spi1*, *Irf8*, *Ciita*, *Lmo2* and *Nr1h3* as transcriptional core, *Fcer1g*, *Itgam*, *Csf1r*, *Tyrobp* and *Ptprc* as receptor machinery, and *Cd74*, *H2-Aa*, *H2-Ab1* and *H2-Eb1* as a complete MHC class II module (Supplementary Fig. 7f). Their ontologies were non-overlapping. The adipogenic set enriched for triglyceride and lipid biosynthesis, regulation of lipolysis in adipocytes and brown fat cell differentiation; the myeloid set for phagocytosis, antigen processing and presentation via MHC class II, regulation of cell activation and leukocyte activation (Fig. 7c). The two signatures mapped to largely separate functions, so cells scoring high on either signature had acquired the metabolic or immune function of that identity rather than its markers alone.

### Cebpα and SPI1 generate distinct myeloid outcomes and a dual-identity adipocyte– macrophage state

Integrating RNA with a 37-marker ADT panel spanning hematopoietic stem and progenitor, mature lineage, and mesenchymal epitopes assigned 38 transcriptome clusters to 16 discrete states, annotated by marker expression and immunophenotype (Fig. 7d,e, Supplementary Fig. 7b,g). Twenty markers were detected above background; T, B, erythroid and megakaryocyte epitopes were not, restricting output to the myeloid and adipogenic lineages.

Each reprogramming cocktail generated its own endpoint. Cebpα alone passed through a transient Sca-1⁺CD34⁺CD140a⁺ progenitor (*Dcn*, *Sfrp1*, *Grem2*) to the production of Fap⁺ activated adipogenic progenitor (*Fap*, *Epha7*, *Robo1*, *Zbtb20*, *Dio2*), which lost CD34 and CD140a. SPI1 alone gave a cKit⁺ progenitor and an antigen-presenting macrophage-like endpoint (*Igfbp2*, *Mfge8*, *Rbp1*), driven by JAK–STAT and interferon signaling (*Stat1*, *Stat6*, *Irf7*, *Ifnar1*, *Ifit2*/*Ifit3*). The dual Cebpα/SPI1 condition produced the Pparg⁺ adipocyte–macrophage hybrid at the branch point (Supplementary Fig. 7h) and, from it, an early adipocyte population (*Ryr2*, *Anxa1*, *Serpinb1b*) and a separate inflammatory macrophage population (*Apoe*, *Saa3*, *Colec12*, *Ccr5*, *Marco*), alongside a cycling myeloid-like state (*Spp1*, *S100a4*, *Cenpa*). All myeloid-marked states were CD45⁺Mac-1⁺, with the macrophage endpoints carrying CLEC12A⁺CD38⁺ surface proteins. The hybrid carried CD45 and Mac-1 markers while retaining *Pparg*, *Fabp4*, *Cd36* and *Retn*, so the same cell score high for both lineages on surface protein and transcript independently (Supplementary Fig. 7g). Proliferation was equally informative: *Mki67* and *Top2a* persisted only in unconverted fibroblasts, in the cycling myeloid-like state, and in the hybrid, while every terminal state was negative (Supplementary Fig. 7d).

Gene ontology confirmed the hybrid’s dual identity and separated the two adipogenic outputs (Fig. 7f, Supplementary Fig. 7i). The Pparg⁺ adipo-mac hybrid enriched simultaneously for lipid storage, peroxisome, cold-induced thermogenesis, and for antigen processing and presentation of endogenous antigen. The two adipogenic endpoints recovered differed in composition: the Cebpα-alone arm was dominated by the Fap⁺ progenitor, enriched for chondrocyte differentiation on *Sox5*, *Sox6*, *Runx2* and *Bmpr1a*/*1b*/*2*, whereas the dual condition yielded a terminal early adipocyte enriched for branched-chain amino-acid and sphingolipid catabolism on *Acat1*, *Hmgcll1* and *Asah1*. Since droplet capture under-recovers mature, lipid-laden adipocytes, this likely reflects the sampled compartment rather than the endpoint reached. Cebpα alone produces terminal adipocyte-like cells by both morphology and bulk transcriptome (Fig. 1, Fig. 3).

The myeloid endpoints differed reciprocally. The SPI1 state carried no lipid term, whereas the dual inflammatory macrophage was enriched for myeloid leukocyte activation on *Tlr2*, *Ccr5,* and *Il6ra,* as well as for triglyceride metabolism on *Apoe*, *C3,* and *Sorl1*. Its higher CD16/32 and IL-12p40 secretion, detected only with both factors and without a rise in p70 (Fig. 7g and Supplementary Fig. 7j), places it in a more activated state. Finally, the cycling myeloid state arose only with SPI1, and since a transiently proliferative intermediate accounts for elite reprogramming in B cells^20, 21^ it is a candidate for the efficiency of these conversions.

### Cebpα and SPI1 build discrete chromatin landscapes that combine additively

To define the chromatin underlying these cell types, we profiled the same three conditions at day 10 by scATAC-seq (30,454 cells). The recovered populations matched the CITE-seq annotation, establishing that each factor combination builds its own discrete set of accessible states (Fig. 8a,b, Supplementary Fig. 8a). ChromHMM^58^ resolved 20 chromatin states grouping into constitutive, fibroblast-, adipogenic-, pan-myeloid-and myeloid-accessible compartments, the latter four tracking the clusters of the condition that generated them (Fig. 8b,c, Supplementary Fig. 8a,b). Fibroblast-accessible states were lost in the terminal clusters of both arms, suggesting that the reprogramming processes dismantle the fibroblast epigenome rather than building over it (Fig. 8b,c, Supplementary Fig. 8a,b) in agreement with transcriptional data (Fig. 4,7).

**Fig. 8.**
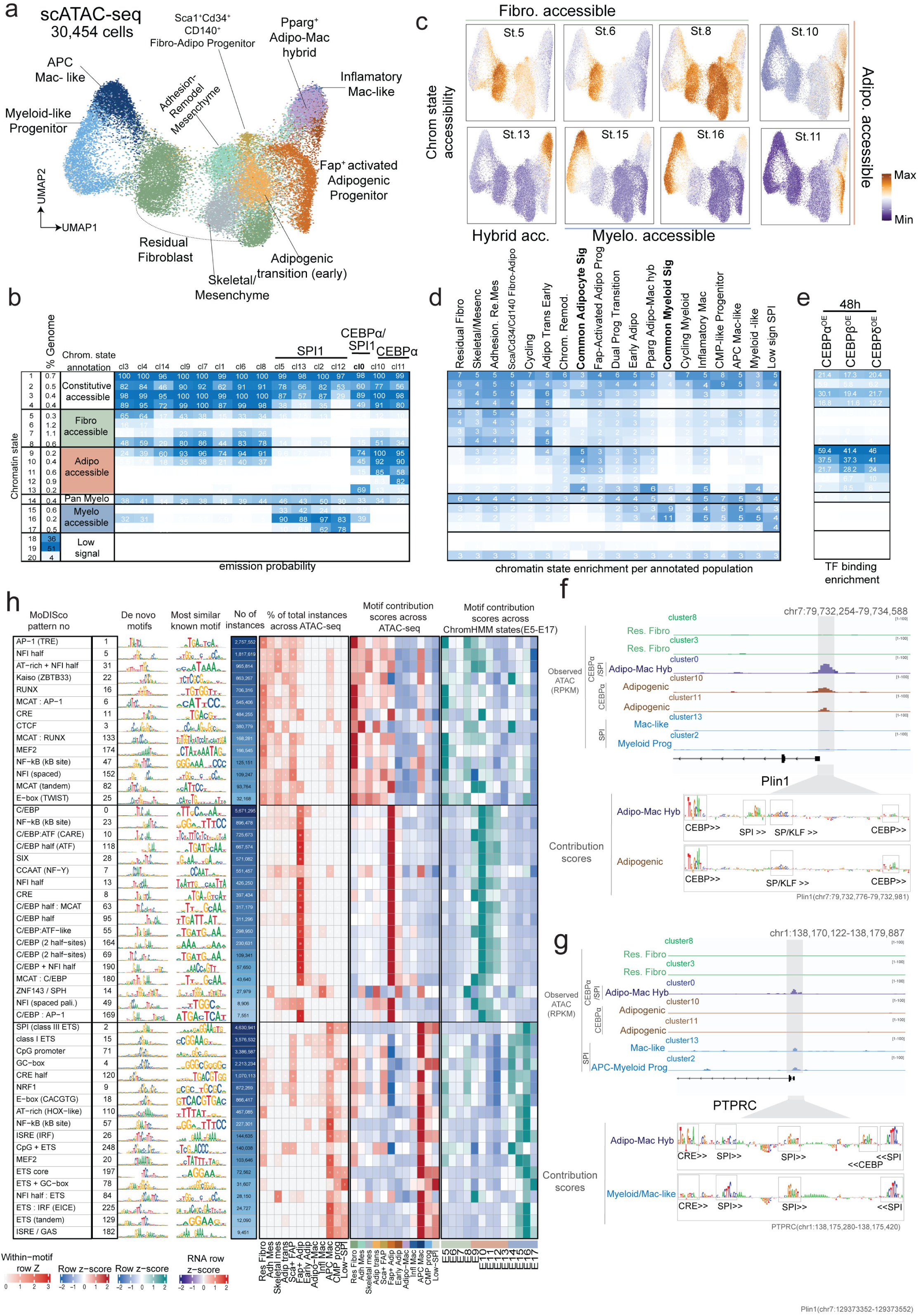
Cebpa and SPI1 encode separate adipogenic and myeloid chromatin grammars that coexist in a Pparg^+^ adipo-mac hybrid. **a**, scATAC-seq of MEFs transduced with hCebpα, hSPI1 or hCebpα/hSPI1 at day 10 (30,454 cells); UMAP coloured by annotated population. The recovered populations mirror the CITE-seq annotation in Fig. 7d. **b**, ChromHMM segmentation of the scATAC data into 20 states. Left, genome fraction per state; right, emission probability of accessibility across the 15 scATAC clusters (cl0–cl14), grouped as constitutive accessible (1–4), fibroblast accessible (5–8), adipogenic accessible (9–13), pan-myeloid (14), myeloid accessible (15–17) and low signal (18–20). Clusters ordered by dominant condition; cl0 (bold) is specific to hCebpα/hSPI1. **c**, Accessibility of representative chromatin states projected on the scATAC UMAP: fibroblast-accessible states 5, 6 and 8; hybrid-accessible state 13; myeloid-accessible states 15 and 16; adipogenic-accessible states 10 and 11 (right); Color, log_2_(RPKM+1). **d**, Enrichment of each chromatin state within ±20 kb of the transcription start sites of the genes defining each population signature (Fig. 7e) and the common adipocyte and common myeloid signatures (Fig. 7c; bold). **e**, Enrichment of Cebpα, Cebpβ and Cebpδ occupancy per chromatin state (48 h ChIP–seq, Fig. S4a-d). **f**, Observed accessibility (RPKM, scale 1–100) at the *Plin1* locus (chr7:79,732,254–79,734,588) in clusters associated with residual fibroblasts, Pparg⁺ adipo–mac hybrid, adipogenic, and myeloid/macrophage-like populations. Below, ChromBPNet per-base contribution scores over the highlighted promoter window (chr7:79,732,776– 79,732,981) for the hybrid and adipogenic clusters; contributing motif calls boxed and oriented. Two C/EBP elements and an SP/KLF element are called in both; while a SPI element contributes only in the hybrid. **g**, As in f, at the *Ptprc* locus (chr1:138,170,122–138,179,887), zoom window chr1:138,175,280–138,175,420, contribution scores for the hybrid and myeloid/macrophage-like populations. A CRE element and three SPI elements are called in both, whereas a C/EBP element is called only in the hybrid. **h**, ChromBPNet motif contribution across the scATAC data. Rows, 50 de novo TF-MoDISco motifs representing each chromatin compartment, grouped into shared/fibroblast-, C/EBP-and ETS-weighted blocks; left to right, element name, MoDISco pattern number, de novo contribution weight matrix, most similar known motif, total genomic instances, within-motif row Z of the percentage of population specific locations having each motif across the 12 annotated populations (numbers, percentage; color, row Z-score); motif importance (FiNeMo hit importance per Mb) across the same populations (row Z-score); and motif importance across ChromHMM states States 5 to State 17 (row Z-score).

Chromatin remodeling was cell-type-specific and organized around cell-type specific gene signatures. Scoring each state within ±20 kb of the TSS of the genes defining each population placed the adipogenic-accessible states around the adipogenic signatures and the myeloid-accessible states around the myeloid ones (Fig. 8b,d). Each reprogramming arm therefore opens its own set of regulatory regions to drive its defining gene program. For the adipogenic arm, this reconfiguration traced back to Cebp: Cebpα, Cebpβ and Cebpδ bound these regions within 48 h of factor induction (Fig. 8e, Supplementary Fig. 8c), nucleating accessibility that persisted through day 10. Cebp on its own opened no myeloid chromatin.

The dual condition opened both epigenetic landscapes at once. hCebpα/hSPI1 cells formed a chromatin state of their own (cl0; Supplementary Fig. 8a), in which the adipogenic-and myeloid-accessible sites were open together, a property no population from either single-factor arm displayed (Fig. 8b). At individual loci this state carried the parental grammar plus one element of the opposite arm, and accessibility tracked transcript. *Plin1* was accessible only in the Cebpα adipogenic clusters and the dual state, its promoter read through two C/EBP elements flanking an SP/KLF element, with an SPI element contributing only in the dual state, matching its induction in the adipocyte program. *Ptprc* was accessible and transcriptionally active only in the SPI1 myeloid clusters and the dual state, its promoter read through a CRE and three SPI elements, with C/EBP contributing exclusively in the hybrid cell state (Fig. 7e, 8f,g).

What the two loci showed at single promoters held across the genome. Ranking every de novo element by its contribution to each population separated the same three compartments (Fig. 8h). Residual fibroblasts scored highest for RUNX, MEF2, TWIST E-box and AP-1, and contribution at these elements declined in the early adipocyte, Pparg⁺ hybrid and inflammatory macrophage-like states, in parallel with the downregulation of the corresponding factors (Supplementary Fig. 7c). The adipogenic states were dominated by C/EBP in its single, half-site and two-half-site forms, with SIX, NFI half and CCAAT (NF-Y); the myeloid states by class III SPI, class I ETS, ISRE (IRF) and NF-κB, the κB element being the only one positive in the inflammatory macrophage-like state. Both compartments also carried composites, C/EBP:ATF (CARE), C/EBP + NFI half and MCAT:C/EBP on the adipogenic side, ETS:IRF (EICE) and NFI half:ETS on the myeloid side. The partners fused into these elements, ATF, NFI, NF-Y and MCAT, are the same factors recruited beside Cebp as separate motifs at de novo sites at 48 h (Fig. 5c), suggesting that partners engaged independently during initiation are later read as single composite elements. Each arm therefore converges on a grammar of one shared anchor and a compartment-specific partner.

## DISCUSSION

We delivered one factor into four cells and asked what each cell contributed. Cebp gave four outcomes, and one predicted partner installed or abolished a fate in a single starting cell. Identity, not potency, determined the outcome: fibroblasts and neural stem cells both differentiated into fat, embryonic stem cells gained trophectoderm characteristics, and B cells converted into macrophages. A Cebp factor carries no specific instruction of its own. It assembles one at the DNA from three ingredients: its concentration, the partners the cell supplies, and the grammar it must displace (Supplementary Fig. 8d).

Concentration determines how many sites a factor can reach, while the partners a cell expresses determine which of them it can use. Raising Cebp did not change the sequence it reads: the same contribution weight matrices underlie constitutive and de novo sites, which differ in motif copy number. Regions carrying several copies are occupied at basal factor levels, while single-copy regions cross threshold only as the factor accumulates. The partner contribution was legible in two forms, at two stages. First, the cis-regulatory elements that Cebp opened in each starting cell carried a distinct set of partner motifs alongside the Cebp motif, NFI, NF-Y and AP-1/ATF in fibroblasts and neural stem cells, SPI1 in B cells, TFAP2C in embryonic stem cells, so the elements grouped the four systems by outcome. Second, as conversion progressed, the same partners reappeared as composites in which the two motifs were fused at fixed spacing: Cebp with ATF and an NFI half-site in the adipogenic arm; SPI with IRF in the myeloid one.

Modeling which motifs support a peak, rather than whether it is open, showed that equivalent accessibility can be assembled in more than one way. The *Plin1* promoter was supported by C/EBP elements in the Cebpα arm and the *Ptprc* promoter by SPI elements in the SPI1 arm, while in the dual arm both promoters drew on both families, each factor contributing at its own motif rather than through a shared site^69^. The same accessible region can therefore be built by different combinations of factors in different conditions. This reading accounts for the hybrid. When supplied together, Cebpα and SPI1 did not fuse into a single instruction: their contributions remained separate and additive, and a cell receiving both ran the adipogenic and myeloid networks in parallel. The *Pparg*⁺CD45⁺ adipocyte–macrophage state is the cell-level arrangement. The population appears transient, maximal at day 6, giving way by day 10 to the early adipocyte and inflammatory macrophage-like endpoints. The subsequent appearance of both endpoints argues that this coexistence is resolved rather than maintained.

Both single-cell atlases contained proliferative intermediates: a cycling myeloid-like state and a cycling *Pparg*⁺ hybrid, each arising only where SPI1 was supplied. The same proliferative stage has been read in opposite ways. In leukaemia, Cebpα with PU.1 converts blasts into antigen-presenting cells and lowers Ki-67, but conversion is incomplete and the residual cycling fraction limits disease control^70^. In B cells, the proliferative intermediate produced by a Cebpα pulse plausibly makes them elite substrates for OSKM^20, 21^. Our data place both in one frame: Cebpα with a myeloid partner generates a proliferative intermediate as a normal stage of conversion, permissive for what follows rather than deciding it, and whether it reads as an obstacle or an advantage depends on what the cell is asked to do next.

The reprogramming factors are not exempt from this logic. We previously showed OSKM relocating the fibroblast’s own factors as it engages the genome; here the traffic reverses. Elevated Cebp drew Oct4, Sox2 and Klf4 onto its own regions, and pluripotency failed not because it was repressed but because its factors were elsewhere. This also settles the route taken. Fibroblasts expressing OSKM with Cebp became adipocytes within three to five days, before iPSC colonies arise, and pluripotent cells given Cebpα made trophectoderm rather than fat, so the conversion is direct transdifferentiation and not a passage through pluripotency followed by redifferentiation. The two accounts then converge on one prediction: supplying SPI1 to the fibroblast arm should partition Cebp away from the adipogenic elements that hold OSK, thereby releasing those factors while retaining the proliferative intermediate and thereby permitting rapid colony emergence.

Context-dependent transcription factor function is therefore programmable: assembled from partners rather than fixed in the protein, readable from the grammar a cell exposes and alterable one factor at a time.

## Methods

### Contact for Reagent and Resource Sharing

Please direct any requests for further information or reagents to the lead contact, Dr Constantinos Chronis, Department of Biochemistry and Molecular Genetics, University of Illinois Chicago, Chicago, IL 60607, USA.

### Experimental Model and Subject Details

Mice harboring the R26-M2rtTA allele and either a single dox-inducible polycistronic cassette coding for OKSM in the Col1A locus (tetO-OSKM)^71^ or a dox-inducible cassette coding for OSK (tetO-OKSmCherry) in the Col1A locus, were used for reprogramming, ChIP-seq, ATAC-seq and expression (RNA-seq) assays. All animal procedures were approved by the Institutional Animal Care and Use Committee (Animal Care Committee) of the University of Illinois Chicago under protocols 20-227 and 23177 and were performed in accordance with institutional and federal guidelines. Embryos of both sexes were used and were not genotyped for sex.

### Primary cell isolation and culture

Mouse embryonic fibroblasts (MEF). Primary MEFs were isolated from the husks of E13.5 embryos of the tetO-OSKM, tetO-OSKmCherry reprogrammable mouse model, ROSA26-M2rtTA, or C57BL/6 after removal of the head and internal organs. Embryos of both sexes were used and were not genotyped for sex. MEFs were maintained in fibroblast medium composed of DMEM, 10% fetal bovine serum (FBS), 50 U ml−1 penicillin/streptomycin, 1x GlutaMAX, 1x non-essential amino acids, and β-mercaptoethanol, and were passaged at sub-confluence.

Neural stem cells (NSCs). NSCs were isolated from the brains of E13.5 embryos of the reprogrammable mouse model and cultured as neurospheres on non-treated culture dishes in NSC medium containing 50% Neurobasal medium, 50% DMEM/F12, 1x N-2 Supplement, 1x GlutaMAX, 1x penicillin/streptomycin, 10 ng/ml basic fibroblast growth factor (bFGF), and 20 ng/ml epidermal growth factor (EGF). Neurospheres were expanded, dissociated with TrypLE Express Enzyme (Gibco), and seeded onto Matrigel (Corning)-coated dishes for culture as an adherent monolayer. Isolated NSCs were validated by immunostaining for SOX2, Vimentin, SOX1, and NeuroD1 (see Immunofluorescence). Transgene expression was induced by adding doxycycline (2 µg/ml) to the culture medium.

Pro-B cells. Bone marrow cells were isolated from the femurs, tibias, and humeri of tetO-OSKM mice by flushing marrow with PBS. Cleaned bones were placed cut-end down in perforated PCR tubes nested in 1.7 mL centrifuge tubes and marrow was recovered by centrifugation at 10,000g for 1 min. Cells were washed twice with HBSS (5 min, 1500 rpm, 4 °C), resuspended in 50 µL cold HBSS. Red blood cells (RBCs) were lysed in 150 µL lysis buffer (BD Biosciences) for approximately 1.5 mins at room temperature. RBC lysis was quenched with 5 mL column buffer (HBSS, 0.5% FBS, 2mM EDTA). Cells were washed in HBSS and counted. B-lineage cells were enriched by magnetic-activated cell sorting (MACS) using positive selection for CD19. Briefly, cells were resuspended in column buffer at 10⁶ cells/mL and Fc receptors were blocked at 0.1 µg per 10⁶ cells for 10 min on ice. Cells were then resuspended in column buffer at 4 µL per 10⁶ cells and incubated with biotin-conjugated anti-CD19 at 0.25 µg per 10⁶ cells for 30 min on ice. After washing, cells were resuspended in 90 µL of column buffer plus 10 µL of streptavidin microbeads per 10⁶ cells and incubated for 30 min on ice with occasional agitation. Cells were washed with 1 ml column buffer per 10⁶ cells, resuspended in 500 µL column buffer and applied to an LS Column mounted on a QuadroMACS Separator (Miltenyi Biotech). The column was washed with 3 × 3 mL cold column buffer to remove the CD19⁻ fraction. The column was then removed from the magnet and the retained CD19⁺ cells were eluted with 5 ml cold column buffer. CD19 expression was verified via flow cytometry. Pro-B cells were cultured at 1–1.3 x 10⁶ cells/mL in Opti-MEM supplemented with 20% FBS, 1x penicillin-streptomycin, 1x GlutaMAX, β-ME and 10 ng/mL IL-7 (BioLegend, 577804) at 37°C in a humidified 5% CO₂ incubator. IL-7 was replenished every 1–2 days and cells were expanded for at least one week before reprogramming experiments. Transgene expression was induced by adding doxycycline (2 µg/ml) to the culture medium.

Published RNA-seq, ATAC-seq and C/EBPα ChIP-seq data for CEBPα-pulsed B cells were obtained from GEO accession GSE71218. Chromatin marks H3K27ac, H3K27me3, H3K36me3, H3K4me1, H3K4me2, H3K4me3, H3K9ac, H3K9me3 and H3K79me2 of isolated and expanded pro-B cells were profiled by Cut-N-Run.

### Plasmids and inducible transgene constructs and cell lines

Inducible ESC lines were generated by FLPe-mediated cassette exchange at the Col1a1 locus,^72^ of KH2 ESCs. Insert sequences for 3×FLAG-hCEBPA-E2A-ZsGreen and for ZsGreen alone were amplified by inverse PCR with Phusion High-Fidelity PCR Master Mix (Thermo Fisher). The pBS31 exchange vector, which carries a tetracycline-responsive promoter, a 5′ FRT site and a promoterless, ATG-less selection element, was linearized with EcoRI and the inserts introduced by In-Fusion cloning (Takara), giving pBS31-3×FLAG-hCEBPA-E2A-ZsGreen1 (KCPL195) and the pBS31-ZsGreen control (KCPL325). Plasmids were propagated in E. coli, purified (ZymoPURE II Plasmid Maxiprep Kit) and verified by Sanger sequencing across the insert and both FRT junctions. KH2 ESCs carry the M2rtTA transactivator in the Rosa26 locus and a single FRT homing cassette downstream of the Col1a1 3′ UTR. FLPe-mediated recombination at the FRT site inserts the cassette as a single copy in the defined Col1a1 configuration and simultaneously reconstitutes the selection marker, so that only correctly targeted clones survive selection in hygromycin B. Resistant clones were picked, expanded on irradiated MEF feeders and validated by doxycycline-dependent ZsGreen fluorescence, and by 3×FLAG immunofluorescence, confirming nuclear localization of the induced protein. Transgene expression was induced with doxycycline (2 µg/ml). Control lines carrying the ZsGreen1-only cassette were generated in parallel and handled identically.

### Viral production and transduction

Retroviruses (pMX vectors) and lentiviruses were produced by transfection of Platinum-E or HEK293T with the transfer vector and appropriate packaging/envelope plasmids. Viral supernatants were collected at 48 and 72 hours, filtered, and concentrated. Target MEFs and NSCs were transduced twice on consecutive days in the presence of polybrene.

### Reprogramming and fate-conversion assays

Cebpα/β adipogenic-potential assay in MEFs. C57BL/6 MEFs were transduced twice with pMX-Cebpa-FLAG (KCPL63), pMX-Cebpb-FLAG (KCPL60), or empty pMX (KCPL53). Cells were cultured in mESC medium (FBS/LIF) for the first two days and switched from day 3 onward to KO DMEM medium in which FBS was replaced with 15% knockout serum replacement (Gibco, 10828028) supplemented with 0.1 µg/ml recombinant leukemia inhibitory factor (LIF) (KSR/LIF). Lipid accumulation was assayed by Oil Red O (ORO) staining at day 10, and ORO-positive events were quantified across microscopic fields as described under Statistics and reproducibility.

Concurrent Cebp overexpression and OSKM-mediated MEF reprogramming. Reprogrammable MEFs were transduced with pMX-Cebpa-FLAG or pMX-Cebpb-FLAG (or empty pMX), and OSKM was induced with doxycycline (2 µg/ml). Conditions included OSKM-only, OSKM + Cebpα, OSKM + Cebpβ, and empty-vector controls. Cells were cultured in MEF media for two days and in KSR/LIF from day 3. Readouts included ORO staining (day 6), Nanog immunofluorescence to score reprogramming efficiency (day 10), and LipidTOX/FLAG/DAPI immunofluorescence (day 6). Nanog-positive colonies were defined as clusters of five or more Nanog-positive cells and were quantified manually in ImageJ.

Time course imaging of hCEBP-induced adipocyte-like emergence. Reprogrammable MEFs were co-transduced with inducible ZsGreen-linked hCEBPA (KCPL178 or KCPL437) or hCEBPB (KCPL179 or KCPL436) constructs, imaged in bright field and FITC via ECHO Revolution microscopy at 4X over a doxycycline induction time course. Representative single cells were annotated on day 6 and traced backward through the series using TVL1 optical-flow lineage tracing. Scale bar, 100 µm.

Adipocyte doxycycline-induction withdrawal assay. Doxycycline was added at day 0 and withdrawn after defined intervals, alongside a continuously induced control; cultures were collected at day 10 and terminal adipocyte differentiation was read out by LipidTOX quantification.

Human CEBP-family screen in MEFs. Reprogrammable MEFs were lentivirally transduced with individual tetO-3xFLAG-hCEBP-ZsGreen constructs (hCEBPA, KCPL178; hCEBPB, KCPL179; hCEBPD, KCPL180; hCEBPE, KCPL181; hCEBPZ, KCPL182) or a ZsGreen control (KCPL24) and cultured under OSKM reprogramming conditions with doxycycline (2 µg/ml). Factor expression and localization were validated by FLAG/ZsGreen immunofluorescence (confocal, 20x) antibodies. ORO staining (day 6 or day 10, as appropriate) and Nanog immunofluorescence (day 10) were quantified as described.

OSK plus Cebp reprogramming assay. Reprogrammable MEFs carrying Col1a1::tetO-OSKmCherry; ROSA26::M2rtTA were transduced with inducible hCEBPA (KCPL437) or hCEBPB (KCPL436) constructs and cultured with doxycycline to co-induce OSK and CEBP. OSK expression was validated by Oct4/Sox2/Klf4 antibodies. and ORO-positive events were quantified.

B-cell reprogramming-medium comparison. Reprogrammable MEFs transduced with pMX-Cebpa-FLAG or pMX-Cebpb-FLAG were cultured in FBS/LIF for two days and then switched to N2B27+2iL B-cell reprogramming medium^20^: 50% Neurobasal medium, 50% DMEM/F12, 1x N-2 Supplement, 1x B-27 Supplement, 1x GlutaMAX, 1x penicillin/streptomycin, 1x NEAA, β-ME, 0.1 µg/ml recombinant LIF, and the small-molecule inhibitors 0.4 µM PD0325901 and 3 µM CHIR99021, with concurrent OSKM induction. ORO-positive events were quantified as described.

Cebpα induction in ESCs. Doxycycline-inducible Cebpα-ZsGreen mESCs and control ESCs were treated with doxycycline (2 µg/ml) and analyzed by immunofluorescence at 24, 48, and 72 h post-induction and by RNA-seq at 24 and 48 h post-induction.

ESC morphology and lipid assessment. ESC colony morphology following Cebpα induction was imaged by confocal microscopy (LSM 900). ZsGreen, LipidTOX, and DAPI staining were used to determine whether ESCs underwent lipid accumulation/adipogenic conversion.

Trophectoderm-marker immunofluorescence. Cebpα-induced ESCs were immunostained for trophectoderm-associated markers EOMES, GATA3, KRT7, KRT8, TP63, and TFAP2C, together with ZsGreen and DAPI antibodies.

Cebpα/β adipogenic-potential assay in NSCs (Fig. 2G–K). NSCs from E13.5 embryos were transduced twice with pMX-Cebpa-FLAG (KCPL63), pMX-Cebpb-FLAG (KCPL60), or empty pMX (KCPL53) and assayed by ORO staining at day 10. LipidTOX/FLAG/DAPI immunofluorescence was performed at day 6 using antibodies and the overlap between LipidTOX-positive and FLAG-positive cells was quantified.

Concurrent Cebp overexpression and OSKM-mediated NSC reprogramming. Reprogrammable NSCs were transduced with pMX-Cebpa-FLAG (KCPL63) or pMX-Cebpb-FLAG(KCPL60) and induced with doxycycline (2 µg/ml) to activate OSKM, with parallel no-doxycycline controls. NSCs were cultured in NSC medium and switched after six days of expression to N2B27+2iL medium for the remainder of reprogramming. Cells were profiled at day 6 and day 12. Readouts included ORO staining (day 6) and Nanog immunofluorescence (day 10, where applicable), with ORO-positive events quantified as described.

Concurrent Cebpα overexpression and Spi1-mediated MEF reprogramming. CD45-CD11b-CD44-Ly6G-CD48-R26-M2rtTA MEFs were sorted on a Sony MA900 multi-application cell sorter using FACS. MEFS were transduced twice with hCEBPA (KCPL437), hSPI1 (KCPL291), or the bicistronic hCEBPA-E2A-hSPI1 (KCPL547) and induced with doxycycline (2 µg/ml). Cells were profiled at day 2 and day 10. FLAG/SPI1/DAPI immunofluorescence was performed at day 2, ORO staining and the overlap between LipidTOX-positive/CD11b-positive FLAG-positive SPI1-positive cells was quantified at day 10, along with immunofluorescence to characterize LipidTOX and CD11b-positive CD45-positive cell populations at the end point of reprogramming using antibodies. CITE-seq was performed at days 3, 6 and 10 as described.

Myeloid cell doxycycline-induction withdrawal assay. Doxycycline was added at day 0 and withdrawn after defined intervals, alongside a continuously induced control; cultures were collected at day 10 and terminal myeloid conversion was read out by CD11b quantification.

Concurrent Cebpα overexpression and Tfap2c-mediated MEF reprogramming. R26-M2rtTA MEFs were transduced twice with SFFV-3×Flag-hCebpα-E2A-hTfap2c-ZsGreen (KCPL559). MEFs were cultured in reprogramming media for 10 days. Cells were profiled at day 2 and day 10. FLAG/TFAP2C/DAPI immunofluorescence was performed at day 2 using antibodies. and ORO-staining at day 10, with ORO-positive events quantified as described.

Time course imaging of hCEBP-hTFAP2C induced MEFs. Reprogrammable MEFs were transduced twice with SFFV-3×Flag-hCebpα-E2A-hTfap2c-ZsGreen (KCPL559). Cells were imaged in bright field and FITC via ECHO Revolution microscopy at 4X over the induction time course. Representative single cells were annotated at day 6 and tracked backward through the series by TVL1 optical-flow lineage tracing. Scale bar, 100 µm.

### Immunofluorescence

Cells were grown on glass coverslips precoated with 0.2% porcine skin gelatin (Sigma G2500; for MEFs and most stains) or Matrigel (for NSCs). Cells were fixed with 4% paraformaldehyde for 10 min, permeabilized with 0.1–0.5% Triton X-100 in PBS, and blocked in blocking buffer (5–10% normal donkey serum) Primary antibodies, with host species, supplier, catalogue number, clone and working dilution, were applied in blocking buffer overnight at 4°C; species-matched secondary antibodies were applied in blocking buffer for 30–60 min at room temperature in the dark. Coverslips were washed three times with 1x PBS-Tween between incubations, with one wash containing DAPI (100 ng/ml) to label nuclei and were mounted with Aqua-Poly/Mount (Polysciences 18606).

### Oil Red O staining and quantification

A 0.5% (w/v) Oil Red O (ORO; Sigma O0625) stock was prepared in 100% isopropanol. A 1x working solution was made by mixing 3 parts stock with 2 parts ddH2O and filtering through a 0.22-µm PVDF syringe filter. Cells on gelatin-coated coverslips were fixed in 4% paraformaldehyde for 10 min, washed twice in ddH2O, incubated in 60% isopropanol for 1 min, and stained with 1x ORO working solution for 15–20 min at room temperature. Coverslips were washed thoroughly with ddH2O and mounted with Aqua-Poly/Mount (Polysciences 18606). Whole-coverslip images were acquired at 4x on an Echo Revolve microscope using mosaic tiling; high-resolution ORO images were taken on a widefield Leica DMi8 microscope at 40x. Lipid droplets/ORO-positive events were counted manually in ImageJ using the multi-point tool across five fields from three biological replicates.

### LipidTOX staining and quantification

Neutral lipid droplets were visualized with HCS LipidTOX Neutral Lipid Stain (Thermo Fisher, H34475; Thermo Fischer, H34477). Cells were fixed with 4% paraformaldehyde and permeabilized with 0.1% Triton X-100 in PBS. Cells were stained with LipidTOX working solution (1:200 in PBS) for 30 min at room temperature To identify CEBP-expressing cells, samples were immunostained with anti-FLAG (Sigma F1804, 1:200) overnight at 4°C followed by secondary antibody as described and mounted with Aqua-Poly/Mount. Imaging was performed by confocal microscopy on a Zeiss LSM900 (20x). Quantification was performed by determining the percentage of cells that were both FLAG-positive and LipidTOX-positive.

### Microscopy and image analysis

Brightfield images of reprogramming cultures were acquired at 4x on an Echo Revolve microscope. Whole-well/stitched imaging for Nanog and ORO assays used the Echo Revolve mosaic feature; high-resolution ORO images used a widefield Leica DMi8 (40x). Immunofluorescence was imaged by confocal microscopy on a Zeiss LSM900 (20x). Image quantification was performed in ImageJ/Fiji.

### Fluorescence-activated cell sorting (FACS)

R26-M2rtTA MEFs were sorted on a Sony MA900 Multi-Application Cell sorter using FACS. CD45^-^ CD11b^-^ CD44^-^ and Ly6G^-^ cells were collected using the following marker–fluorochrome–clone combinations: CD45 (FITC, 30-F11), CD11b (Alexa Fluor 488, M1/70), CD44 (FITC, IM7), Ly6G (APC), and CD48 (APC, HM48-1), with DAPI for viability. Sorted populations were collected by centrifugation (1000 rpm, 5 min), resuspended in MEF media, and plated for reprogramming experiments.

### Flow cytometry

Multiparameter flow cytometry of the myeloid/monocyte–macrophage–neutrophil compartment was conducted on a BD FACSCelesta Flow cytometer using the following marker–fluorochrome–clone combinations: CD45 (Super Bright 780, 30-F11), CD11b (PE-Cy7 / APC-Cy7 / PE / APC-eFluor 780, M1/70), Ly6G (PE, 1A8), Ly6C (APC-eFluor 780, HK1.4), F4/80 (APC, BM8), CD64 (PerCP-Cy5.5, X54-5/7.1), CD11c (PE-Cy7, N418), MHC-II (BV421, M5/114.15.2), CD68 (PerCP-eFluor 710, FA-11), CD86 (APC-eFluor 780, GL1), CD115/CSF1R (PE-Cy7, AFS98), CD18 (PE, M18/2), and CD16/32 (FcγRII/III; PerCP-Cy5.5, clone 93) (see Immunofluorescence) with DAPI or propidium iodide (PI) for viability.

### LEGENDplex™ assay (cytokine protein level analysis)

R26-M2rtTA MEFs transduced with hCEBPA (KCPL437), hSPI1 (KCPL291), hCEBPA-E2A-hSPI1 (KCPL547), and untransduced MEF controls were cultured in doxycycline-containing reprogramming media in a six well plate for ten days. Doxycycline-containing media was refreshed into standard reprogramming media 24 hours prior to media collection. Media was collected and the cytokine expression was profiled using a mouse 13-plex bead-based LEGENDplex® assay panel (LEGENDplex® Mouse Macrophage/Microglia Standard Cocktail, according to manufacturer’s instructions (740852, BioLegend). Mouse proinflammatory cytokines were profiled via beads targeting CXCL1, TGF-β1, IL-18, IL-23, CCL22, IL-10, IL-12p70, IL-6, TNF-α, G-CSF, CCL17, IL-12p40, IL-1β. Beads were analyzed using a CytoFLEX flow cytometer and Biolegend Qognit software. Only IL-12p40 was within detectable range.

### RNA-seq

RNA was isolated using the Direct-zol RNA Miniprep Kit (Zymo) with on-column DNase treatment. mRNA was selected and strand-specific libraries were prepared with the TruSeq Stranded mRNA Kit (Illumina) per manufacturer instructions. Libraries were sequenced on Illumina NovaSeq 6000 or NovaSeq X Plus platforms.

### ATAC-seq

ATAC-seq was performed as described^73^. 100,000 cells were resuspended in 100 µl lysis buffer (10 mM Tris-HCl, 10 mM NaCl, 3 mM MgCl2, 0.1% Tween-20, 0.1% NP-40, 0.01% digitonin, 1% BSA), incubated 5 min at 4°C, and washed (10 mM Tris-HCl, 10 mM NaCl, 3 mM MgCl2, 0.1% Tween-20, 1% BSA). Nuclei were pelleted (500 × g, 5 min, 4°C) and resuspended in 50 µl transposition mix (22.5 µl water, 2.5 µl Tagment DNA Enzyme 1 [Illumina], 25 µl 2x Tagment DNA Buffer [Illumina]). Transposition proceeded for 30 min at 37°C, and DNA was purified with the DNA Clean & Concentrator Kit (Zymo). Transposed DNA was amplified in two sequential PCR reactions (7 then 5 cycles); after the first PCR, libraries were size-selected on a 2% low-melt agarose gel for 150–700-bp fragments using the Zymoclean Gel DNA Recovery Kit (Zymo), and second-PCR libraries were purified identically. Libraries were sequenced on Illumina NovaSeq 6000 or NovaSeq X Plus.

### CUT&Run

CUT&Run^74^ was performed following the EpiCypher CUTANA CUT&RUN Protocol v2.0 with minor modifications. 250,000 cells were resuspended in Wash Buffer (20 mM HEPES-pH 7.5, 150 mM NaCl, 0.5 mM spermidine, Roche cOmplete EDTA-free Protease Inhibitor). Concanavalin-A beads (11 µl/sample; Polysciences) were activated in Bead Activation Buffer (20 mM HEPES-pH 7.9, 10 mM KCl, 1 mM CaCl2, 1 mM MnCl2) and bound to cells for 10 min at room temperature. Bead-bound cells were resuspended in 50 µl antibody buffer (Wash Buffer + 0.01% digitonin, 2 mM EDTA) and incubated overnight at 4°C (650 rpm) with 0.5 µg antibody per sample: H3K27ac (Active Motif 39133), H3K4me3 (Abcam ab1012), H3K4me2 (Epicypher 13-0027), H3K9ac (Invitrogen MA5-33384), H3K36me3 (Abcam ab9050), H3K27me3 (EMD Millipore 07-449), or CTCF (EpiCypher 13-2014). Samples were washed twice in Digitonin Buffer (Wash Buffer + 0.01% digitonin), resuspended in 50 µl Digitonin Buffer with 2.5 µl 20x pAG-MNase (EpiCypher), and incubated 10 min at room temperature. After two washes, chromatin was digested in 50 µl Digitonin Buffer + 1 µl 100 mM CaCl2 (2 h, 4°C, 650 rpm), and the reaction was stopped with 33 µl Stop Buffer (340 mM NaCl, 20 mM EDTA, 4 mM EGTA, 50 µg/ml RNase A, 50 µg/ml glycogen). Fragments were released (10 min, 37°C) and purified by 1.5x AMPure bead selection.

### Cross-linked ChIP-seq

Endogenous CEBPβ occupancy in uninduced MEFs (the 100 and 010 binding groups, Fig. 4a) was obtained from Chronis et al. (2017). X-ChIP was performed as described (Chronis et al., 2017) for FLAG (Sigma F1804), Oct4 (RnD AF1959), Sox2 (RnD AF2018), Klf4 (RnD AF3158), H3K27ac (Active Motif 39133), H3K4me1 (Millipore 07-436), H3K79me2 (EMD Millipore 04-835), and H3K79me1 (Active Motif 39145). Cells were cross-linked in 1% formaldehyde (10 min, RT), quenched with 0.125 M glycine, resuspended in Swelling Buffer (25 mM HEPES, 1.5 mM MgCl2, 10 mM KCl, 0.1% NP-40, 1 mM DTT, Roche protease inhibitor), and incubated 10 min on ice. Nuclei were pelleted (1000 rpm, 5 min, 4°C), resuspended in Sonication Buffer (50 mM HEPES, 140 mM NaCl, 1 mM EDTA, 1% Triton X-100, 0.1% Na-deoxycholate, 0.2% SDS), and sonicated on a Diagenode Bioruptor (48 cycles, 30 s ON/45 s OFF). 5 µg antibody-conjugated beads were incubated overnight at 4°C with 10 µg sonicated chromatin in RIPA buffer (50 mM Tris-HCl pH 8.0, 150 mM NaCl, 2 mM EDTA, 1% NP-40, 0.1% Na-deoxycholate, 0.1% SDS). Beads were washed twice each in Low-Salt, High-Salt, LiCl, and TE buffers, and DNA was recovered by overnight reverse cross-linking at 55°C (20 mg/ml proteinase K, 1% SDS) followed by phenol-chloroform-isoamyl alcohol extraction.

Library preparation and sequencing (CUT&Run and X-ChIP). DNA from both methods was prepared into TruSeq-compatible libraries using the NEBNext Ultra II End Repair/dA-Tailing Module (NEB) and ligated to Illumina dual-index adaptors with the NEBNext Ultra II Ligation Module (NEB). Reactions were purified with AMPure beads and amplified in two sequential PCR steps (9 then 7 cycles). Libraries were sequenced on Illumina NovaSeq 6000 or NovaSeq X Plus.

### Bulk RNA-seq processing

Paired-end RNA-seq reads of 50, 100 or 150 bp were trimmed with Trim Galore in paired-end mode, and FastQC reports were generated after trimming. Trimmed reads were aligned to the mouse GRCm39 genome with STAR v2.7.10b^75^ in two-pass mode using 10 threads. GENCODE^76^ mouse vM33 was used for gene annotation. Read-length-specific STAR indices were used with sjdbOverhang values of 49, 99 or 149 for 50, 100 or 150 bp reads, respectively. Alignments were output as coordinate-sorted BAM files, and gene-level counts were generated with STAR --quantMode GeneCounts. Alignments with noncanonical intron motifs were filtered using --outFilterIntronMotifs RemoveNoncanonical. Gene counts from the appropriate strandedness column of STAR GeneCounts were used for downstream analysis.

Differential expression analysis was performed in R with DESeq2 using a design formula of ∼ condition. Size-factor-normalized counts and variance-stabilizing transformed (VST) values were generated with DESeq2. Principal component analysis was performed on the 500 most variable genes. Pairwise contrasts were tested across all conditions, and log2 fold changes were shrunk using ashr^77, 78^. Genes with a Benjamini–Hochberg-adjusted P value ≤ 0.05, absolute log2 fold change ≥ 1 and baseMean ≥ 10 were considered differentially expressed. The union of differentially expressed genes across all contrasts was used for clustering and visualization.

For heatmaps, VST values were averaged across biological replicates within each condition and z-scored by gene across conditions. Differentially expressed genes were grouped by k-means clustering. TPM values were calculated from raw counts using GRCm39 GENCODE mouse vM33 gene lengths, averaged across biological replicates, transformed as log2(TPM + 1), and visualized by cluster using violin. Pairwise differences in gene-level TPM distributions between conditions within each cluster were assessed using two-sided Wilcoxon rank-sum tests with Benjamini–Hochberg correction implemented in rstatix.

### ATAC-seq and ChIP-seq processing

Raw paired-end FASTQ files generated across multiple lanes or sequencing runs were concatenated by sample and read direction before alignment. Reads were aligned to the mouse mm10 genome using Bowtie2 ^5^ with vendor quality filtering enabled (--qc-filter). SAM files were converted to BAM format using SAMtools, followed by coordinate sorting and indexing. PCR duplicates were removed with Picard ^6^ MarkDuplicates using REMOVE_DUPLICATES=true, and duplication metrics were retained for quality-control assessment.

For ATAC-seq, CPM-normalized bigWig tracks were generated from deduplicated BAM files using deepTools ^7^ bamCoverage with --normalizeUsing CPM, --binSize 25, --extendReads 200, --centerReads. Peaks were called with MACS2 callpeak using -g mm and -q 0.01 (and with -p 0.01 for relaxed peaks); paired-end samples were analyzed in BAMPE mode using -f BAMPE. For ChIP-seq, CPM-normalized bigWig tracks were generated from deduplicated BAM files using deepTools bamCoverage with --normalizeUsing CPM, --binSize 25 and --extendReads. Signal tracks and called peaks were inspected in IGV.

To generate the heatmaps shown in Figures 4a, 4b, 4c, 4j, S4b, S4c, S4d, S4g, S4m, and S4n, we followed the approach described in Chronis et al.^5^. Briefly, genomic features of interest, such as transcription factor peaks, were aligned at their summits, which were provided as summit-centered BED intervals. The flanking regions upstream and downstream of each summit were tiled within ±2 kb using 100 bp bins. For each region, sequencing reads from the corresponding sorted and deduplicated BAM file were counted across each 100 bp bin, excluding unmapped reads. Read counts were normalized to FPKM-like values using the total number of mapped reads and bin size. No input or control sample normalization was applied. The resulting binned signal matrices were exported in CDT format and visualized as heatmaps using Java TreeView^79^ Metaplots were generated by averaging the normalized signal value for each 100 bp bin across all regions in the corresponding feature set.

### Classification of CEBPβ binding events across MEF reprogramming conditions

We generated seven sets of CEBPβ binding events across three conditions: MEFs, 48 h OSKM-induced MEFs, and 48 h OSKM + CEBPβ-induced MEFs. MACS2 summit intervals were compared using BEDTools. To allow nearby summits to be considered overlapping, summit intervals were expanded using slopBed with the mm10 chromosome size file.

Condition-specific binding events (100, 010, and 001) were identified by expanding summit intervals by ±250 bp and removing intervals that overlapped either of the other two conditions using subtractBed -A. Pairwise shared binding events (110, 011, and 101) were identified by intersecting summit intervals expanded by ±100 bp between the two relevant conditions, followed by removal of intervals overlapping the third condition within ±250 bp. Binding events shared across all three conditions (111) were identified by sequential intersection of ±100 bp expanded summit intervals from all three conditions.

Original summit coordinates were recovered using intersectBed -u, producing seven mutually exclusive BED files corresponding to the binary overlap groups 100, 010, 001, 110, 011, 101, and 111, where each digit indicates the presence (1) or absence (0) of a CEBPβ binding event in MEFs, 48 h OSKM-induced MEFs, and 48 h OSKM + CEBPβ-induced MEFs, respectively.

CEBPβ binding groups were further filtered to retain binding events with ChIP-seq and ATAC-seq signal patterns consistent with their assigned binary labels. For each binding event, signal was quantified across a 4 kb region centered on the group summit, spanning 2 kb upstream and 2 kb downstream, and summed separately for each condition. Binding events in condition-specific groups (100, 010, and 001) were retained only when the expected positive condition showed the highest total signal. Binding events in pairwise shared groups (110, 101, and 011) were retained only when both expected positive conditions showed higher total signal than the excluded condition.

### Classification of overexpressed CEBPα/β binding events across cell types

For Fig. S5b, the same BEDTools-based seven-group classification strategy was applied to ChIP-seq summit sets from 48 h OSKM + C/EBPβ-induced MEFs, 48 h CEBPα-induced ESCs, and 18 h B cells expressing hCEBPα alone (B′). The binary group labels indicate the presence (1) or absence (0) of binding in these three datasets, respectively.

### Classification of OSKM-bound accessible regions during early ipsc reprogramming

OSKM-bound regions were defined by generating genome-wide 500 bp windows across the mm10 genome and retaining windows that overlapped at least one MACS2 summit from 48 h Oct4, Sox2, Klf4, or c-Myc ChIP-seq. Windows overlapping any of the four factors were merged to generate a union set of OSKM-bound windows.

To classify these regions by accessibility status, MEF and 48 h OSKM-induced MEF ATAC-seq summits within OSKM-bound windows were compared using BEDTools. Regions were assigned to three groups: MEF-accessible only (10), 48 h OSKM-accessible only (01), or accessible in both conditions (11). Condition-specific groups were defined by excluding summits overlapping the other condition within ±250 bp, whereas shared regions were defined by overlap between MEF and 48 h OSKM ATAC-seq summits within ±100 bp.

### ChromHMM analysis

Chromatin-state segmentations were generated using ChromHMM v1.27^58^ across the analyzed cell types and experimental conditions. Separate ChromHMM models were generated for histone-mark datasets and ATAC-seq profiles. Reads were binarized in non-overlapping 200-bp genomic bins using BinarizeBam with the -mixed option.

To derive histone-based chromatin-state segmentations across the four starting cell types, MEF, NSC, ESC and B cell, we used nine histone marks in a stacked ChromHMM strategy, following the approach described in Chronis et al^5^. Briefly, we constructed a single virtual cell type in which datasets from the four starting cell types were represented as individual marks, with each input label encoding both the source cell type and histone-mark name. ChromHMM LearnModel was then used to learn chromatin-state models and annotate the genome in this virtual cell type. Models with 5–40 states were evaluated, and a 22-state model was selected for downstream analyses based on interpretability and its ability to capture combinatorial histone-mark patterns across cell types.

For ATAC-seq ChromHMM modeling, the same stacked strategy was applied to ATAC-seq BAM files from MEF, NSC, ESC and B cell, together with their CEBPA-and/or CEBPB-overexpression conditions, comprising 20 conditions in total. Each ATAC-seq dataset was represented as an individual mark in a single virtual cell type, with labels encoding the source cell type and experimental condition. Models with 5–40 states were evaluated, and a 23-state model was selected for downstream analyses because it provided an interpretable representation of open and closed chromatin patterns across cell types and perturbation conditions.

### Preparation of ChromBPNet input regions

ChromBPNet^28^ input regions were generated from processed ATAC-seq BAM files for 20 mouse cell-state and reprogramming conditions: MEF, MEF-derived Day2^OSKM^, MEF-derived D2^OSKM+A^, MEF-derived D2^OSKM+B^, MEF-derived D2^OSKM+hA^, MEF-derived D2^+A^, MEF-derived D2^+B^, NSC, NSC-derived D2^OSKM+A^, NSC-derived D2^OSKM+A^, primary B cells, ESCs, ESCs + hCEBPα (Day1, Day2), and B cells expressing CEBPα alone (B^α^) or with OSKM for 1 (D1^OSKM+A^) or 2 days (D2^OSKM+A^). The mm10 no-alt ENCODE analysis-set FASTA, mm10 chromosome sizes file and mm10 ENCODE blacklist file ENCFF547MET were used throughout preprocessing. BAM files were sorted, indexed and filtered with SAMtools using samtools view -b -F 780 -q 30. Relaxed ATAC-seq peaks were called independently for each sample with MACS2 using callpeak -f BAM -g mm -p 0.01 --shift -75 --extsize 150 --nomodel --keep-dup all --call-summits. Peaks were ranked by MACS2 signalValue, and the top 200,000 peaks per sample were retained.

Blacklist filtering was performed after extending the mm10 blacklist by 1,057 bp on each side with BEDTools slop, corresponding to one half of the 2,114-bp ChromBPNet input window. Peaks overlapping the expanded blacklist were removed with bedtools intersect -v. For each sample and chromosome split, GC-matched background regions were generated with chrombpnet prep nonpeaks using the mm10 FASTA, chromosome sizes file, blacklist BED file, blacklist-filtered peak set and fold-specific chromosome split JSON. Default ChromBPNet background-region settings were used^28^.

Models used fivefold chromosome hold-out cross-validation across mm10 chromosomes 1-19, X and Y. Each fold assigned non-overlapping chromosomes to training, validation and test sets, and each chromosome was included in the test set in at least one fold. Fold compositions are available at: https://mitra.stanford.edu/kundaje/oak/zhangby/mouse-atlas/splits/.

### ChromBPNet model training and interpretation

ChromBPNet models were trained separately for each sample and fold in ATAC mode. ChromBPNet uses 2,114-bp one-hot encoded DNA sequence from peak and matched background regions to predict both the base-resolution accessibility profile and total natural log accessibility counts over the central 1,000 bp.

Tn5 bias models were trained with chrombpnet bias pipeline for each sample, fold and bias-threshold factor. Threshold factors of 0.5, 0.6 and 0.7 were evaluated. Bias models were interpreted with DeepLIFT^28^ and inspected to select the threshold factor that captured Tn5-associated sequence preference without strong transcription-factor motif signal. The selected bias threshold factor was 0.5 for all MEF conditions and untreated B cells, 0.6 for all NSC samples and ESC D1^+hCEBPA^, and 0.7 for ESC, ESC D2^+hCEBPA^, B cells expressing CEBPα alone (B^α^), B cell D1^OSKM+A^ and B cell D2^OSKM+A^. In total, the bias-model selection stage evaluated 300 bias models across 20 samples, three threshold factors and five folds.

Full ChromBPNet models were trained with chrombpnet pipeline using the filtered BAM file, blacklist-filtered top 200,000 peaks, fold-specific GC-matched background regions, mm10 genome FASTA, chromosome sizes file, fold-specific split file and selected pre-trained bias model. This produced 100 full ChromBPNet models across 20 samples and five folds.

Model training was run on Intel GPUs of ALCF Aurora compute nodes using the system TensorFlow backend from the Aurora frameworks module. ChromBPNet was installed from the official GitHub repository in a Python virtual environment created with system site packages enabled; TensorFlow entries in the ChromBPNet requirements file were omitted to preserve compatibility with the Aurora TensorFlow installation. Aurora jobs were managed with Parsl as ensembles of single-node tasks. Bias-model training used one task per sample–fold–parameter combination, while full ChromBPNet training used one task per sample–fold combination.

ChromBPNet interpretation used chrombpnet contribs_bw to compute DeepLIFT ^28^ contribution scores for each nucleotide in each 2,114-bp input sequence with respect to the predicted counts output. Fold-specific count contribution scores were generated for all peak regions from the bias-corrected ChromBPNet component (chrombpnet_nobias.h5) and then averaged across folds. Fold-specific DeepSHAP HDF5 files were averaged across the five chromosome hold-out models for each sample, and averaged contribution scores were converted to bigWig format. Predicted accessibility tracks were generated with chrombpnet pred_bw from the bias-corrected ChromBPNet model for each fold and then averaged across folds.

### De novo motif discovery

De novo motif discovery was performed independently for each ATAC-seq condition using TF-MoDISco (https://github.com/jmschrei/tfmodisco-lite/) on the averaged ChromBPNet count contribution scores. TF-MoDISco identifies high-importance seqlets and clusters them into recurrent contribution-weighted sequence patterns. Motifs were discovered with modisco motifs using up to 5,000,000 seqlets (-n 5000000), a minimum metacluster size of 2 (-l 2), and a 1,000-bp search window (-w 1000). Motif reports were generated with modisco report and annotated against a curated MEME-format JASPAR^80^ motif database.

### Assembly of the CEBP reprogramming motif lexicon

To define a non-redundant motif lexicon across CEBP-driven reprogramming conditions, we adapted the consolidation strategy^47^. TF-MoDISco was applied independently to each ATAC-seq condition to identify recurrent high-contribution seqlets and assemble them into de novo motifs. Positive TF-MoDISco patterns from the count models were used for the primary motif lexicon. For each of 20 conditions, TF-MoDISco seqlets were linked to the corresponding one-hot sequence and nucleotide contribution-score tracks from the averaged interpretation files. Interpreted regions were centered on peak summits, and motif merging was performed within the central 1,000 bp.

Similar motifs were collapsed across conditions using a modified TF-MoDISco-lite SimilarPatternsCollapser that preserved seqlet provenance, including sample label, original peak index, seqlet coordinates and strand. Motif pairs were aligned using Pearson correlation on magnitude-transformed contribution scores with a minimum fractional overlap of 0.7. Motifs with more than 300 seqlets were subsampled to 300 seqlets for pairwise similarity estimation. Motif pairs were merged iteratively using TF-MoDISco-lite merge thresholds: (0.8, 0.8), (0.5, 0.85) or (0.2, 0.9), with dealbreaker thresholds of (0.4, 0.75), (0.2, 0.8), (0.1, 0.85) and (0.0, 0.9).

After each merge, motifs were trimmed to the region supported by at least 20% of seqlets and at least 30 seqlets. Merged motifs were expanded by 5 bp on each side, trimmed to the 20-bp window with the highest information content relative to the empirical nucleotide background, and expanded again by 5 bp. Final merged motifs were stored in TF-MoDISco HDF5 format with the position probability matrix, contribution-weight matrix, hypothetical contribution matrix and seqlet-level metadata.

### Known-motif annotation and merged-lexicon motif-instance calling

Known-motif annotation of the merged lexicon was performed with TOMTOM^81^ against the curated MEME-format JASPAR^15^ motif database. For each merged motif, the contribution-weight matrix was trimmed to positions with total absolute contribution at least 30% of the maximum positional contribution. Trimmed contribution-weight matrices were converted to position probability matrices by retaining positive contribution values, adding a pseudocount of 1 x 10^-6 and normalizing each position to sum to one. TOMTOM was run with -no-ssc -text -min-overlap 5 -mi 1 -dist kullback -evalue -thresh 10.0. For every motif, we manually inspected the most similar PWMs to assign a provisional label, typically at the transcription factor family level. This resulted in a set of 443 non-redundant consolidated motifs (unique ID is a value from 0 to 442) used for downstream analysis.

Predictive motif instances for the merged lexicon were identified with FiNeMo (https://githun.com/austintwang/finemo_gpu), which performs motif scanning in a contribution-aware manner. FiNeMo was run with finemo call-hits --global-lambda 0.7 -b 200 –compile on merged motif HDF5 file and merged NPZ contribution-score input.

### Single-cell library preparation and sequencing

Cells were dissociated using the Animal Component-Free Cell Dissociation Kit (STEMCELL Technologies, 05426) according to the manufacturer’s instructions. Cells were counted and resuspended in Cell Staining Buffer (BioLegend, 420201), blocked with TruStain FcX (BioLegend, 101320) for 10 min on ice and stained with TotalSeq-A anti-mouse hashtag oligonucleotides (HTOs) and a cocktail of 37 antibody-derived tag (ADT) conjugated antibodies for 30 min on ice. Cells were washed three times with Cell Staining Buffer, counted, pooled and encapsulated using the Chromium GEM-X Single Cell 3′ v4 Gene Expression kit (10x Genomics). cDNA amplification, purification and construction of gene expression and feature-barcode libraries were performed according to the manufacturer’s protocol.

For the single cell ATAC experiment, cells were dissociated using the same procedure. Following dissociation, cells were permeabilized with ice cold lysis buffer containing 10 mM Tris HCl, 10 mM NaCl, 3 mM MgCl2, 0.1% Tween 20, 0.1% Nonidet P40 Substitute, 0.01% Digitonin, 1% BSA, 1 mM DTT and 1 U/ul RNase inhibitor for 5 min on ice. Cells were then washed with wash buffer containing 10 mM Tris HCl, 10 mM NaCl, 3 mM MgCl2, 1% BSA, 0.1% Tween 20, 1 mM DTT and 1 U/ul RNase inhibitor. Tagmentation, GEM incubation and library construction were performed according to the Chromium Next GEM Single Cell Multiome ATAC + Gene Expression user guide. Although the Multiome chemistry was used for library preparation, the gene expression component from this experiment was not used for downstream biological interpretation, and these data were analyzed as single cell ATAC.

CITE-seq gene expression, ADT and HTO libraries were pooled, quantified by qPCR and sequenced on a 25B lane of a NovaSeq X Plus with V1.0 sequencing kits using paired-end reads (Read 1, 28nt; Read 2, 150nt, i7 and i5 index reads, 10nt), and gene expression libraries were additionally re-sequenced on a second 25B lane with the same configuration. Multiome ATAC libraries were sequenced on a 25B lane of a NovaSeq X Plus using paired end reads (Read 1, 100nt; Read 2, 100nt; i7 sample index, 8nt; i5 cell barcode, 24nt). Sequencing output was converted to FASTQ files with bcl-convert (Illumina; v4.3.6 and v4.4 for the CITE-seq libraries) and used as input for Cell Ranger. FASTQ files from gene expression, ADT and HTO libraries were processed with Cell Ranger v9.0.1. Reads were aligned to the 10x Genomics mouse reference ‘refdata-gex-mm10-2020-A’ using ‘cellranger count’ with a custom TotalSeq-A mouse feature reference containing 37 ADT features and sample timepoint HTOs. Cell Ranger outputs from the RNA sample libraries were aggregated with ‘cellranger aggr’ using ‘--normalize=none’. The aggregated filtered feature barcode matrix was used for downstream analysis, with gene expression counts loaded as the RNA assay and antibody capture counts loaded as the raw ADT assay in R with Seurat^82^.

Single-cell ATAC libraries for each sample was processed independently with Cell Ranger ARC v2.0.2 using ‘cellranger-arc count’ and the 10x Genomics mouse multiome reference ‘refdata-cellranger-arc-mm10-2020-A-2.0.0’. The resulting Cell Ranger ARC outputs from the individual ATAC libraries were then combined with ‘cellranger-arc aggr’ using the same reference and ‘--normalize=none’. The aggregated Cell Ranger ARC output was used for downstream single-cell ATAC analysis.

### HTO Calling and Quality Control for CITE-seq Analysis

Cells were multiplexed with HTOs to distinguish sample timepoints within each sample. HTO barcode count matrices were generated with CITE-seq-Count v1.4.5 (doi: 10.5281/zenodo.2590196, https://github.com/Hoohm/CITE-seq-Count/releases) using detected barcodes from Cell Ranger as a whitelist. For each sample, paired HTO FASTQ files were counted against a mouse TotalSeq-A HTO tag list. The HTO count matrices were then imported into Seurat and added into the multimodal object. For each cell, the HTO with the highest count was used to assign the collection timepoint. Cells that could not be assigned to a valid timepoint annotation were not included in the downstream multimodal analysis.

Quality control (QC) was performed separately for each sample. Mitochondrial fraction was calculated for each cell and ambient RNA contamination was estimated with decontX^83^. Cells were removed using scater’s ‘isOutlier’ function via the thresholds on log-transformed count distributions^84^, with low RNA UMI count defined as more than 3 median absolute deviations below the sample median, high RNA UMI count as more than 2 median absolute deviations above the sample median, and low or high ADT counts as more than 3 median absolute deviations from the sample median. Cells were also removed if mitochondrial content exceeded 20% and if the decontX contamination estimate exceeded 0.5. After this initial QC step, doublets were identified with scDblFinder^85^ using sample identity as the grouping variable, and only cells classified as singlets were kept for the downstream analysis.

To further reduce ambient background signal beyond decontX-based filtering, raw Cell Ranger feature barcode matrices were processed with CellBender ‘remove-background’^86^. CellBender was run independently for each sequencing libraries using the corresponding ‘raw_feature_bc_matrix.h5’ files as input. CellBender corrected gene expression and ADT matrices were imported into the final Seurat object as separate RNA and ADT assays, with RNA and ADT features restricted to those within the filtered object. CellBender corrected RNA counts were log-normalized, and CellBender corrected ADT counts were centered log-ratio (CLR) normalized using the cell-wise geometric mean procedure described below. We used CellBender corrected values for all downstream expression analysis such as trajectory and signature scoring, marker heatmaps except noted otherwise.

### Multimodal Analysis of CITE-seq

RNA counts were normalized with SCTransform^87^ in Seurat, regressing out total RNA UMI count and mitochondrial fraction. ADT counts were normalized by centered log-ratio (CLR) transformation. Briefly, a pseudocount of 1 was added to each ADT count, the geometric mean across ADTs was calculated separately for each cell, and each ADT value was divided by the corresponding cell-specific geometric mean before log transformation. CLR-normalized ADT values were stored as a separate assay. The 37 ADT features were scaled and used for principal component analysis.

A joint RNA-ADT representation was generated using Seurat’s weighted nearest-neighbor (WNN) approach. RNA PCA dimensions 1-50 and ADT PCA dimensions 1-20 were used to construct the multimodal nearest-neighbor graph. Clusters were identified from the WNN shared nearest-neighbor graph, and WNN UMAP was used as the primary embedding for visualization and downstream annotation.

Cell state associated gene signatures and selected marker genes were evaluated using CellBender-corrected, log normalized RNA expression and visualized on the WNN UMAP. MEF-like, adipocyte-like and myeloid-like scores were calculated as described in the Trajectory Score Analysis.

For the single-cell heatmap shown in Figure 7e, population-associated gene signatures were selected using CellBender-corrected RNA expression. Mean log1p-normalized expression across all cells within each population was used to evaluate gene assignments. Genes were assigned to individual populations based on their expression patterns, while genes showing low variation across adipocyte-like or myeloid-like populations were assigned to common adipocyte and myeloid signatures, respectively. The heatmap included 2,739 genes across 9,791 selected cells. RNA expression was scaled to gene-wise z-scores across the selected cells, and CLR-normalized ADT expression was scaled separately. Genes were grouped by their final population signatures and displayed in the assigned order, with representative genes labeled.

Gene ontology (GO) enrichment analysis of the gene lists was performed using an online annotation suite Metascape^88^ (http://metascape.org) where the statistical significance was calculated using the cumulative hypergeometric test and terms with a P value below 0.01 were considered significantly enriched.

### Trajectory Score Analysis

To quantify progression of cells along putative MEF-like-to-adipocyte-like and MEF-like-to-myeloid-like trajectories, we calculated gene module-based directional trajectory scores from CellBender-corrected RNA expression. Curated marker gene sets associated with MEF-like (GS_MEF_), adipocyte-like (GS_Adipo_) and myeloid-like (GS_myeloid_) populations were used to calculate the trajectory scores. For each cell *c* and gene set GS_k_ where k ∈ {MEF, Adipo, Myeloid}, the module score was calculated as the mean log-normalized expression of the genes in the gene set:

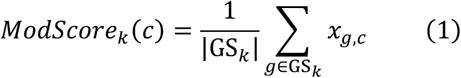

where *x_g,c_* is the log-normalized expression of gene g in cell c, and |GSk| is the number of genes in the set detected in the dataset. Module scores were scaled to the [0,1] range using the 1^st^ and 99^th^ percentiles across all cells with values outside this range is clipped to 0 or 1. For a score *v* with value *v_c_* in cell c:

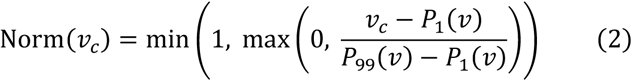

where *P*_1_(*v*) and *P*_99_(*v*) are the 1^st^ and the 99^th^ percentiles of *v* across all cells, giving us the normalized module scores:

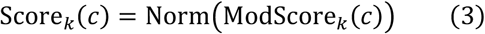

Directional trajectory scores were then calculated by subtracting the MEF-like score from the adipocyte-like or myeloid-like score and normalizing again with the same percentile-based approach:

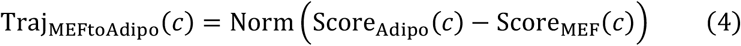

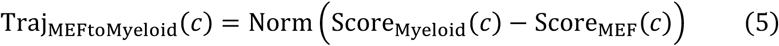

Trajectory scores therefore range from 0 to 1, with lower values indicating a more MEF-like transcriptional state and higher values indicating the progression toward the adipocyte-like or the myeloid-like state. The resulting scores then were plotted on the WNN UMAP for visualization.

### Single-cell ATAC quality control and clustering

Single-cell libraries were prepared using multiome chemistry but were analyzed as single-cell ATAC data with Signac^89^. RNA-derived measurements were used only for initial cell quality control and were not included in downstream biological analysis. ATAC fragments were used to calculate nucleosome signal and transcription start site enrichment, and ambient RNA contamination was estimated with decontX. Quality control was performed separately for each sample using scater isOutlier. Cells with RNA counts more than 3 median absolute deviations below or 2 median absolute deviations above the sample median were removed. Cells with ATAC counts more than 3 median absolute deviations below or 1.5 median absolute deviations above the sample median were removed. Cells with TSS enrichment more than 4 median absolute deviations below or above the sample median were also removed. Cells were additionally excluded when mitochondrial content exceeded 25% or the decontX contamination estimate exceeded 0.5. Doublets were identified with scDblFinder, and only cells classified as singlets were retained.

Following quality control, peaks were called from retained ATAC cells using MACS2 through Signac CallPeaks. Peaks on nonstandard chromosomes were removed, and a new cell-by-peak matrix was generated from the ATAC fragment files using FeatureMatrix. Peak accessibility was normalized with term frequency–inverse document frequency transformation, followed by selection of top accessible features and singular value decomposition. ATAC UMAP coordinates were calculated from latent semantic indexing (LSI) dimensions 2 to 50. A nearest-neighbor graph was constructed from LSI dimensions 2 to 50, and cells were clustered using Seurat graph-based clustering with algorithm 3 at a resolution of 1.2. These ATAC clusters were used as the starting point for chromatin-state modeling and population annotation.

### Single-cell ATAC ChromHMM Analysis

For each ATAC cluster at resolution 1.2, barcodes were grouped by cluster, and cluster-specific BAM files were generated from the Cell Ranger ATAC BAM files using sinto (https://github.com/timoast/sinto). These BAM files were used to generate a 20-state ChromHMM^58^ model. The same cluster-specific BAM files were also used to generate RPKM-normalized accessibility tracks at the Plin1 and Ptprc loci using deepTools bamCoverage. ChromHMM states were interpreted based on their accessibility patterns across the ATAC clusters, identifying states associated with adipogenic, myeloid, mixed adipogenic–myeloid, and MEF-like chromatin programs. For each ATAC cluster, mean accessibility scores were calculated for the different ChromHMM programs and converted to z-scores across clusters. A ChromHMM-associated program was assigned when its z-score was at least 0.5 and exceeded the second-highest program score by at least 0.10. The mixed adipogenic–myeloid program was separated into adipogenic-and myeloid-associated branches based on their relative accessibility in the corresponding lineage-associated states. Sample composition was also considered when assigning the final ATAC population labels, which were used for downstream ChromBPNet analysis.

ChromHMM state accessibility was assessed at the single-cell level by intersecting the 20-state ChromHMM segmentation with ATAC-seq fragments. Fragment counts were calculated for each genomic interval in each cell using Signac FeatureMatrix. For each state, fragments were summed across all genomic intervals assigned to that state and normalized by the total number of fragments per cell and the total genomic length of the state to get a RPKM-normalized accessibility score. Scores were transformed as log2(1 + RPKM) and visualized for each ChromHMM state on the ATAC UMAP.

To determine whether ChromHMM states were associated with CEBP occupancy, we calculated overlap enrichment using ChromHMM OverlapEnrichment. We calculated the fold enrichment of the seven previously defined CEBPβ binding groups from Fig. 4a within each ChromHMM state relative to their genome-wide representation. Enrichment of CEBPA, CEBPB and CEBPD ChIP-seq peaks was also calculated for each state to compare CEBP-family occupancy across the different chromatin states.

We also examined the relationship between ChromHMM states and the population-associated gene signatures from the CITE-seq analysis. For each gene set, genomic intervals were generated by extending 20 kb upstream and downstream of the transcription start site. ChromHMM overlap enrichment was calculated for these regions to determine whether specific chromatin states were enriched around genes associated with each RNA population.

### Defining single-cell ATAC Populations

We matched CITE-seq cell identities to the ATAC dataset by directly pairing ATAC and RNA cells. For each sample, highly variable RNA genes were identified, and ATAC peaks were assigned to their nearest gene transcription start site. Peaks located 2–100 kb from the nearest transcription start site were retained, and the most variable accessible peaks for each gene were selected based on accessibility variation across ATAC LSI-derived pseudobulks. Accessibility across these peaks was summed to generate a gene-level accessibility matrix for each ATAC cell. The RNA expression and ATAC gene-accessibility matrices were then normalized and projected into a shared canonical correlation analysis (CCA) space. ATAC and RNA cells were directly paired within this shared space, and the identity of the paired RNA cell was transferred to the corresponding ATAC cell.

These RNA-supported assignments were used to refine the broad ATAC populations defined by ChromHMM and sample composition. For most populations, the predominant identity among directly paired RNA cells was used to assign the corresponding ATAC population. Low-score CEBPA and CEBPA+SPI1 groups were kept separate according to their original ATAC clusters and annotated based on their RNA-pairing profiles. Two myeloid-associated populations required additional refinement. The mixed CEBPA+SPI1 myeloid population was subclustered in ATAC LSI space and separated into Pparg+ Adipo-mac hybrid and Inflammatory Mac-like populations based on enrichment of the corresponding paired RNA identities. The SPI1 myeloid-like population was similarly separated into CMP-like progenitor and Low-signature-SPI1 populations using enrichment of these RNA identities within its existing ATAC subclusters. These assignments produced the final 12 RNA-matched ATAC populations used for downstream regulatory analyses.

### Single-cell ATAC population peak calling, and defining population associated locations

After final ATAC annotation, we separated fragments by the 12 final ATAC populations and pooled them to generate population-level pseudobulk fragment files. MACS3 peaks were called independently for each population using FRAG input, --nolambda, --max-count 1, a P-value threshold of 0.01, and summit calling. Regions overlapping the mm10 blacklist expanded by 1,057 bp were removed, and retained peaks were further filtered at a q-value threshold of 0.01.

To define population-specific regulatory locations, MACS3 peak summits were extended by 250 bp in both directions and compared across the 12 ATAC populations. Regions overlapping summit-centered regions from another population were considered shared and excluded from the population-specific sets. Overlapping kept intervals were then merged within each population to generate the final population-specific ATAC regulatory locations. For downstream ChromBPNet-based motif analyses, these locations were further restricted to regions overlapping genomic intervals interpreted by the original ChromBPNet model.

### Single-cell ATAC ChromBPNet Analysis

We used ChromBPNet^28^ to investigate the underlying sequence grammar contributing to accessibility across the single-cell ATAC populations. Model training and interpretation followed the same general ChromBPNet framework described above for the bulk ATAC analysis. However, models were generated from pseudobulk ATAC data corresponding to eight broad ChromHMM-based populations established before RNA-supported refinement of the final ATAC identities. These populations included two SPI1-associated myeloid populations, two MEF-like populations, two CEBPA-associated adipogenic populations, and two CEBPA+SPI1 populations representing an adipogenic branch and a myeloid branch. During final annotation, the two SPI1-associated myeloid populations were represented by an antigen-presenting myeloid-like population and a myeloid-like progenitor population, while the two MEF-like populations were represented by residual fibroblasts and the Sca1+Cd34+CD140a+ fibro-adipogenic progenitor population. The two CEBPA-associated adipogenic populations contributed to the Fap+ activated adipogenic progenitor population. Within the CEBPA+SPI1 populations, the adipogenic branch corresponded to the early-adipocyte population, while the myeloid branch was further resolved into inflammatory macrophage-like and Pparg+ adipocyte–macrophage hybrid populations. Thus, the ChromBPNet training populations did not map one-to-one to the final 12 ATAC populations. For ChromBPNet training, population-specific filtered BAM files and blacklist-filtered MACS3 peak sets were used.

As in the bulk analysis described above, we generated ChromBPNet contribution scores from the bias-corrected models and used them to identify sequence features contributing to predicted accessibility. We summarized de novo sequence patterns using TF-MoDISco^90^, merged motif patterns across the eight broad ATAC populations and compared them with known motif databases using TomTom, and used FiNeMo to identify predictive genomic instances of the merged motifs from the ChromBPNet contribution scores. Each FiNeMo instance was defined by its genomic position, motif identity, and hit importance.

For visualization, motif names were manually assigned by considering related transcription factor family members and selecting the most appropriate family-level label when the motif match was not specific to a single transcription factor.

To resolve these ChromBPNet-derived motif instances across the final 12 ATAC populations, we intersected FiNeMo motif instances with the population-specific regulatory locations described above. Duplicate assignments representing the same physical motif instance were collapsed, retaining the instance with the highest hit importance. For each motif and final ATAC population, we calculated the percentage of regulatory locations containing at least one motif instance as the number of distinct motif-positive locations divided by the total number of retained locations for that population and multiplied by 100. The values shown in the heatmap therefore represent the actual percentage of population-specific regulatory locations containing each motif. For visualization, the percentages were converted to z-scores separately for each motif across the 12 ATAC populations, and these row z-scores were used for the grey-to-red color scale. The color therefore indicates whether a motif is relatively more or less common in one population compared with the other populations for that same motif, while the value shown in each cell gives the actual percentage of regulatory locations containing the motif.

We also calculated motif importance using FiNeMo hit importance. For each motif and final ATAC population, hit importance was summed across kept motif instances, divided by the total kept genomic width for that population, and multiplied by 1,000,000 to obtain motif importance per Mb. These values were displayed as row z-scores across the 12 ATAC populations using a blue-to-white-to-red color scale centered at zero. FiNeMo motif instances were also assigned to ChromHMM states based on genomic overlap. For the final summary, chromatin states E5 through E17 were shown. Hit importance was summed for each motif and state, normalized by the retained genomic width of that state, and converted to motif importance per Mb. These values were also displayed as row z-scores using a blue-to-white-to-teal color scale centered at zero. Because both importance heatmaps were standardized separately for each motif, they show how the relative importance of a given motif changes across ATAC populations or ChromHMM states. Total FiNeMo instance columns were colored using log10(count + 1), while the values shown in the cells represent the corresponding instance counts.

The final motif summary included the curated motif identity, TF-MoDISco pattern number, de novo contribution-weight motif, most similar known motif, total FiNeMo instance counts, the percentage of population-specific regulatory locations containing each motif across the 12 ATAC populations, motif importance across those populations, and motif importance across ChromHMM states E5 through E17. Fifty representative motifs were ordered within each group by decreasing total FiNeMo instance count.

### Statistics and reproducibility

No statistical method was used to predetermine sample size. No data were excluded from the analyses. Experiments were not randomized, and investigators were not blinded to allocation during experiments and outcome assessment. Imaging and staining experiments were performed in n = 3 independent biological replicates, with at least 5 fields scored per replicate; ORO-positive events and Nanog-positive colonies were counted manually in ImageJ by [one/two] scorer(s) blinded to condition. Bulk RNA-seq was performed in at least n = 2 biological replicates per condition, ATAC-seq in at least n = 2, CUT&Run in n = 2, and X-ChIP in n = 2. For analysis replicates were merged to increase sequencing counts.

Data are presented as mean ± s.d. / mean ± s.e.m; the center value and error bars are defined in each figure legend, together with replicate number(s). Comparisons between two conditions were made with a two-tailed unpaired Student’s t-test / Mann–Whitney U-test; comparisons across more than two conditions with one-way ANOVA with Tukey’s post hoc test. For RNA-seq gene-cluster comparisons, log2(TPM + 1) distributions between conditions were compared using two-sided Wilcoxon rank-sum tests with Benjamini– Hochberg correction. Significant P values are reported in the figure legends. Statistical analyses were performed in R. For sequencing-based analyses, significance thresholds are given in the relevant processing sections; differential expression used a Benjamini–Hochberg-adjusted P ≤ 0.05 with |log2 fold change| ≥ 1 and baseMean ≥ 10, and GO enrichment used an accumulative hypergeometric test with P < 0.01.

For the single-cell analyses, CITE-seq and single-cell ATAC-seq were performed once per condition. The final CITE-seq atlas contained 110,751 cells (38,214 CEBPA, 35,436 CEBPA+SPI1 and 37,101 SPI1), and the final single-cell ATAC dataset contained 30,454 cells (12,219 CEBPA, 6,427 CEBPA+SPI1 and 11,808 SPI1) after quality control. Cell-level filtering and quality-control criteria are described in the relevant Methods sections.

For single-cell RNA visualization, selected population-associated and common-lineage signature genes were displayed using CellBender-corrected expression and row-scaled z scores. The final heatmap gene sets were not restricted solely to genes passing a single differential-expression significance threshold. Gene ontology and pathway enrichment of the selected gene sets was performed using Metascape, with significance assessed using the accumulative hypergeometric test and P < 0.01 considered significant.

For single-cell ATAC analyses, statistical thresholds and filtering criteria used for quality control, ChromHMM state assignment, peak calling, motif enrichment and ChromBPNet/FiNeMo analyses are described in the corresponding Methods sections.

### Cell lines

V6.5 mouse embryonic stem cells were a generous gift from Kathrin Plath. Platinum-E and HEK293T packaging cells were obtained from ATCC. Primary MEFs, NSCs, and pro-B cells were derived in-house from the mouse lines described above and were used at low passage. All lines were routinely tested for mycoplasma contamination by PCR methods and were negative. None of the lines used appears on the ICLAC register of commonly misidentified cell lines.

### Data availability

All sequencing data generated in this study, including bulk RNA-seq, ATAC-seq, CUT&Run, X-ChIP-seq, CITE-seq and single-cell ATAC-seq, and their processed downstream analysis files have been deposited in the Gene Expression Omnibus under SuperSeries accession **GSE344075**. Previously published datasets re-analyzed in this study are available from the accessions cited in the corresponding Methods sections. Additional processed data supporting the findings of this study will be made available through Zenodo upon publication. All other data are available from the corresponding author on reasonable request.

### RShiny app development

An interactive RShiny web application, CEBP Compass, was developed to explore the CITE-seq and scATAC-seq datasets and is available at https://cebpcompass.org/. The application was built with shiny (v1.13.0) and uses the final CITE-seq and scATAC-seq Seurat objects, which will be deposited in Zenodo upon publication. For the CITE-seq data, users can view sample, timepoint, WNN cluster annotations, final cell identities and MEF-to-Adipocyte-like and MEF-to-Myeloid-like trajectory scores on the WNN UMAP, and the CellBender-corrected RNA expression for any gene and CLR-normalized ADT expression alongside the corresponding RNA transcripts. For the scATAC-seq data, users can view the sample, cluster and final identity annotations on the peak based UMAP, together with per-cell accessibility (log2(RPKM+1)) for each 20 ChromHMM states. Cluster-level RPKM normalized scATAC-seq coverage tracks are displayed in an embedded IGV genome browser (igvShiny v1.9.51) on the mm10 genome, where users can search any gene or locus and select which cluster tracks to display on the shared scale.

### Materials availability

Plasmids generated in this study are available from the lead contact on request. The inducible ESC lines generated in this study are available from the lead contact under a completed materials transfer agreement.

## Author Contributions

Conceptualization: K.J., S.P and C.C. Data curation: H.F., A.P., I.S., E.H.W.L., S.Z., M.M. and C.C. Formal analysis: H.F., K.J., A.P., I.S., E.H.W.L., S.Z., M.M., A.F., L.D., S.P. and C.C. Funding acquisition: C.C. and S.P. Investigation: K.J., I.S., E.H.W.L., S.Z. and C.C. Methodology: H.F., K.J., A.P. M.M and C.C. Project administration: C.C. Resources: N.H, S.P. and C.C. Software: H.F. and A.P. Supervision: S.P. and C.C. Validation: H.F., A.P., I.S., E.H.W.L., S.Z., M.M. and C.C. Visualization: H.F., A.P., I.S., E.H.W.L., S.Z., M.M. and C.C. Writing – original draft: C.C. Writing – review & editing: H.F., K.J., A.P., I.S., S.P. and C.C.

## Acknowledgements

We thank I. Dror for discussions on computational approaches to define motif usage, and members of the Chronis and Pinho laboratories for advice and critical reading of the manuscript. We thank A. Kenter for providing pro-B cell culture reagents and protocols and J. Liang, W. Jiang and C. Simpson for the INCITE allocations that supported the computational work. The single-cell multiome assays benefited from consultation with the UIC Spatial and Genome Technologies Core, supported by the UIC Research Resources Center and the UIC Cancer Center Shared Resources, and we thank Melissa Leone for her guidance. This work was supported by National Institutes of Health grants R01HL170286 and P01HL160469 to C.C., R01HL162584 and R21CA306039 to S.P. I.S. was supported by the Chicago KUH FORWARD training program (NIDDK TL1DK132769). A.F was supported by the Ruth L. Kirschstein National Research Service Award from the NIH National Heart, Lung, and Blood Institute (F30HL176078). S.Z was supported by the Glenn Foundation for Medical Research Postdoctoral Fellowship Continuation Awards. Awards of computer time (2023, 2025) were provided by the U.S. Department of Energy’s (DOE) Innovative and Novel Computational Impact on Theory and Experiment (INCITE) Program. This research used resources from the Argonne Leadership Computing Facility, a U.S. DOE Office of Science user facility at Argonne National Laboratory, which is supported by the Office of Science of the U.S. DOE under Contract No. DE-AC02-06CH11357.

---

**Fig. S1.**
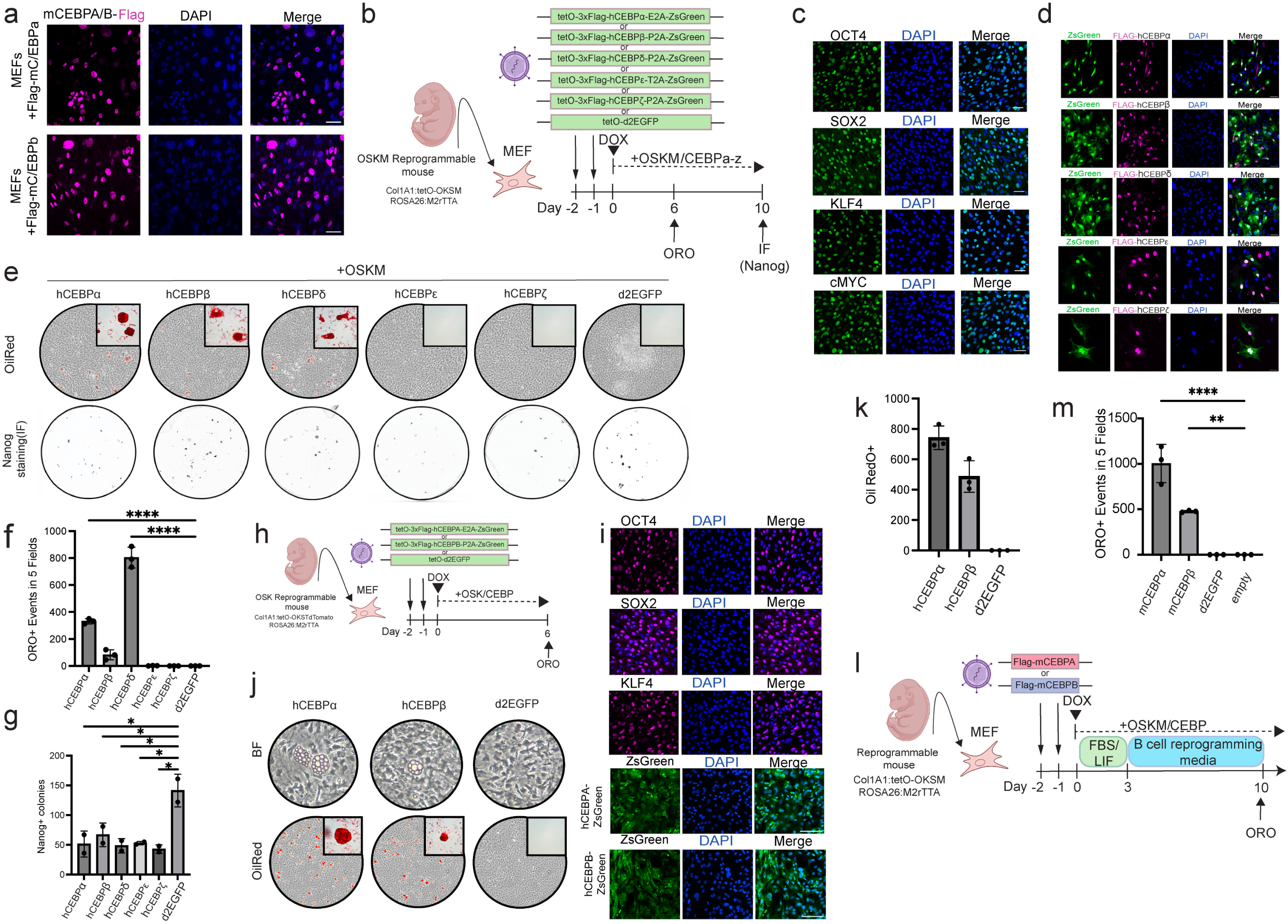
Multiple Cebp family members drive adipogenic conversion independently of species homolog, factor order and reprogramming conditions. **a**, Immunofluorescence of MEFs retrovirally transduced with Flag-tagged mCebpα (top) or mCebpβ (bottom), stained for Flag (magenta) and DAPI (blue); merge at right. **b**, Schematic of the inducible human Cebp constructs (tetO-3×Flag-hCebpα, β, δ, ε or ζ fused to ZsGreen) and experimental design. Reprogrammable MEFs lentivirally transduced and cultured with dox to co-induce OSKM and hCebp; ORO at day 6, Nanog immunofluorescence at day 10. **c**, Immunostaining for Oct4, Sox2, Klf4 and cMyc (green) and DAPI (blue) 48 h after dox induction of the polycistronic OSKM cassette in MEFs. **d**, Immunostaining for Flag (magenta) and DAPI (blue) in MEFs transduced with ZsGreen-fused tetO-3×Flag constructs encoding hCebpα, hCebpβ, hCebpδ, hCebpε or hCebpζ (green), confirming expression and nuclear localization of each factor in transduced (ZsGreen+) cells. **e**, Representative wells showing ORO staining (top, day 6) and Nanog immunostaining (bottom, day 10) across all hCebp family members and the EGFP control co-expressed with OSKM; ORO-positive lipid accumulation occurs selectively in the hCebpα, hCebpβ and hCebpδ conditions. **f**, Quantification of ORO-positive events across five microscopic fields at day 6 for each hCebp family member and the EGFP control co-expressed with OSKM (mean ± s.d.; n = 3). **g**, Quantification of Nanog-positive iPSC colonies at day 10 across hCebp family members and the EGFP control co-expressed with OSKM (mean ± s.d.; n = 2). \*\**P* < 0.01, \**P* < 0.05, unpaired two-tailed Student’s t-test. **h**, Reprogrammable MEFs (Col1a1:tetO-OSKtdTomato; ROSA26:M2rtTA) transduced with inducible hCebpα or hCebpβ and cultured with dox to co-induce OSK; ORO at day 6. **i,** Immunostaining for Oct4, Sox2, Klf4 (magenta), hCebpα or hCebpβ (ZsGreen) and DAPI (blue) 48 h after OSK induction. **j**, Representative wells showing brightfield (top) and ORO staining (bottom) at day 6. **k**, Quantification of ORO-positive events across five microscopic fields at day 6 for hCebpα, hCebpβ condition and the EGFP control co-expressed with OSK (mean ± s.d.; n = 3). **l**, Schematic of the reprogramming assay comparing mCebpα and mCebpβ activity under B-cell reprogramming conditions. Reprogrammable MEFs (OSKM) transduced with Flag-mCebpα or Flag-mCebpβ retroviruses were cultured in FBS/LIF from day −1 to day 3 and then in B-cell reprogramming medium (N2/B27/2i) with concurrent OSKM induction; ORO staining was performed at day 10. **m**, Quantification of ORO-positive events across five microscopic fields at day 10 for the Flag-mCebpα, Flag-mCebpβ, GFP and empty-vector conditions (mean ± s.d.; n = 3). ****denotes *P* < 0.001, ** denotes *P* < 0.01, unpaired two-tailed Student’s t-test.

**Fig. S2.**
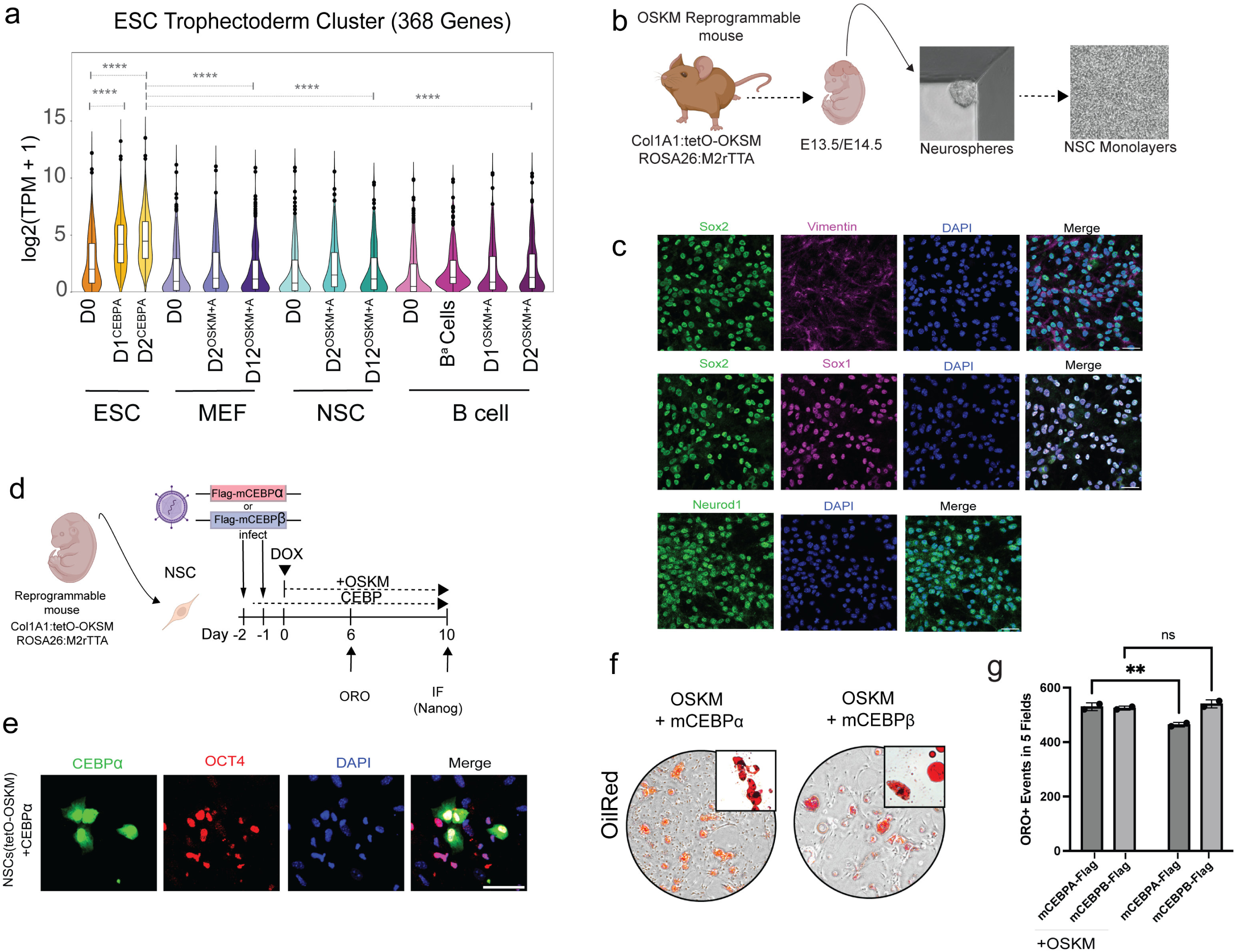
Cebp redirects OSKM-reprogramming NSCs toward adipocytes but induces a non-adipogenic, trophectoderm signature in ESCs. **a**, Expression (log_2_(TPM + 1)) of the ESC trophectoderm cluster (cluster 5; Fig. 2e) across the reprogramming time courses: ESCs ± hCEBPα (Day1, Day2), MEF, MEF-derived Day2^OSKM+A^ and MEF-derived D12^OSKM+A^, NSC, NSC-derived Day2^OSKM+A^ and NSC-derived D12^OSKM+A^, primary B cells, and B cells expressing CEBPα alone (B^α^) or with OSKM for 1 (D1^OSKM+A^) or 2 days (D2^OSKM+A^). The cluster is selectively upregulated in Cebpα-induced ESCs only. Box, median, and interquartile range (IQR); dots, outliers. \*\*\*\**P* < 0.0001. **b**, NSC derivation scheme. Neurospheres were isolated from the brains of E13.5/E14.5 embryos of reprogrammable mice (Col1a1:tetO-OSKM; ROSA26:M2rtTA), and NSC monolayers were established on poly-D-lysine/laminin-coated surfaces. **c**, Multicolor immunostaining of NSC monolayers for the neural stem/progenitor markers Sox2 (green) with Vimentin (magenta) or Sox1 (magenta), and for Neurod1 (green), each counterstained with DAPI (blue); merges at right. **d**, Schematic of concurrent mCebp overexpression and OSKM-mediated reprogramming. Reprogrammable NSCs (Col1a1:tetO-OSKM; ROSA26:M2rtTA) were transduced with Flag–mCebpα or Flag–mCebpβ and then induced with dox; cultures were assayed by ORO at day 6 and Nanog IF at day 10. **e**, Immunofluorescence of reprogrammable NSCs (tetO-OSKM) transduced with Flag-hCebpα-ZsGreen, stained for Cebpα (green), Oct4 (red) and DAPI (blue) at day 2 post-induction; merge at right. **f**, Representative brightfield/ORO-stained (red) images at day 6 of NSCs co-expressing OSKM with mCebpα or mCebpβ; insets show high-magnification ORO-positive lipid droplets. Cebp-expressing cultures display adipocyte-like morphology with lipid accumulation. **g**, Quantification of ORO-positive events across five microscopic fields per experiment (mean ± s.d.; n = 2) in NSCs expressing Flag-mCebpα or Flag-mCebpβ together with OSKM (+OSKM; left two bars) or alone (right two bars). ** denotes *P* < 0.01 (mCebpα + OSKM vs mCebpα alone); ns, not significant (mCebpβ + OSKM vs mCebpβ alone); unpaired two-tailed Student’s *t*-test.

**Fig. S3.**
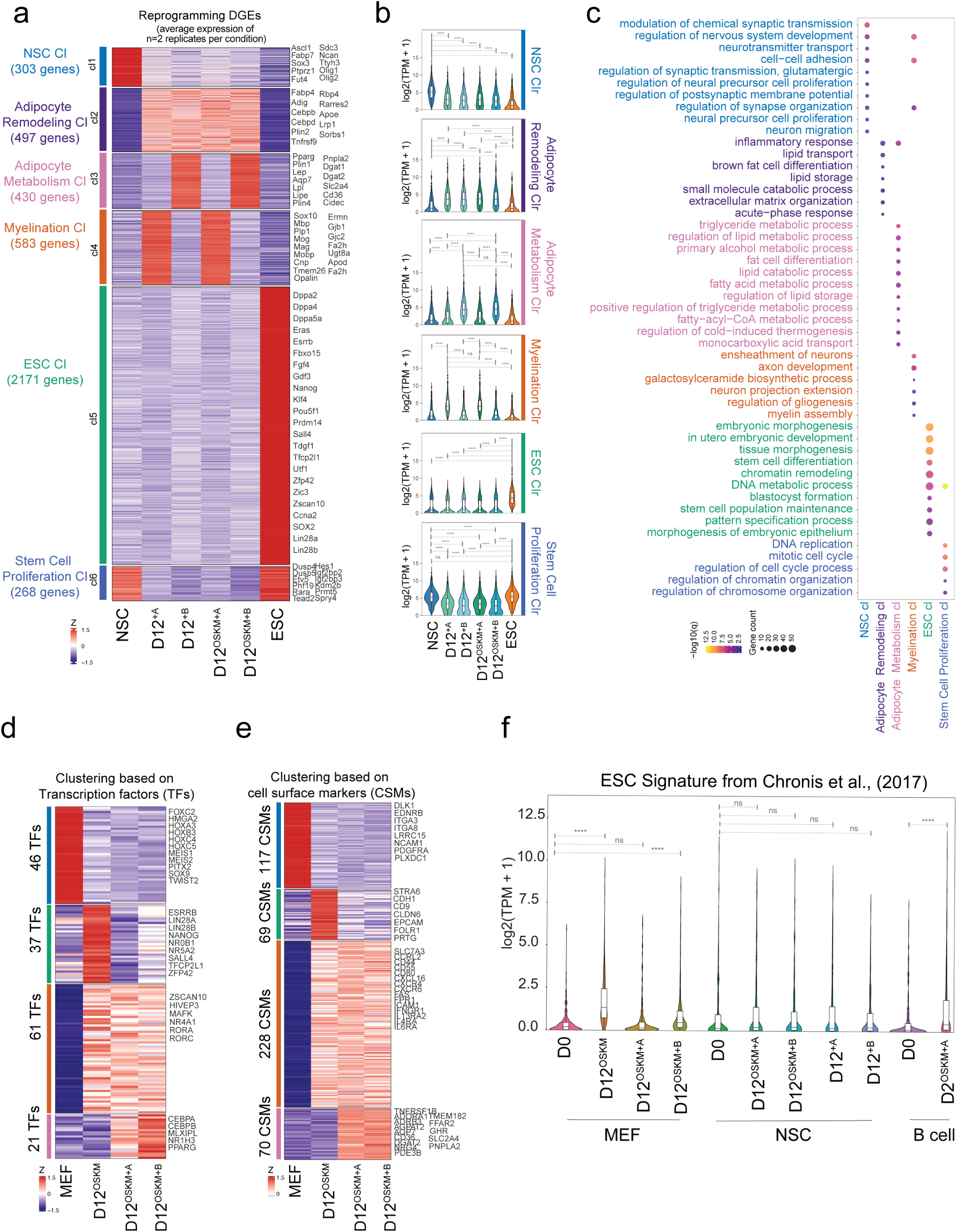
Cebpα/β rewires the NSC and MEF transcriptomes toward an adipogenic program at the expense of the OSKM-driven pluripotency signature. **a**, k-means clustering of differentially expressed genes across NSC; NSCs expressing Cebpα or Cebpβ for 12 days, alone or with OSKM; and ESCs (Z-scored), resolving six clusters (naming by ontology): NSC (303 genes), Adipocyte Remodeling (497), Adipocyte Metabolism (430), Myelination (583), ESC (2,171) and Stem-Cell Proliferation (268); representative genes at right. **b**, Violin plots of aggregate log_2_(TPM + 1) expression of each cluster across the samples in a. Asterisks mark significant differences between conditions (Benjamini–Hochberg correction for multiple comparisons); ns, not significant. **c**, GO biological-process enrichment of the clusters in a; dot size indicates gene count and color indicates −log_10_(*P*). **d**, k-means-clustered heatmap of transcription-factor (TF) expression across MEF, OSKM, OSKM + hCebpα and OSKM + hCebpβ at day 12 post-induction (Z-score-normalized), resolving four clusters (46, 37, 61 and 21 TFs). The 21-TF cluster, selectively enriched in OSKM + hCebpα and OSKM + hCebpβ at day 12, comprises core adipogenic regulators including *Cebpa*, *Cebpb*, *Nr1h3*, *Mlxipl* and *Pparg*; representative TFs annotated at right. **e**, k-means-clustered heatmap of cell-surface-marker (CSM) expression across the samples in d, resolving four clusters (117, 69, 228 and 70 CSMs); representative markers annotated at right. **f**, Expression (log_2_(TPM + 1)) of the Chronis et al. (2017; GSE90895) ESC signature across MEF (D0, D12^OSKM^, D12^OSKM+A^ and D12^OSKM+B^), NSC (D0, D12^OSKM+A^ and D12^OSKM+B^, D12^Cebpa^, D12^Cebpβ^), primary B cell, and B cells expressing OSKM for 48h following a 12h pulse with Cebpα. \*\*\*\**P* < 0.0001; ns, not significant; Benjamini–Hochberg correction for multiple comparisons.

**Fig. S4.**
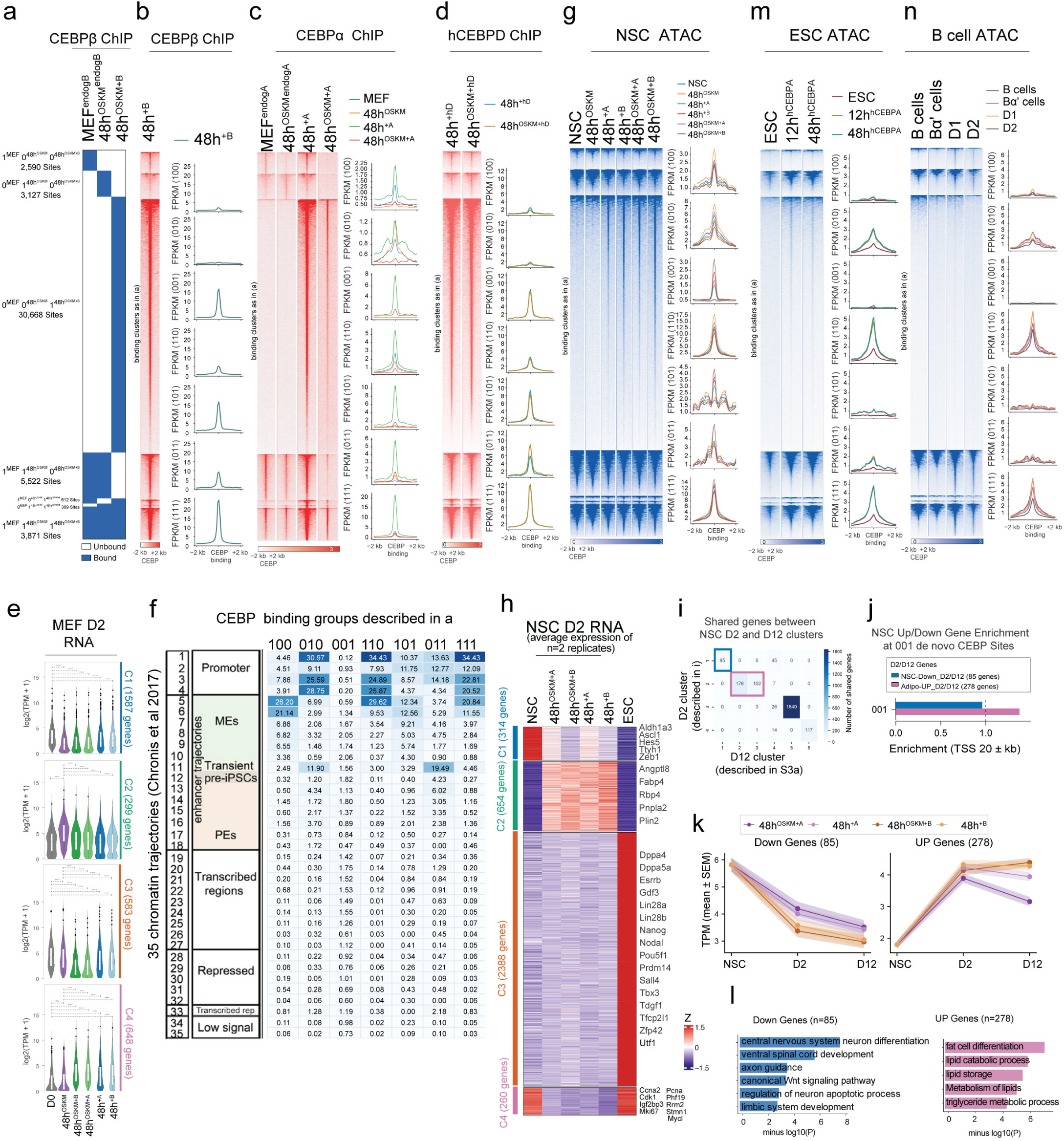
Conserved CEBP occupancy in MEFs and NSCs directly engages and activates the adipogenic gene regulatory network. **a,** Cebpβ ChIP-seq at peaks called at 48 h of OSKM or OSKM+Cebpβ expression, with endogenous MEF binding; peaks grouped by presence or absence of calls across MEF (endogenous), 48h^OSKM^ (endogenous) and 48h^OSKM+B^ (over-expressed) MEFs into seven occupancy clusters; digits, binding in MEF, 48h^OSKM^ and 48h^OSKM+B^ (bound, blue; unbound, white). Site counts left; centered on CEBP summits (±2 kb). **b**, Cebpβ ChIP-seq at MEFs retrovirally infected with Cebpβ overexpression construct at the sites defined in a, ordered by cluster; centered on CEBP summits (±2 kb). **c**, Cebpα ChIP-seq at the sites in a: MEF (endogenous), 48h^OSKM^ (endogenous), 48h^+CEBPA^, 48h^OSKM+A^; ordered as in a (±2 kb). Metaplots (FPKM): MEF, blue; 48h^OSKM^, orange; 48h^+CEBPA^, green; 48h^OSKM+A^, red. **d**, FLAG ChIP-seq signal of over-expressed FLAG-tagged human Cebpδ (hCebpδ) ± OSKM at the sites in a (±2 kb). **e**, Violin plots of the gene clusters in Fig. 4d across MEF and MEFs induced 48 h with OSKM, Cebpα, Cebpβ, OSKM + Cebpα or OSKM + Cebpβ. **f**, Enrichment of the seven CEBP binding groups (100–111; columns, as defined in a & Fig. 4a) across the 35 chromatin trajectories from Chronis et al. 2017 (rows; promoter, MEs, transient pre-iPSC, PEs, transcribed regions, repressed, transcribed-repressed, low-signal). Color and value, enrichment. **g**, ATAC-seq of NSC samples at sites defined in a: NSC, NSCs induced with OSKM, Cebpα alone, Cebpβ alone, OSKM+A, OSKM+B for 48 hrs, ordered as in a (±2 kb). Metaplots (FPKM) coloured by condition as in c. **h**, *k*-means clustering of differentially expressed genes (row Z-score) across NSC, NSCs induced for 48 h with OSKM + Cebpα, OSKM + Cebpβ, Cebpα or Cebpβ, and ESC (mean of *n* = 2 replicates), resolving four clusters: C1–C4; representative genes at right. **i**, Overlap between the NSC day-2 gene clusters (C1– C4 from h; rows) and the NSC day-12 clusters (1–6 from S3a; columns). Cell color and number indicate the number of shared genes. **j**, Enrichment of the Cebpβ occupancy clusters from a within ±20 kb of TSSs, for genes commonly downregulated at days 2 and 12 (NSC-Down_D2/D12; *n* = 85) or commonly upregulated by day 2 within the day-12 adipocyte cluster (Adipo-UP_D2/D12; *n* = 278). Dashed line, enrichment = 1. **k**, Expression trajectories (TPM; mean ± s.e.m.) across NSC, D2, D12 for Cebpα (blue) and Cebpβ (red): downregulated genes (n = 85; left), upregulated genes (n = 278; right). **l**, GO biological-process enrichment for downregulated (*n* = 85, left) and upregulated (*n* = 278, right) gene sets in k;−log₁₀(*P*). **m**, ESC ATAC-seq at the sites in a for samples ESC, ESCs induced to overexpress hCEBPA for 12 or 48h; ordered as in a (±2 kb). **n**, B-cell ATAC-seq at the sites defined in a for samples: B cells, B cells transiently induced with Cebpα, and additionally for 1 or 2 days with OSKM; ordered as in a (±2 kb).

**Fig. S5.**
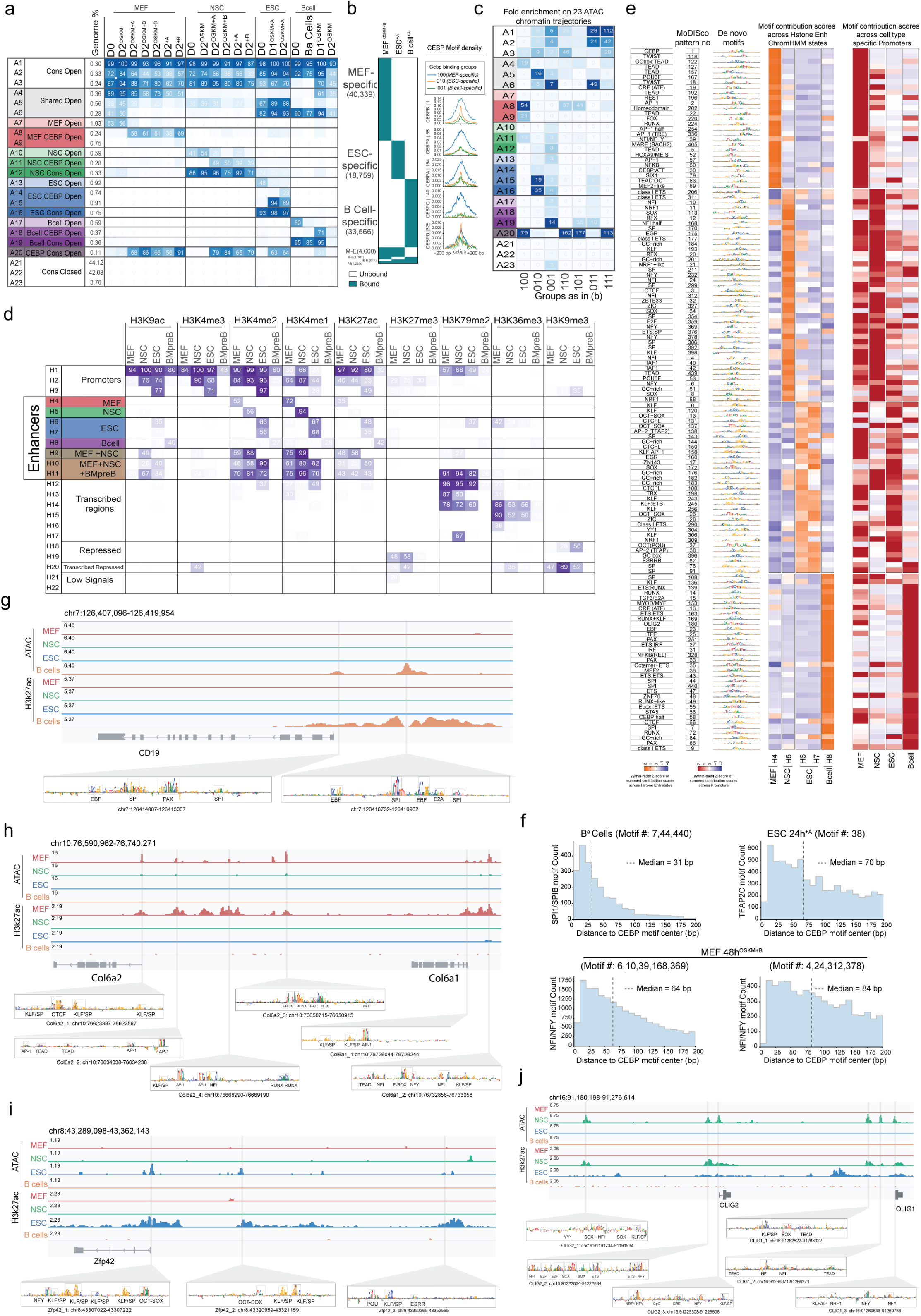
Chromatin trajectories, Cebp binding, and motif grammar underlying cell-type-specific accessibility remodeling. **a**, The 23 ATAC chromatin trajectories (A1-A23): number, annotation and genome fraction, then accessible-region frequency per condition (basal and Cebp-and/or OSKM-induced), 0 (white) to 100 (blue). **b**, Cell-type-specific Cebp binding groups and their Cebp motif content. Left, Cebpα/β ChIP-seq occupancy (bound, teal; unbound, white) at Cebp peaks in OSKM + Cebpβ MEFs, Cebpα-ESCs and Cebpα-B cells (GSE284146; GSE71218), defining MEF-specific (100; 40,339), ESC-specific (010; 18,759), B-cell-specific (001; 33,566) and shared groups (M–E, 4,660; M–B, 1,101; E–B, 911; all three, 1,235). Right, density of the de novo Cebp CWMs (CEBP_1, CEBP_58, CEBP_154, CEBP_140, CEBP_325) centered on Cebp peaks (±200 bp) for the MEF (100), ESC (010), and B-cell (001) groups. **c**, Fold enrichment of the binding groups in b across the 23 ATAC chromatin trajectories from a. **d**, The 22 ChromHMM chromatin states (H1–H22): number, annotation and genome fraction, then per-cell-type emission frequency by mark (H3K9ac, H3K4me3/2/1, H3K27ac, H3K27me3, H3K79me2, H3K36me3, H3K9me3; MEF, NSC, ESC, pre-B). **e**, TF-MoDISco de novo motif repertoire. Left to right: putative associated TF and MoDISco pattern number; de novo motif as a CWM; within-motif Z-score of summed per-base contribution scores across the five enhancer ChromHMM states (H4–H8); and within-motif Z-score of summed contribution scores across cell-type-specific promoters (MEF, NSC, ESC, B cell). **f**, Spacing between Cebp and partner motifs. Histograms of the distance from each partner-motif instance to the nearest Cebp motif center (±200 bp): SPI1/SPIB (Bα cells; patterns 7, 44, 440; median 31 bp); TFAP2C (Cebpα-ESC; pattern 38; median 70 bp); and two NFI/NFY motif sets (OSKM + Cebpβ MEF)-patterns 6, 10, 39, 168, 369 (median 64 bp) and patterns 4, 24, 312, 378 (median 84 bp). **g**, The *Cd19* locus, a B-cell-specific gene, showing ATAC and H3K27ac profiles across MEF, NSC, ESC and B cells, with per-base contribution scores and annotated motif instances (EBF1, SPI, PAX, E2A). **h**, As in g for the *Col6a* loci (MEF-associated), with annotated KLF, CTCF, SP, TWIST, RUNX, NFI, AP-1, TEAD, NFY and EGR motif instances. **i**, As in g for the *Zfp42* locus (gaining accessibility selectively in ESCs), with annotated OCT/SOX, ESRR, KLF/SP and NFI/NFY motif instances. **j**, As in g for the *Olig1*/*Olig2* loci (NSC-associated), with annotated SOX, NFI, TEAD, KLF, NFY, YY1 and TAF1 motif instances.

**Fig. S6.**
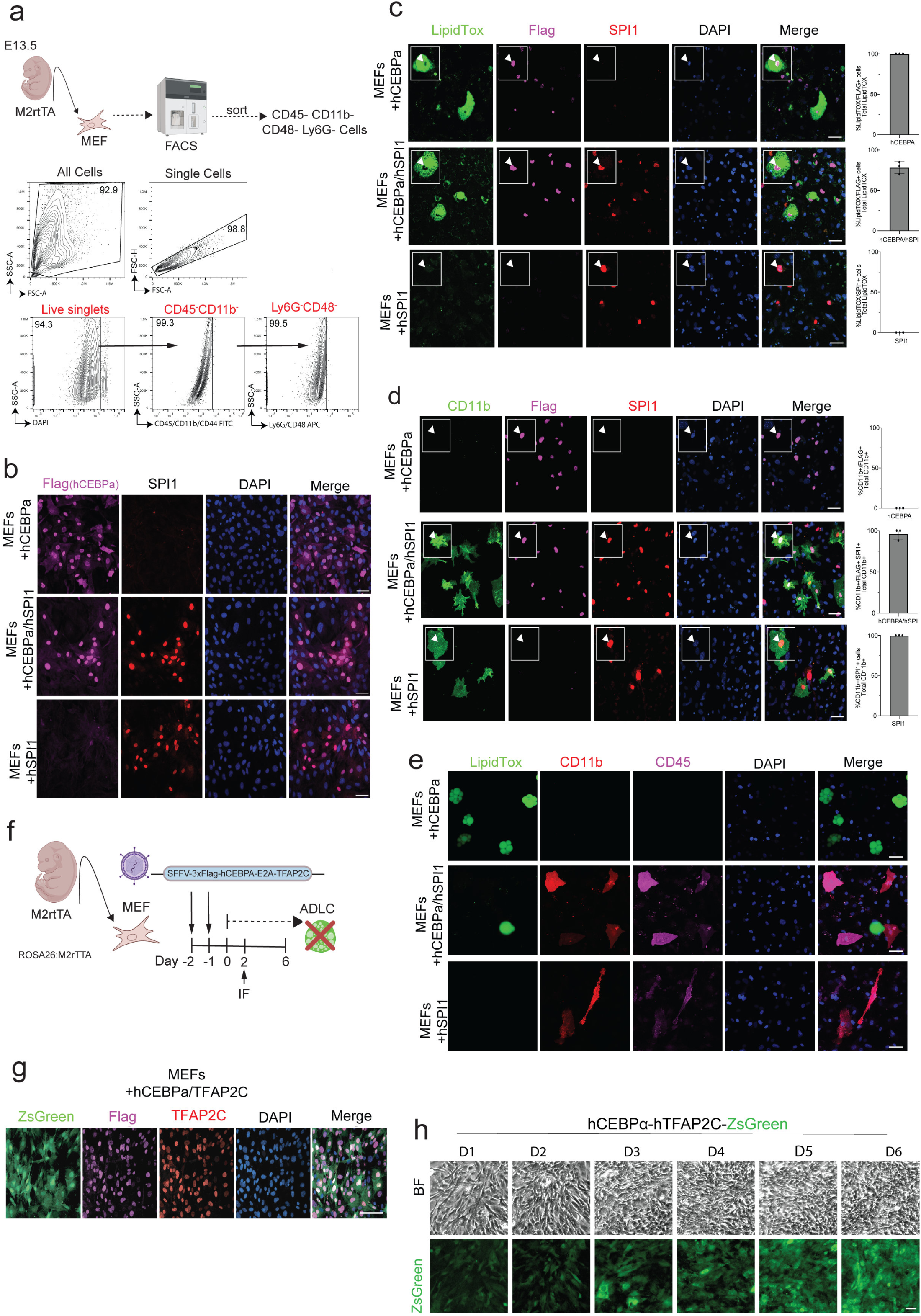
Reprogramming outcome is set by the active factor combination, not factor silencing. **a**, Negative-selection sort of the starting MEF population. MEFs from E13.5 embryos FACS-sorted for absence of hematopoietic/myeloid markers: cells (SSC-A/FSC-A), single cells (FSC-H/FSC-A), live (DAPI^−^) singlets, then CD45^−^CD11b^−^CD44^−^ and Ly6G^−^CD48^−^; numbers, % of parent gate. Input population free of pre-existing hematopoietic cells. **b**, Immunofluorescence at day 2 for Flag (3×Flag-hCebpα; magenta), SPI1 (red) and DAPI (blue) in hCebpα, hCebpα/hSpi1 and hSpi1; expected per-construct expression (Flag; Flag and SPI1; SPI1). **c**, Immunofluorescence at day 10 for LipidTOX (green), Flag (magenta), SPI1 (red) and DAPI (blue); arrowheads and insets, LipidTOX-positive adipocyte-like cells co-expressing Flag (hCebpα) in hCebpα and hCebpα/hSpi1. Right, LipidTOX^+^Flag^+^ cells as a fraction of total LipidTOX^+^ (mean ± s.d.; n = 3). **d**, As in c, stained for CD11b (Mac-1; green); CD11b-positive macrophage-like cells in hCebpα/hSpi1 and hSpi1 co-express Flag and/or SPI1. Right, CD11b^+^ cells co-expressing factor(s) as a fraction of total CD11b^+^ — Flag in hCebpα, Flag and SPI1 in hCebpα/hSpi1, SPI1 in hSpi1 (mean ± s.d.; n = 3). With c, both fates arise in factor-expressing cells: fate choice is instructive, not factor silencing or selection. **e**, Immunofluorescence at day 10 for LipidTOX (green), CD11b (red), CD45 (magenta), and DAPI (blue) in hCebpα, hCebpα/hSpi1, and hSpi1; CD11b-positive macrophage-like cells co-express the pan-hematopoietic marker CD45, LipidTOX-positive adipocyte-like cells do not. **f**, Cebpα/TFAP2C assay. MEFs (ROSA26:M2rtTA) transduced with SFFV-3×Flag-hCebpα-E2A-hTfap2c-ZsGreen, assessed through day 6 for adipocyte-like cell (ADLC) emergence; none observed. **g**, Immunofluorescence at day 2 for ZsGreen (green), Flag (magenta), TFAP2C (red) and DAPI (blue) in hCebpα/hTfap2c; co-expression confirmed. **h**, Brightfield (top) and ZsGreen (bottom), days 1–6, hCebpα/hTfap2c-ZsGreen MEFs; fibroblast morphology retained, no adipocyte-like conversion.

**Fig. S7.**
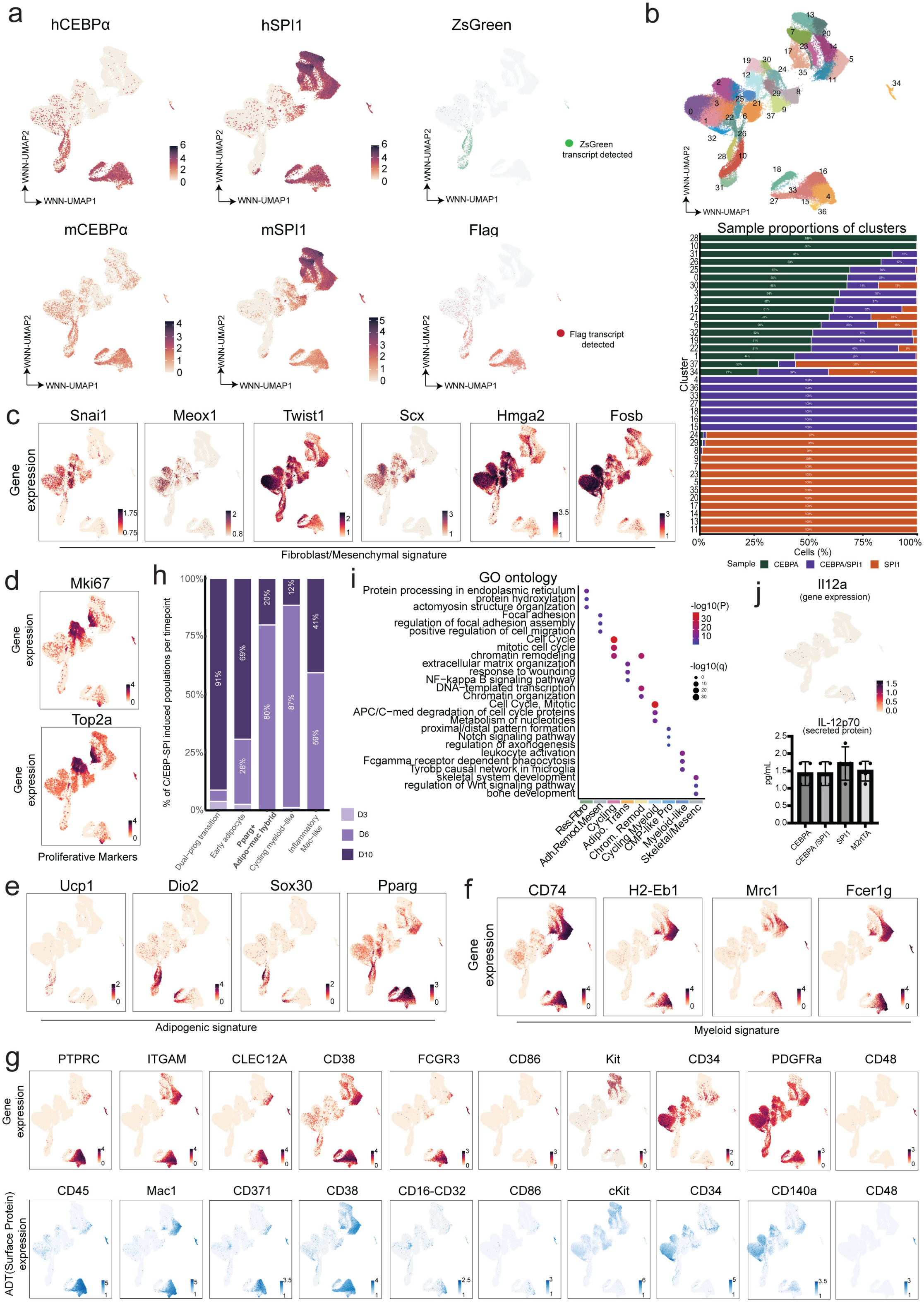
Cluster composition, lineage markers, and surface-protein phenotypes assign each CITE-seq state to a reprogramming arm and timepoint. **a**, Transgene-derived (*hCEBPα*, *hSPI1*) and endogenous (*mCebpa*, *mSpi1*) transcripts across the combined atlas (n = 110,751 cells); colour, log-normalised expression. Right of each row, ZsGreen (top) and Flag (bottom) reporter transcripts; cells with ≥ 1 transcript are colored. **b**, Top, WNN-UMAP colored by unsupervised cluster annotation (0–37). Bottom, per-cluster proportion of cells from MEF^CEBPα^, MEF^CEBPα/SPI1^,and MEF^SPI1^, ordered by CEBPα contribution. **c**, Fibroblast/mesenchymal markers *Snai1*, *Meox1*, *Twist1*, *Scx*, *Hmga2* and the AP-1 family member *Fosb* across the combined atlas; colour, log-normalised gene expression. **d**, Proliferation markers *mKi67* and *Top2a* as in c. **e**, Adipogenic markers *Ucp1*, *Dio2, Sox30, and Pparg* as in c. **f**, Myeloid markers *Cd74*, *H2-Eb1, Mrc1 and Fcer1g* as in c. **g**, Matched transcript (top) and surface-protein (ADT; bottom) measurements for ten marker pairs: *Ptprc*/CD45, *Itgam*/Mac-1, *Clec12a*/CD371, *Cd38*/CD38, *Fcgr3*/CD16–CD32, *Cd86*/CD86, *Kit*/cKit, *Cd34*/CD34, *Pdgfra*/CD140a, *Cd48*/CD48. RNA, log-normalized counts; ADT, centered log-ratio-normalized counts. **h**, Timepoint composition (day3, day6, day10 after dox induction) of five MEF^CEBPα/SPI1^ induced populations. The Pparg+ adipo-mac hybrid is maximal at day 6, whereas the early adipocyte and inflammatory Mac-like states skew to day 10. **i**, GO and pathway enrichment for the remaining nine states not shown in Fig. 7f; dot size, −log∼10∼(*q*); color, −log∼10∼(*P*). **j**,Top *Il12a* expression across the combined atlas. Bottom, IL-12p70 protein in day-10 culture supernatants of MEF^CEBPα^, MEF^CEBPα/SPI1^,MEF^SPI1^and dox-only (M2rtTA) cultures (mean ± s.d.; n = 4);secretion does not rise above the M2rtTA control.

**Fig. S8.**
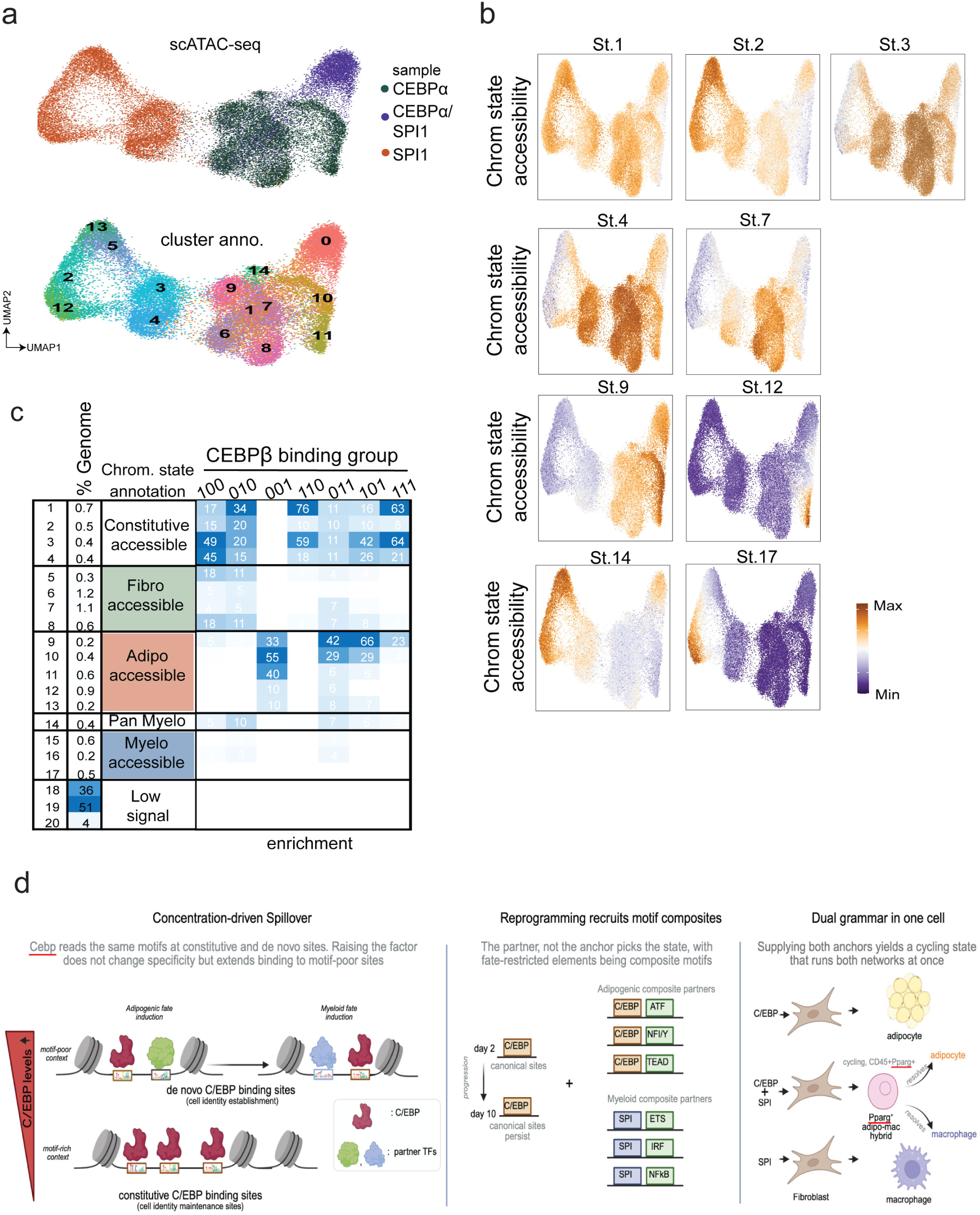
De novo Cebpβ occupancy is confined to the adipogenic chromatin states and supports a concentration-and partner-dependent model of Cebp-directed fate choice. **a**, scATAC-seq UMAP (embedding as in Fig. 8a) colored by sample (top; hCebpα, hCebpα/hSPI1, hSPI1) and by unsupervised cluster (bottom; cl0–cl14). **b**, Accessibility of the remaining chromatin states projected on the scATAC UMAP, as in Fig. 8c: fibroblast-and constitutive-accessible states 1, 2, 3, 4 and 7; adipogenic-accessible states 9 and 12; pan-myeloid state 14; and SPI myeloid-accessible state 17. Color, log_2_(RPKM+1). **c**, Enrichment of each of the 20 chromatin states (Fig. 8b) within the seven Cebpβ occupancy groups defined in Fig. 4a; digits denote binding in MEF, 48 h OSKM and 48 h OSKM + Cebpβ, respectively. **d**, Model of Cebp-directed fate selection. Left, concentration-driven spillover: Cebp reads the same elements at constitutive and de novo sites, so raising factor levels extends binding from motif-rich maintenance sites into motif-poor contexts rather than altering specificity; the partners redeployed at neighboring sites dictate whether the de novo site drives an adipogenic or myeloid program. Middle, composite recruitment during reprogramming: canonical C/EBP sites engaged at day 2 persist to day 10 alongside fate-restricted composite elements — C/EBP:ATF, C/EBP:NFI/NF-Y and C/EBP:TEAD in the adipogenic arm; SPI:ETS, SPI:IRF and SPI:NF-κB in the myeloid arm. Right, dual grammar in one cell: Cebpα alone converts fibroblasts to adipocyte-like and SPI1 alone to macrophage-like cells, whereas supplying both anchors yields a cycling CD45⁺Pparg⁺ adipo–mac hybrid that runs both networks and subsequently resolves to either fate. Created with BioRender.com under agreement number WE2AAMTSI5.

## Notes

https://cebpcompass.org

### Competing Interest Statement

The authors have declared no competing interest.

